# Hazard Structure and Repair Kinetics: A 2×2 Classification that Resolves the Stochastic-Death / Individual-Tolerance Dilemma

**DOI:** 10.64898/2026.11.03.764101

**Authors:** Jong-Hyeon Lee, Chan Young Joe

## Abstract

The two variants of GUTS (General Unified Threshold model of Survival), stochastic death and individual tolerance, disagree about no-effect concentrations under time-variable exposure, and no standard test separates them. Each variant fixes two independent choices, which form a 2×2 classification: whether the hazard accumulates while damage exceeds a threshold (cumulative) or is set by the highest damage reached (peak-based), and whether repair is proportional or saturating. Calibration fit cannot choose: each structure fits by adjusting its own damage recovery, the cumulative one faster in twelve of fourteen compounds. For two identical pulses separated by complete recovery, the cumulative hazard predicts equal survival through both whatever the parameters, so a carry-over ratio (second-to-first survival) above one excludes it; in *Gammarus pulex* fourteen of fifteen ratios are at or above one, seven significantly. The peak-based hazard predicts the ratio’s maximum, approached as intervals lengthen in five of six compounds. For organophosphates, whose recovery outlasts the published pulse intervals, the one test on measured damage favours the peak-based hazard: diazinon-exposed *Daphnia magna* stopped being immobilised once acetylcholinesterase, still inhibited, began to recover. The peak-based hazard is the appropriate default for toxicokinetic–toxicodynamic risk assessment of narcotics and uncouplers, and provisionally of organophosphates.

## 1. Introduction

When a pesticide enters a stream after a rain event, aquatic organisms experience a pulse of toxicant that rises, peaks and then declines as the chemical is diluted or degraded. Predicting how many survive such events, and at what concentrations protection fails, is the central task of aquatic risk assessment. The General Unified Threshold model of Survival (GUTS) is the standard toxicokinetic–toxicodynamic (TKTD) framework for this task, adopted by EFSA, implemented in openGUTS and applied to a widening range of compounds and species.^1–4^

GUTS offers two reduced (damage inferred, not measured) variants: GUTS-RED-SD (stochastic death) and GUTS-RED-IT (individual tolerance). In SD, organisms exposed above a shared damage threshold die at a rate proportional to the current excess, so the hazard (the death rate) keeps accumulating for as long as damage stays above the threshold. In IT, each organism carries its own threshold and dies as soon as damage first crosses that threshold.

This difference matters for regulation in two ways. Under prolonged exposure the SD hazard accumulates without limit, so any concentration above the shared threshold is eventually lethal and the LC50 converges on that threshold, whereas IT predicts mortality levelling off, so the LC50 settles at the median tolerance (§S1.1–S1.2).^2^ Under time-variable exposure, fitted to the same data, they give different species sensitivity distributions and rankings of exposure profiles,^3^ and which is the more conservative changes with exposure duration and is undetermined at the durations validation tests use (§4.2). Yet in standard constant-exposure tests both fit the data equally well — the “death dilemma” (§4.3).^2^

An experiment designed to separate them has been tried. Sowa *et al.*^4^ dosed *Poecilus cupreus* repeatedly with recovery between exposures and concluded in favour of individual tolerance. Their own AIC comparison favoured stochastic death in both beetle populations tested; for the meadow population they chose individual tolerance on other goodness-of-fit measures. The model comparison and the conclusion pointed in opposite directions: the fit did not decide the question.^5,6^ Nor is the failure of fit to decide particular to that design. Across more than a hundred published datasets neither variant fits consistently better,^3,5^ and the reason is structural: a killing rate and a spread of thresholds produce the same survival curves, so a model carrying both is not identifiable from survival data (§S9.4).

What is missing, then, is evidence that does not rest on which model fits better. Obtaining it requires first taking the dilemma apart, because each variant fixes two logically independent choices at once.

The first: does death depend on the **total accumulated** damage or on the **highest level** damage ever reaches? The second: does repair scale proportionally with damage, or saturate at a maximum rate? The two are independent: the first — the hazard structure, cumulative or peak-based — decides which feature of the damage trajectory the hazard depends on, the second only how that trajectory rises and falls. Separating them produces a 2×2 classification in which SD (cumulative) and IT (peak-based) each occupy one cell (§2.2); both assume proportional repair, so within the classification they differ only in the hazard structure. What a hazard structure can produce at any parameter value is settled by a theorem; the rate at which repair runs is settled by fitting, once the structure is fixed (§5.3). The dilemma therefore turns on the hazard structure — the choice a theorem can settle without a fit.

This paper develops the classification and its two theorems (§2.3). Both concern one measured quantity, the carry-over ratio: survival through a second pulse, among survivors of the first, relative to survival through the first. A ratio above unity excludes the cumulative structure, and a ratio approaching the maximum the peak-based structure predicts confirms it. §4 tests the theorems in published counts (§4.1), asks which structure is the more conservative at matched parameters (§4.2), shows by simulation that calibration fit cannot identify the hazard structure (§4.3), and shows on two organophosphate datasets that fit comparison does not settle the structure once the data and toxicokinetics are those published (§4.4); §5.3 shows why fit cannot decide and turns to the one organophosphate test on measured damage. A companion manuscript, submitted alongside,^7^ takes the classification as given and shows that the observations which separate SD from IT in calibration reflect the damage model — the lesion’s chemistry and the test’s length — rather than the hazard structure. That finding is one reason the recommendation of §5.5 rests on structural results rather than on calibration fit (§5.4).

## 2. Theory

### 2.1 Two independent modelling choices

Irreversible damage can be read in two ways, as a rain gauge and a flood marker read the same storm: the total that fell, or the highest level reached. The two hazard structures take one reading each: the damage accumulated over the whole exposure, or its highest level.

In GUTS-SD, the hazard grows with **both how long and how far** damage stays above a threshold — the “rain gauge” approach:

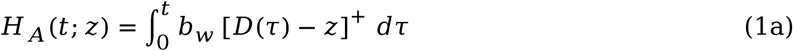

We call this Class A (cumulative hazard). In Class B (peak-based hazard), the hazard depends only on the **highest damage ever experienced** — the “flood marker”:

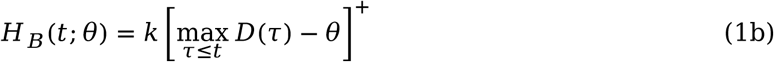

Here *D* is damage, *z* a threshold shared by all organisms, *θ* an individual’s own tolerance, and *b_w_* and *k* killing rates. Once damage exceeds *θ* the Class B hazard is set by the level reached; Class A instead keeps accumulating for as long as damage stays above *z*. Neither hazard ever decreases, so the irreversibility of the lesion does not discriminate between them. Acute aquatic toxicity is dominated by threshold transitions rather than accumulating processes: mitochondrial permeability transition,^8^ irreversible phosphorylation of acetylcholinesterase,^9^ oxidative collapse.^10^ The same distinction places the existing hazard models in one framework.^11^ Those mechanisms are consistent with the peak-based structure without discriminating by themselves (§S6.4). For Class B with a tolerance distribution (§2.4) we propose the term **GUTS-PM** (Peak Maximum), extending the GUTS naming convention (§5.5).

The second independent choice concerns the repair function *R*(*D*) in the damage equation *dD*/*dt* = *k_a_C_b_*(*t*) − *R*(*D*), with *C_b_* the internal concentration and *k_a_* the damage-formation rate. The standard GUTS assumption is proportional repair (*R* = *k_r_D*); the alternative is constant-rate repair (*R* = *k_r_*_,0_). A saturating form with repair capacity *K* contains both as limits: *K* → ∞ proportional, *K* → 0 constant-rate (§S5). Enzymatic repair — AChE re-synthesis, glutathione regeneration — runs on machinery of finite capacity and may saturate at high damage, although plasma cholinesterase in workers over-exposed to an organophosphate recovered exponentially (half-life about 12 days), the proportional form.^13^ The saturating form nonetheless leaves no signature the data analysed here can resolve, so everything below works in the proportional limit *K* → ∞ (§S5).

### 2.2 The 2×2 classification

Crossing the two hazard structures with the two repair types produces four survival behaviours (Table 1); the bottom row is each hazard with constant-rate repair, and each cell’s trajectories are in Figure S1 (§S1). Under the peak-based structure the threshold applies to a level of peak damage, so the survivors of a pulse are those whose threshold lies above its peak — a selected subset that a repeat of the same pulse cannot kill. Under the cumulative structure the threshold is shared and death is a chance event driven by accumulated hazard, so surviving a pulse selects nobody: once damage has reset, the survivors face the second pulse as the whole population faced the first. That is why the carry-over ratio of §2.3 is exactly one for the cumulative structure (Proposition S4, §S6.1.2).

**Table 1.** The 2×2 classification of TKTD survival models. *K* is the repair capacity (§2.1): *K* → ∞ proportional repair, *K* → 0 constant-rate repair. Haber’s law sits in the peak-based column because its constant post-threshold death rate follows from toxicokinetics alone (§S1.4).

|  | **Class A: Cumulative hazard** | **Class B: Peak-based hazard** |
|---|---|---|
| **Proportional repair**<br>(*K* → ∞) | **GUTS-SD:** hazard keeps accumulating, so all organisms eventually die | **GUTS-IT:** hazard stops at the peak, so tolerant organisms survive permanently |
| **Constant-rate repair**<br>(*K* → 0) | Death rate rises with time — *accelerating mortality* | Constant death rate after threshold — *Haber’s law*<sup>14</sup> |

### 2.3 Two theorems: exclusion and confirmation

Neither the irreversibility of the lesion (§2.1) nor goodness of fit decides between the two structures: within a standard test they produce nearly indistinguishable survival curves.^2^ The choice has to rest instead on what each structure can and cannot represent, and on whether the parameter values a fit requires are chemically plausible. Two theorems are available: the first says what Class A must predict, so a measurement can exclude it (§S6.1); the second says the most Class B can predict, so a measurement can confirm it (§S6.2).

For two identical pulses separated by recovery, write the carry-over ratio as ℛ = *S* /*S* : second-pulse survival among survivors of the first, over first-pulse survival. If damage resets between the pulses, each pulse is an independent trial under Class A (§2.2) and ℛ is exactly unity at every parameter value, provided the animals share those parameters — a theorem, not an assumption (Proposition S4, §S6.1.2; Figure S3b). A measured ratio above unity therefore rules out GUTS-SD. The inequality runs one way: residual damage can only lower the ratio, so a ratio above unity excludes Class A whatever the recovery interval, while one at or below unity is interpretable only if the interval let damage reset (§S6.1). A killing rate varying between individuals is not covered by the theorem; whether the data allow enough of that variation is an empirical question, taken up in §4.1. A ratio above unity also requires selection: survivors of the first pulse must be a subset selected for tolerance. The running maximum supplies that selection by itself; a hazard driven by the rate of damage formation, or by the damage level in identical animals, does not (§S6.6). Class B is itself refutable in the same experiment: the running maximum freezes once damage stops rising, so deaths after exposure ends refute it (§S3.2).^15^

Excluding GUTS-SD is not accepting Class B; the peak-based structure makes a prediction of its own. Once damage has reset, the survivors of the first pulse are those whose threshold the first peak did not reach, and an identical second peak cannot reach it either, so *S*_2|1_ = 1 and ℛ = 1/*S* exactly, whatever the tolerance distribution (Proposition S5, §S6.2.1). That ceiling is fixed by first-pulse survival, not by the model, so the prediction is an approach to it as the interval lengthens relative to the compound’s recovery time. Neither the cumulative structure nor a version of it with a killing rate varying between individuals predicts that approach.

### 2.4 Established models as limiting cases

The peak-based hazard (Class B) with a tolerance distribution:

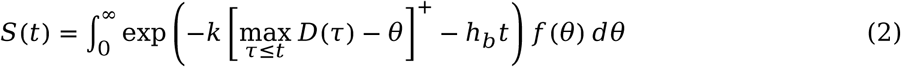

recovers GUTS-IT (*k* → ∞), the damage assessment model DAM^16^ (the case *θ* = 0, constant exposure) and the stress addition model SAM (both limits, with additive stressors) as exact limiting cases, while GUTS-SD occupies a different cell and is not recovered; the limits and their conditions are in §S2. Within Class B, SD-like (ratio at unity) and IT-like (ratio at the ceiling) behaviour are the ends of a continuum: the ratio rises through unity as damage recovery quickens relative to the interval, so *k_r_* places a compound on that continuum (§S6.2).

### 2.5 Why a missing activation stage favours the cumulative hazard

A comparison of hazard structures inside the reduced (RED) framework carries a bias for compounds that act through an activated metabolite. In a multi-stage activation pathway (parent → metabolite → phosphorylated acetylcholinesterase, AChE-P, for organophosphates), each stage adds a power of time to the early damage trajectory (*D*(*t*) ∝ *t^n^*^eff^, *n*_eff_ ≈ 2 for three stages; §S4). With the toxicokinetics of the two compounds bracketing dimethoate (§3.2) the exponent over the four-day test is 1.78–2.43, against 0.98 for the one-compartment framework (Figure S5a, §S8.2). A peak-based hazard driven by a superlinear trajectory produces accelerating mortality. A cumulative hazard driven by the linear trajectory that RED gives produces the same accelerating mortality, because integration adds a power of time, so the two combinations predict the same survival curve. In the reduced framework, which lacks that curvature, the cumulative structure’s integral can supply it, so a better reduced-framework fit for the cumulative structure is weak evidence about the hazard (§4.4). A multi-stage model also lets damage peak days after a brief pulse, fed by residual metabolite, which RED cannot represent (Figure S5b, §S8.2).

## 3. Materials and Methods

### 3.1 Model configurations

For the organophosphates (OPs) the three-compartment model represents the parent → metabolite → AChE-P cascade explicitly:^18^

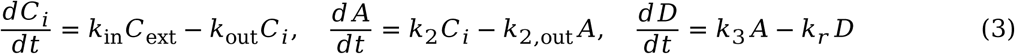

*C_i_* is the internal parent concentration, *A* the activated metabolite, *k*_2_ the activation rate, *k* _out_ metabolite clearance and *k* the damage-formation rate. Both hazard structures were fitted under the reduced configuration (RED, the one-compartment damage equation of §2.1) and the full one (FULL, the three compartments above). This compartmental axis concerns the damage model, not the 2×2 of §2.2. Integrator, quadrature and solver settings are in §S12.1.

### 3.2 Parameter estimation

Parameters were estimated by maximum likelihood on the conditional binomial likelihood, the negative log-likelihood being minimised by Nelder–Mead from 6–30 log-uniformly drawn starting points.^19^ Toxicokinetic parameters were fixed at measured values. For diazinon they are those of the study that produced the survival data.^20^ For dimethoate none has been measured in any freshwater invertebrate, so the three-compartment model is run only with the measured *G. pulex* values of the two compounds that bracket dimethoate in hydrophobicity, imidacloprid and aldicarb, as the two ends of a range, with metabolite clearance assumed within the range measured for three other anticholinesterase metabolites in this species^21^ (§S8.1, Table M2). Background mortality was estimated from each experiment’s controls. Every fitted and fixed value is in §S13 and Table M2. The AChE-P recovery rate was bounded at *t* ≥ 300 d, an assumption rather than a measurement (the recovery fitted to clean-water acetylcholinesterase data in diazinon-exposed *Daphnia magna* takes about 17 d from 60% inhibition to 95% of control^18^); results without it are in §S9.2. The chlorpyrifos–carbaryl parameters come from the published Threshold Damage Model without fitting (§S12.2, §S13.3).

The tolerance shape *β* (log-logistic shape) is held equal for both structures at 5.0, a convention rather than an estimate; estimated, it is 1.1–3.0 for dimethoate where not at a bound and 2.2–2.4 for diazinon (§S9.3). Holding *β* equal isolates the hazard-structure comparison; letting each structure fit its own would vary two things at once (§5.1). §4.4 reports how the margins depend on it.

### 3.3 Model comparison metrics

Three metrics — ΔNLL (NLL_SD_ − NLL_PM_, positive favouring GUTS-PM), gap closure (the fraction of the reduced-framework calibration gap closed by restoring compartments), and Nash–Sutcliffe efficiency — are defined in §S12.3. Inference uses log-scale delta-method intervals for the carry-over ratios (§S12.3), percentile bootstrap intervals for the ΔNLL margins of §4.4 (§S10.2), and a difference of log odds ratios for the chlorpyrifos–carbaryl order effect (§4.4).

### 3.4 Calibration and validation

Parameters were fitted on one dataset, then held fixed to predict independent datasets; for dimethoate, five calibration strategies varying concentration range and number of exposure regimes were compared, as was done for propiconazole^22^ (§S11).

### 3.5 Data sources

All survival data are for *Gammarus pulex*: a 4-day LC50 and pulse treatments for dimethoate,^3^ a sequential chlorpyrifos–carbaryl design in both orders,^23^ a pulsed diazinon experiment with three recovery intervals,^20^ and, for the carry-over ratios of §4.1, the fourteen-compound archive fitted under both structures (§S7, §S3.1). That archive is the only published set with toxicokinetics estimated once and shared across compounds; otherwise a comparison of hazard structures is confounded with kinetics.^2^ Sample sizes and exposure levels are in §S12.4.

### 3.6 Control simulation

*N* = 300 synthetic 96-hour datasets on the OECD TG 203 design were generated from each structure and fitted by both, with only the hazard parameters free and all others at their true values (§4.3). Protocol, parameters and results are in §S10.1.

## 4. Results

Four results are set out in turn. §4.1 tests the theorems of §2.3 in published counts and gives the first two results, the exclusion and the approach to the ceiling; §4.2 gives the third, which structure is the more conservative at matched parameters; §4.4 the fourth, what fitted comparisons show for two anticholinesterases. §4.3 is the premise the last of these needs: the calibration likelihood could not have made the choice. §4.4 also reports out-of-sample predictions, with the parameters frozen — a quantity distinct from calibration fit (Table S7).

### 4.1 Carry-over ratios exclude the cumulative structure

We measured the ratio from published counts alone, nothing fitted, wherever the repeated-pulse data^2^ allow it (§S6.1.4, §S15). Those data yield twenty-one two-pulse comparisons, from eleven of the fourteen compounds, in which the first pulse killed between 5 and 95% of the animals. Fifteen meet the counting rule (a ratio at or below unity counts only if the interval exceeded the compound’s recovery time; §2.3, §S6.1.1); the rule uses the individual-tolerance recovery times, at least as long as the other fit’s in every compound (Table S3) and so the stricter test. In those fifteen (eight compounds, four modes of action) the ratio is at or above unity in fourteen, and in seven, from three narcotics and an uncoupler, its confidence interval lies wholly above unity; six of the seven remain below *q* = 0.05 after Benjamini–Hochberg correction for multiple testing, the seventh at *q* = 0.063. The median is 1.29 and the maximum 4.42 (Figure 1, Table S2). Treatments within a compound share a cohort, so the compound is the independent unit: the largest ratio of each of the eight compounds is at or above unity, and four exclude it (§S6.1.4). The between-individual variation in killing rate that the theorem does not cover also flattens the concentration–response curve (susceptible animals die at lower concentrations), and the variation those curves allow falls short of what the ratios would require in five of the six comparisons that can be checked this way (§S6.1.3, Figure S2).

**Figure 1.**
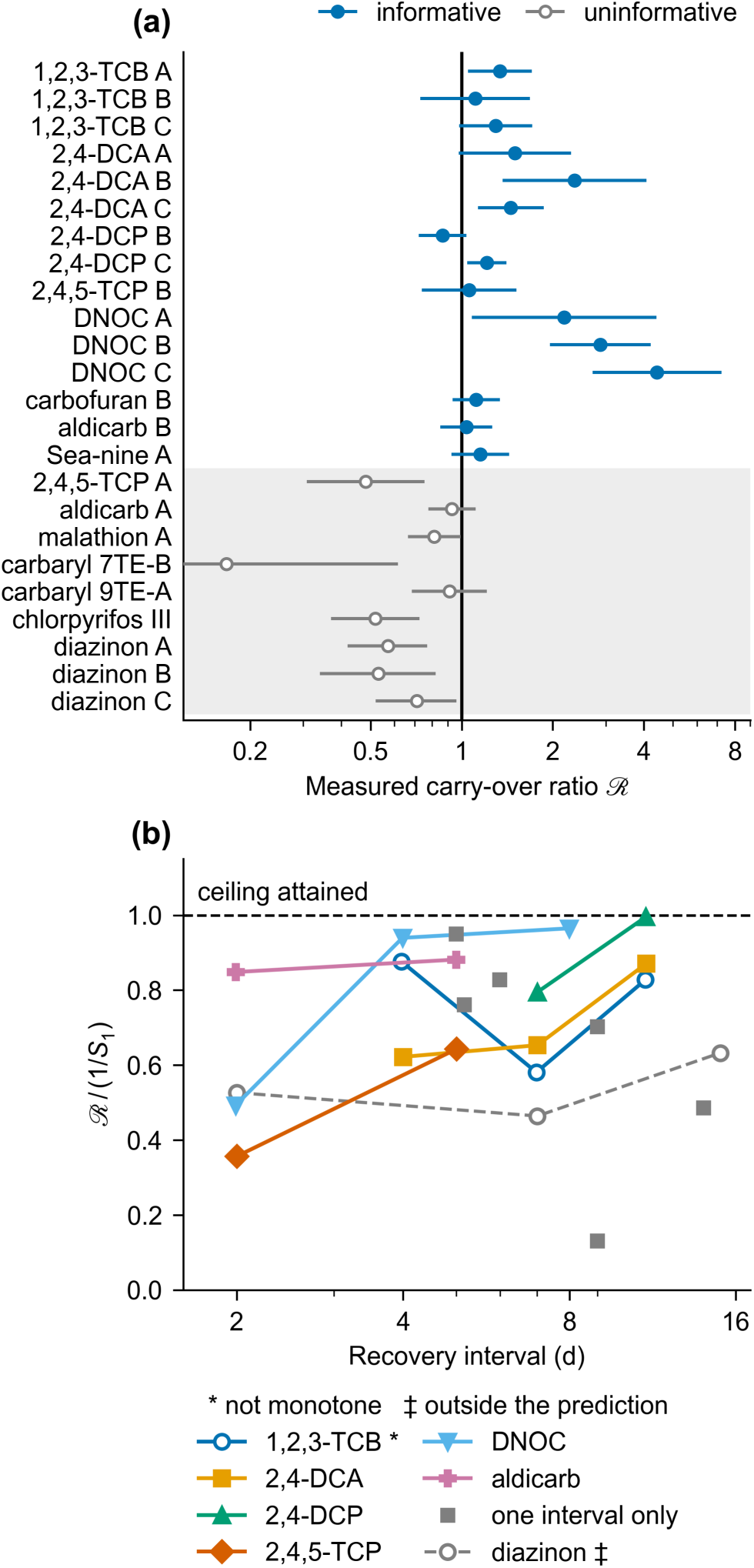
The carry-over ratio in published counts: test and ceiling. (a) The ratio recomputed from published counts, with 95% delta-method intervals on the log scale (§S12.3); the counting rule and the excluded comparisons (open) are in §S6.1.1 and §S3.4. (b) The measured ratios as fractions of their own ceiling, 1/*S*_1_ per treatment, against the recovery interval; treatments of one compound are joined. The peak-based structure attains the ceiling once damage has reset (Proposition S5), so joined points should rise; ten of twelve pairs do (sign test *p* = 0.019, §S6.2.2). Compounds tested at one interval only are shown unjoined. The dashed diazinon points lie below one recovery time, where the prediction does not apply, and are not counted.

A check that assumes no distribution agrees: such variation makes − log *S* grow ever more slowly at constant exposure as the susceptible die first, yet the linear form is preferred in all six compounds (§S6.1.3). The ratios also approach their ceiling 1/*S* (§2.3) as the interval lengthens: in five of the six compounds tested at more than one interval, with ten of twelve treatment pairs rising (Figure 1b, §S6.2.2).

### 4.2 Which structure is the more conservative at matched parameters

At matched parameters — the same damage trajectory, the hazards exchanged — the cumulative hazard is eventually the more conservative at every parameter value (Proposition S3, §S3.3). Over a test’s duration, which is the more conservative depends on two quantities the calibration does not determine — the threshold ratio *z*/*θ* and the tolerance shape *β* (Figure 2). Across twenty thousand draws the fraction in which the cumulative hazard is the more conservative crosses one half only between fourteen and twenty-eight days; at four days the peak-based structure is the more conservative in three draws of four. The median survival difference is at least 0.228 at every duration, so the choice always matters; and the ordering a validation test sees is not the ordering at the exposures an assessment covers (§5.3).

**Figure 2.**
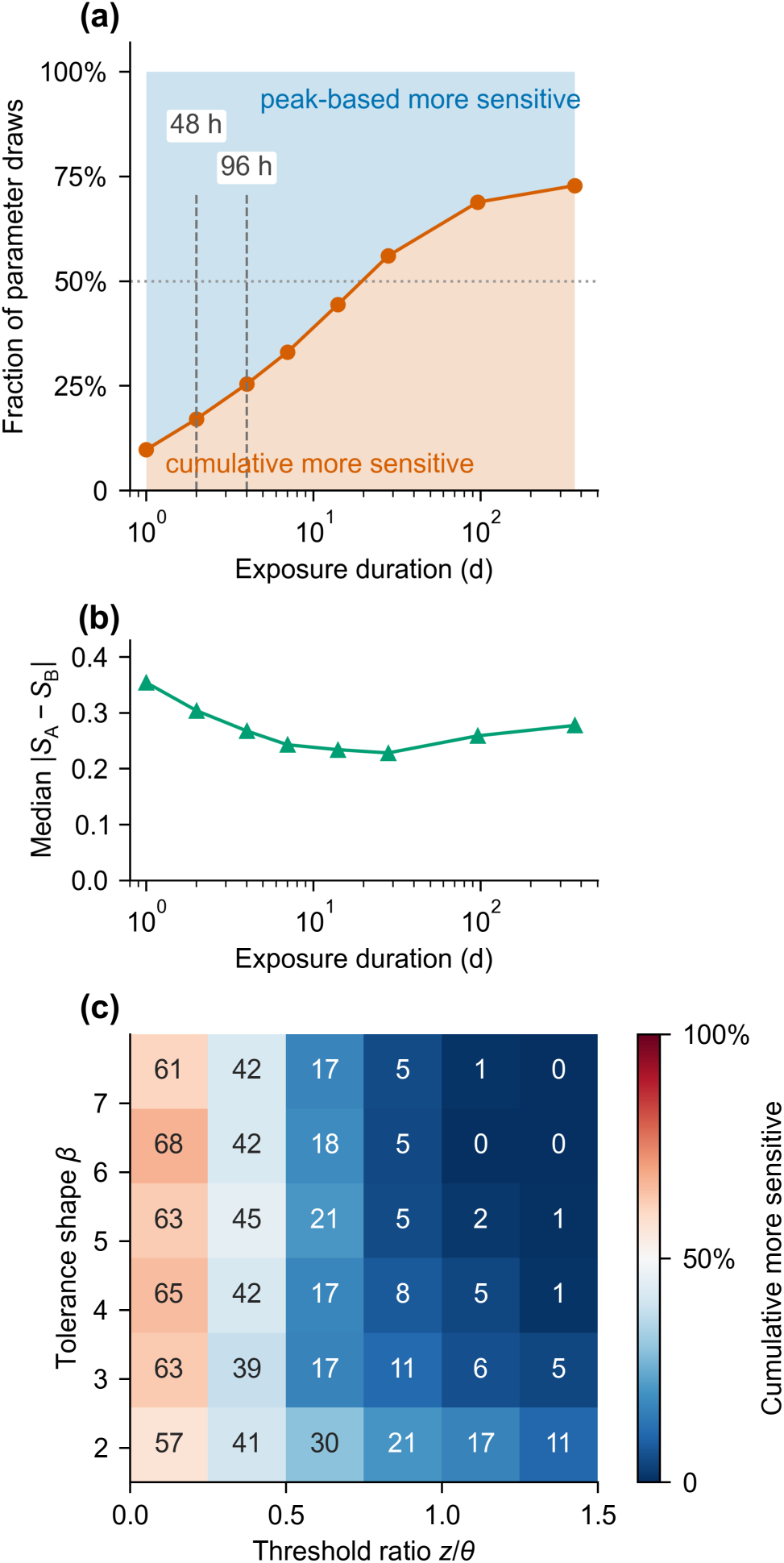
Which structure is the more conservative, at matched parameters. Both structures are driven by the same damage trajectory in a uniform population; 20,000 draws, scored where at least one kills between 5 and 95% (§S3.3). (a) The fraction of draws in which the cumulative hazard is the more conservative, against exposure duration; dashed lines mark standard acute-test endpoints. (b) The median absolute survival difference, never below 0.228, so the choice moves predicted survival by about a quarter of the population. The dip is the overtaking: the cumulative hazard grows with duration until its mortality passes the peak-based hazard’s, and the gap then re-opens toward the asymptotic separation (§S3.3). (c) The same fraction at four days, resolved by the threshold ratio *z*/*θ* and the tolerance shape *β*, which the calibration does not determine.

### 4.3 Control simulation: the likelihood has low power, not preference

The calibration likelihood has low power even when the rest of the model is correct. Six hundred synthetic four-day tests, half from each structure, were fitted with only the hazard free (Figure 3). This is the likelihood’s best case: the generating structure is one of the two candidates and the other parameters sit at their true values, so its performance bounds the power of any comparison at Tier-2C, the toxicokinetic–toxicodynamic tier of EU aquatic risk assessment for pesticides. The likelihood picks the generating structure 92% and 84% of the time for peak-based- and cumulative-generated data respectively, so it is not biased; but the margin is below two units in over half of replicates, median ΔNLL ±1.6 (§S10.1). A real comparison has less power still, with the toxicokinetics estimated and the damage model possibly misspecified. A larger gap in a real case study therefore points to something other than the hazard structure, as for dimethoate in the reduced framework (§4.4).

**Figure 3.**
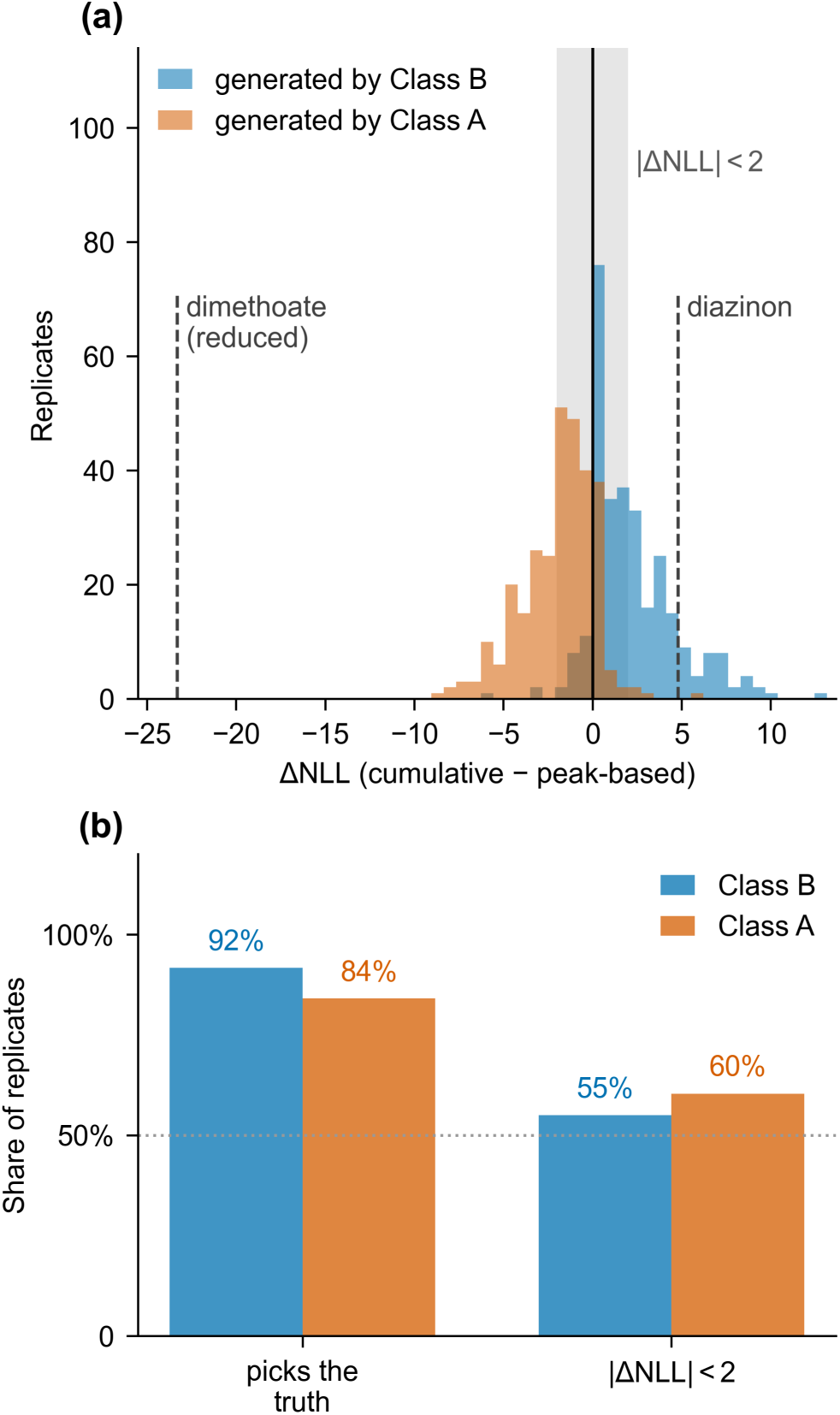
Control simulation: both generating structures, everything but the hazard at its true value. *N* = 300 synthetic four-day tests per generating structure (§S10.1). (a) The distribution of ΔNLL when the peak-based structure generated the data (blue) and the cumulative one (orange); shaded band, the two-unit margin; dashed lines, the calibration margins of dimethoate in the reduced framework and of diazinon (§4.4). (b) The likelihood picks the generating structure most of the time, yet the margin falls inside two units in over half of replicates; margins grow tenfold only when each structure also fits its own damage kinetics (§S7.3, Figure 4).

### 4.4 Anticholinesterases: fit comparison does not settle the structure

Two anticholinesterase datasets allow the fitted comparison whose power §4.3 bounds. **Diazinon** has toxicokinetics measured in the same study as a pulse experiment on *G. pulex* with two one-day pulses 2, 7 and 15 days apart.^20^ With those constants, three compartments and the recovery bound of §3.2, calibrated on the 2-day treatment, the peak-based structure leads by 4.8 units in calibration and 4.6 in prediction, and both 95% sampling intervals include zero (−15.6 to +24.7 and −16.0 to +25.5; §S10.2). The sign of the prediction margin follows two choices the data do not fix: without the recovery bound it is −3.6, and with the tolerance shape estimated for each structure −36.7 (§S9.2, §S9.3, §S13.2).

**Dimethoate** is an indirect organophosphate: the parent is converted to omethoate, which inhibits AChE. The data are a 4-day LC50 test and three 10-day pulse treatments.^3^ In the reduced framework, which needs no toxicokinetics, the cumulative structure fits better in calibration (ΔNLL = −23.3; refitting bootstrap −33.7 to −14.4, §S10.2) and in prediction of the pulse treatments (−125). Its size is consistent with the missing activation stage of §2.5, and §4.3 shows that a margin of this size does not come from the hazard alone. Lacking any toxicokinetic measurement for dimethoate in a freshwater invertebrate, the three-compartment model borrows the constants of its two neighbours in hydrophobicity (§3.2). With those constants, a metabolite clearance of 3.0 d^−1^ and the recovery bound, the calibration ΔNLL is +8.8 (imidacloprid constants) and −10.6 (aldicarb), closing 118% and 68% of the reduced-framework gap, and the prediction ΔNLL is +680.6 and +129.6; with the metabolite clearance estimated it is +493.8 and +72.2 (Tables S5–S7, Figure S6). Prediction favours the peak-based structure at both ends; the calibration sign depends on unmeasured constants, and without the bound the prediction margins fall to +21.2 and +16.8 (§S9.2). In the three-compartment fits damage does not decline within the test (Figure S5b), so dimethoate cannot discriminate on its own. Of five calibration strategies (aldicarb constants), only calibration on all three treatments holds every treatment above a Nash–Sutcliffe efficiency of +0.83 (Table S15, §S11).

A sequential design of two anticholinesterases does not choose between the structures either. Given a fortnight apart in both orders, carbaryl killed 31% of the animals when given first and 55% when it followed chlorpyrifos, against 33% and 47% predicted by the Threshold Damage Model published with the experiment.^23^ In a simplified re-implementation with the published parameters, both structures reproduce the direction of that order effect but not its size, and neither reproduces the smaller one for chlorpyrifos (§S13.3, Figure S10), so the experiment separates neither.

## 5. Discussion

### 5.1 Why the dilemma resisted: two choices in one question

The dilemma has been posed for two decades as a question about the mechanism of death: does an organism die because the damage it accumulated reached a lethal total, or because damage crossed its own threshold? Framed that way, the question cannot be answered: the two models on offer differ in two respects at once. Each variant pairs one hazard structure with one tolerance assumption (shared threshold or individual thresholds; §2.1, §2.2), and a likelihood comparison is asked to settle both choices from a single number. The control simulation shows how little that number can settle: with everything else correct, the likelihood separates them by less than two units in more than half of replicates (§4.3). Within the peak-based family the tolerance half of that pairing is not even identifiable: chance and between-animal difference enter survival only through their sum, so whether chance or between-animal difference produces the spread of times to death has no answer in survival data (§S3.1). What remains answerable is the hazard structure.

It is answerable only over part of an experiment: the two hazards coincide while damage rises and place the onset of mortality at the same event, and they diverge only once damage peaks and falls, which depends on the compound’s recovery chemistry and the test’s length, not on which structure is true (§S3.4).

Crossing the hazard and repair choices rather than confounding them changes what has to be decided. A claim about hazard structure is settled by what each structure can and cannot represent: the cumulative form can neither produce a carry-over ratio above one (§2.3, measured in §4.1) nor hold a lethal residue (the body residue at death) constant as exposure lengthens (§S6.3). Nothing requires the two to be decided together, and all four cells of Table 1 are occupied by models already in use.

### 5.2 What the measured ratio settles: exclusion and confirmation

What settles the choice is a measurement outside the fit, and the archive supplies it: a ratio above unity rules out GUTS-SD (§2.3), and fourteen of the fifteen informative comparisons reach or exceed unity, seven with confidence intervals excluding it (§4.1).

Exclusion is half of what the ratio gives. The other half is a prediction only the peak-based structure makes: once damage has reset the ratio equals 1/*S* exactly, a ceiling the experiment fixes and no fitting can move (Proposition S5). What is testable is the approach to that ceiling, which the archive shows in five of the six compounds tested at more than one interval and in ten of twelve treatment pairs (§4.1). A structure can be excluded by a number it cannot produce; it is confirmed by one it predicts before the data are seen, and the second is the stronger evidence.

### 5.3 Why calibration fit cannot settle the choice

Which structure a dossier should use cannot be settled by calibration fit, for three general reasons and a fourth that applies to activated compounds.

The first is that neither the threshold nor the recovery rate is identifiable on its own. Damage has no observable scale, so the recovery rate, threshold and killing rate are fixed only as a combination, and a reported value is one of many that fit equally well. Across six dimethoate fits the threshold spans 275-fold, but only 4.1-fold relative to the damage each fit produces (§S9.1). The cumulative structure infers a faster recovery than the peak-based structure (as GUTS-IT, §S3.1) in twelve of the fourteen compounds and never a slower one, by a factor of 4–20 where neither fit is on a bound (Figure 4, §S7.1).^2^ The rate a fit returns therefore depends on the structure assumed, and within either structure lets rate and threshold, both free in Tier-2C calibration,^24,25,26^ move together along a line the likelihood cannot resolve.

**Figure 4.**
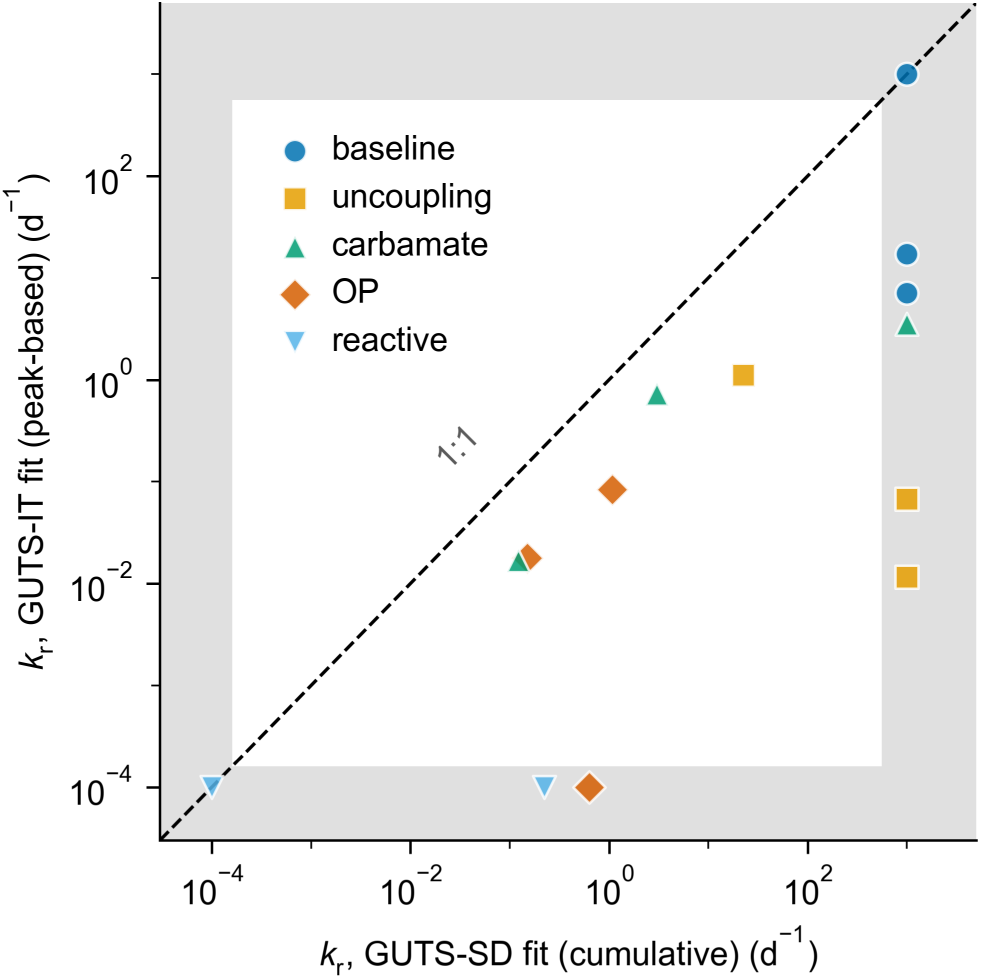
Fitted damage recovery rates under the two hazard structures. The fourteen compounds of Ashauer *et al.*,^2^ each fitted by the original authors under GUTS-SD and GUTS-IT, IT standing for the peak-based structure (§S3.1), with the toxicokinetics held identical; colour marks the mode of action. Twelve of the fourteen fall below the 1:1 line and two lie on it with both fits at a bound; in none does the cumulative structure infer the slower recovery. Grey bands mark the fitting bounds, 10^−4^ and 10^3^ d^−1^; nine compounds have at least one fit on a bound, so the direction holds and the magnitude is not estimated. Values in Table S3.

The second is that the cumulative structure does not depend on pulse history. Without the survivor selection a peak-based hazard produces, it predicts more deaths at a second pulse than the data show, in fourteen of the fifteen informative comparisons (§4.1), so a rule that prefers the more conservative prediction is biased towards it. The guidance suggests that where both variants pass validation the more conservative be used,^25^ but that validation is a goodness-of-fit judgement, which cannot separate them (§4.3). Where the carry-over ratio has been measured, the rule should not be reached at all. Where it has not, the rule does not settle the choice: the ordering of §4.2 is undetermined at the durations the rule is applied to, and fixed only at the longer exposures the assessment covers. A structure kept because it predicts more mortality, when a structural result has excluded it, shifts predicted survival by about a quarter of the population rather than adding conservatism (Figure 2b). The structure-dependent recovery rate and threshold act against each other by an amount the calibration output does not reveal (§S9.1).

The third is a matter of arithmetic between two parameterisations. The peak-based hazard is observationally equivalent to GUTS-RED-IT with a widened tolerance distribution (§S3.1), yet their thresholds differ by 1.2 to 15-fold where the survival curves agree to within 0.035 in probability (§S9.4). A threshold fitted under one parameterisation therefore cannot be used in an assessment that assumes the other, even though no fit would tell the assessor which one produced it.

A fourth source of error is specific to activated compounds. A one-compartment fit gives damage rising linearly where an activated compound’s trajectory rises faster, and a cumulative hazard integrating the linear trajectory mimics a peak-based hazard on the faster one (§2.5), so the omission can show up as an advantage for the structure that integrates. On dimethoate the reduced framework favours the cumulative structure by 23 units in calibration; with the compartments added, and toxicokinetics borrowed from two neighbouring compounds, 68–118% of the peak-based structure’s calibration deficit is recovered (gap closure, §3.3), and with the recovery bound the peak-based structure predicts the pulse treatments better (§4.4). How much of it is the framework cannot be settled without measured dimethoate toxicokinetics. On diazinon, where the toxicokinetics are measured, the margins lie within sampling variation and change sign with the recovery bound and tolerance shape (§4.4).

Model comparisons within the reduced framework therefore cannot by themselves establish the hazard mechanism of a compound whose activation is staged — organophosphates, pro-drugs, azole–pyrethroid combinations.

For organophosphates the evidence that bears on the structure comes from measuring the damage rather than inferring it. Kretschmann *et al.* moved diazinon-exposed *Daphnia magna* to clean water after 21 h and recorded acetylcholinesterase and immobilisation together:^18^ animals went on being immobilised while the enzyme was still falling, and once it began to recover the unaffected fraction stayed constant, although it stayed for a further thirteen hours below the threshold the same group later fitted to this series.^27^ With the measured enzyme trajectory as input, the peak-based hazard predicts this plateau at any parameter value; with that threshold and the killing rate fitted with it, the cumulative hazard predicts survival falling from 22.5% to 5–12% over those hours, an outcome with probability below 0.01 for the nine animals then unaffected, and refitted to the whole series it remains 4.2 log-likelihood units worse (§S3.2). This is one experiment on one species. Survival data alone add no second line: the cumulative structure’s fitted recovery rate reflects how much killing follows a pulse rather than how fast the lesion is repaired (Proposition S6).

### 5.4 Scope and limitations

Four qualifications bound what the structural case establishes. The exclusion of §4.1 and the ceiling it approaches are two readings of one archive; the lethal residue (§S6.3) is independent of it but constrains rather than excludes, and three further considerations are consistent with the peak-based structure without discriminating (§§S6.4–S6.6). The evidence reaches as far as those data do: narcotics in a fish, and narcotics and an uncoupler in *Gammarus*; for organophosphates it is one experiment on measured damage in *Daphnia magna* (§5.3). The peak-based hazard is validated within the proportional-repair sub-framework (*K* → ∞). For a receptor agonist a saturating repair term costs a parameter without improving the fit, *K* being unidentified over seven orders of magnitude, which independently justifies the proportional limit.^7^ For dimethoate the three-compartment comparison rests on unmeasured constants and changes sign within their plausible range (§4.4, Figure S6).

Reduced models differ from the full model in what they can represent at all. Deaths after exposure ends require damage still to be rising, and in the one-compartment framework damage peaks the moment exposure stops; the peak-based hazard freezes with the damage (Proposition S2, §S3.2). No reduced peak-based model can therefore produce post-exposure mortality, whatever its killing rate, while the cumulative structure can.^15^ Restoring the metabolite compartment removes the limitation, since damage there goes on forming for days after a pulse ends (§S8.2).

A further limit is quantitative: in the sequential chlorpyrifos–carbaryl design both structures predict an increase in carbaryl mortality of 1.4 to 3.6 percentage points, against 24 observed (§4.4, §S13.3), so for sequential pulses of unlike compounds the framework’s predictions are unvalidated rather than conservative — the companion paper’s distinction,^7^ applying to the exposure pattern Tier-2C is most often asked to cover.

For the organophosphates the support is that one experiment (§5.3), a level below a measured ratio: the designs outside the archive leave intervals shorter than the compound’s own recovery time, where a ratio at or below unity is uninformative by construction (§S3.4). For the carbamates the two informative ratios are above unity, with confidence intervals spanning it (Table S2). With the published data and measured toxicokinetics, fit comparison does not settle it either: the diazinon margins depend on the recovery bound and the tolerance shape (§4.4).

The falsification condition of §2.3 applies to the structure this paper recommends, and on one compound it fails: a peak-based hazard predicts flat survival once the damage input stops, yet across the published clean-water experiments on imidacloprid animals go on dying after transfer,^28,29^ and the companion analysis rejects the reduced peak-based model on these data (Proposition S2).^7^ The rejection reverses once the damage cascade is deepened,^7^ because Proposition S2 requires the input to the damage state to have reached zero, and receptor-bound imidacloprid keeps it running below what a residue measurement detects (§S3.2).

The imidacloprid reversal moves neither axis of the classification, which limits what the classification settles. Two further decisions can each decide which structure fits better: how deep the damage cascade runs (§5.3), and whether a finite receptor pool caps the rate at which damage forms, as it does for the neonicotinoid thiacloprid in *G. pulex*.^30^ Neither is an axis of the 2×2, so a fully specified comparison can still be reversed by changing one of them: the 2×2 is the minimum set of decisions to make explicit, not the complete set.

For narcotics and uncouplers both decisions are settled by what is known: no activation stage is involved, the uncoupler acting directly on the mitochondrial membrane;^31^ the uncoupler lesion clears with the exposure, since survival stops declining once uncoupler exposure ends but not after the anticholinesterases;^7^ and the uncouplers’ concentration–response series show no plateau that a saturating receptor pool would produce (§S6.1.3). For the anticholinesterases the cascade’s depth has been measured only for diazinon, and for the neonicotinoids neither question is settled. The organophosphate recommendation below therefore rests on the measured-damage test of §5.3 and is provisional; other compounds need the same two questions settled first.

### 5.5 Recommendation and outlook

We propose the designation **GUTS-PM** (Peak Maximum) for the peak-based hazard. Within the reduced framework the peak-based hazard is observationally equivalent to GUTS-REDIT (§S3.1); the name is needed outside it, where the finite killing rate that distinguishes it becomes identifiable once the damage model is mechanistic (§S9.4). The same identity settles an old observation: GUTS-IT fits pulse–recovery data well because, observationally, it is the peak-based hazard.

GUTS-SD should be restricted in domain, not withdrawn, and three things argue for keeping it.

The first is that GUTS-SD has a mechanism of its own, but not the one usually assumed: independent molecular killing events give the constant death rate of Class B, not the rising rate of Class A (§S6.5). A hazard that follows the damage level needs a rate rising with the damage already present, and that arises in two ways. Internally, accumulated damage leaves the organism progressively more fragile, or a reserve is consumed (§S6.5); externally, an encounter whose outcome the animal’s impairment decides gives the same shape, as does a lesion that impairs feeding rather than killing.^32^

The second is coverage: at matched parameters GUTS-SD is the more conservative at long exposure (§4.2) — an advantage that disappears once the parameters are fitted rather than matched (§S3.3) — and it alone among the reduced variants generates post-exposure mortality (§5.4). Those are grounds for a comparator, not for preferring it where the ratio has excluded it.

The third is that an acute test excludes by design the predators, pathogens and competition the external form requires, which a field exposure includes.

The structural argument is not specific to any chemistry — it asks only that death be a threshold crossing (§S6.4) — but §5.4’s further decisions need the activation cascade and receptor pool to be known. On the exclusion and confirmation results of §4.1, Class B is the appropriate default Tier-2C structure for narcotics and uncouplers, and, provisionally, for organophosphates on the measured-damage test of §5.3, which a clean-water phase with acetylcholinesterase measured in *G. pulex* would confirm or overturn (§S14). Carbamates, whose inhibition reverses spontaneously, remain open. A reduced Class B model must not be used where post-exposure mortality is plausible (§5.4). GUTS-SD should be fitted alongside GUTS-PM as a mandatory comparator, and neither structure’s fitted recovery rate read as a recovery time (Figure 4; Proposition S6 for GUTS-SD), including where it sets the depuration time that spaces validation pulses.^25^ Where they agree the choice does not matter; where they diverge, the response is a repeated-pulse experiment with recovery between exposures, not a decision on the calibration likelihood (§S14; §S14.4 for the classes where that design is impractical).

## 6. Associated Content

### Supporting Information

Analytical derivation of the four cells; the established GUTS variants as limiting cases; the peak-based hazard and the individual-tolerance family — their observational equivalence, the conservatism ordering scan and the observation-window conditions; staged activation and the *t^n^* rise of damage; Michaelis–Menten repair and why the data cannot resolve it; the evidence independent of fitting — the carry-over ratio and its ceiling, with the exclusion and confirmation theorems and their proofs, the lethal residue, the frailty bound, and three consistent but non-discriminating considerations; the fourteen-compound archive: fitted parameters and likelihoods; compartmental structure and gap closure; parameter identifiability; the control simulation and the sampling variability of the margins; the choice of calibration data; materials and methods in full; parameter tables, the diazinon results and the chlorpyrifos–carbaryl experiment; the design conditions for a decisive experiment and what to do where they are impractical; code and data availability. Tables S1–S22 and M1–M2, and Figures S1–S10 (PDF). This material is available with this preprint.

### Data and code availability

All data are from published sources, listed in §S12.4. Analysis scripts reproducing the reported numbers and tables (the exceptions are listed in §S15) and the figures listed there are archived at Zenodo (https://doi.org/10.5281/zenodo.23488393).^33^

## 7. Author Information

### Notes

No new experiments were performed and no animals were handled by the authors; all analyses use published data. The authors declare no competing financial interest.

### Author contributions

Jong-Hyeon Lee: conceptualization, methodology, software, formal analysis, investigation, data curation, writing — original draft, writing — review and editing, visualization. Chan Young Joe: software, validation, formal analysis, writing — review and editing.

## 8. Acknowledgments

This work was supported by the Korea Environmental Industry & Technology Institute (KEITI) with funding from the Korea Ministry of Environment (MOE) through the projects “Development of chemical emission factors for consumer products” (RS-2020-KE000904) and “Development of cumulative exposure assessment and prediction for biocide” (RS-2018-KE000099). AI-assisted tools were used during manuscript preparation for editorial refinement and for implementing, running and checking analysis code; all scientific content, analysis and interpretation are the responsibility of the authors.

## References

1. Li, H.; Lan, Q.; You, J. Application of TKTD Models in Predicting Time-Resolved Bioaccumulation and Toxicity of Pesticides to Aquatic Organisms. Aquat. Toxicol. 2025, 286, 107489. DOI: 10.1016/j.aquatox.2025.107489.

2. Ashauer, R.; O’Connor, I.; Hintermeister, A.; Escher, B. I. Death Dilemma and Organism Recovery in Ecotoxicology. Environ. Sci. Technol. 2015, 49 (16), 10136–10146. DOI: 10.1021/acs.est.5b03079.

3. Ashauer, R.; Albert, C.; Augustine, S.; Cedergreen, N.; Charles, S.; Ducrot, V.; Focks, A.; Gabsi, F.; Gergs, A.; Goussen, B.;, et al. Modelling Survival: Exposure Pattern, Species Sensitivity and Uncertainty. Sci. Rep. 2016, 6, 29178. DOI: 10.1038/srep29178.

4. Sowa, G.; Bednarska, A. J.; Laskowski, R. Mortality Pattern of *Poecilus cupreus* Beetles after Repeated Topical Exposure to Insecticide — Stochastic Death or Individual Tolerance? Environ. Sci. Technol. 2024, 58 (4), 1854–1864. DOI: 10.1021/acs.est.3c08031.

5. Ashauer, R. Correspondence on “Mortality Pattern of *Poecilus cupreus* Beetles after Repeated Topical Exposure to Insecticide — Stochastic Death or Individual Tolerance?” Environ. Sci. Technol. 2024, 58 (24), 10874–10876. DOI: 10.1021/acs.est.4c03056.

6. Sowa, G.; Bednarska, A. J.; Laskowski, R. Rebuttal to Correspondence on “Mortality Pattern of *Poecilus cupreus* Beetles after Repeated Topical Exposure to Insecticide — Stochastic Death or Individual Tolerance?” Environ. Sci. Technol. 2024, 58 (24), 10877–10880. DOI: 10.1021/acs.est.4c04127.

7. Lee, J.-H.; Joe, C. Y. The Test Used to Choose Between Pesticide Survival Models Tracks Lesion Persistence, Not Hazard Structure. bioRxiv preprint, 2026. DOI: 10.64898/2026.11.03.764107.

8. Petronilli, V.; Miotto, G.; Canton, M.; Brini, M.; Colonna, R.; Bernardi, P.; Di Lisa, F. Transient and Long-Lasting Openings of the Mitochondrial Permeability Transition Pore Can Be Monitored Directly in Intact Cells by Changes in Mitochondrial Calcein Fluorescence. Biophys. J. 1999, 76 (2), 725–734. DOI: 10.1016/S0006-3495(99)77239-5.

9. Fulton, M. H.; Key, P. B. Acetylcholinesterase Inhibition in Estuarine Fish and Invertebrates as an Indicator of Organophosphorus Insecticide Exposure and Effects. Environ. Toxicol. Chem. 2001, 20 (1), 37–45. DOI: 10.1002/etc.5620200104.

10. Schieber, M.; Chandel, N. S. ROS Function in Redox Signaling and Oxidative Stress. Curr. Biol. 2014, 24 (10), R453–R462. DOI: 10.1016/j.cub.2014.03.034.

11. Ashauer, R.; Brown, C. D. Toxicodynamic Assumptions in Ecotoxicological Hazard Models. Environ. Toxicol. Chem. 2008, 27 (8), 1817–1821. DOI: 10.1897/07-642.1.

12. Lu, S. C. Regulation of Glutathione Synthesis. Mol. Aspects Med. 2009, 30 (1–2), 42–59. DOI: 10.1016/j.mam.2008.05.005.

13. Mason, H. J. The Recovery of Plasma Cholinesterase and Erythrocyte Acetylcholinesterase Activity in Workers after Over-Exposure to Dichlorvos. Occup. Med. 2000, 50 (5), 343–347. DOI: 10.1093/occmed/50.5.343.

14. Miller, F. J.; Schlosser, P. M.; Janszen, D. B. Haber’s Rule: A Special Case in a Family of Curves Relating Concentration and Duration of Exposure to a Fixed Level of Response for a Given Endpoint. Toxicology 2000, 149 (1), 21–34. DOI: 10.1016/S0300-483X(00)00229-8.

15. Jager, T. Identifying and Predicting Delayed Mortality with Toxicokinetic–Toxicodynamic Models. Environ. Toxicol. Chem. 2024, 43 (5), 1030–1035. DOI: 10.1002/etc.5833.

16. Lee, J.-H.; Landrum, P. F.; Koh, C.-H. Prediction of Time-Dependent PAH Toxicity in *Hyalella azteca* Using a Damage Assessment Model. Environ. Sci. Technol. 2002, 36 (14), 3131–3138. DOI: 10.1021/es011202d.

17. Liess, M.; Foit, K.; Knillmann, S.; Schäfer, R. B.; Liess, H.-D. Predicting the Synergy of Multiple Stress Effects. Sci. Rep. 2016, 6, 32965. DOI: 10.1038/srep32965.

18. Kretschmann, A.; Ashauer, R.; Hitzfeld, K.; Spaak, P.; Hollender, J.; Escher, B. I. Mechanistic Toxicodynamic Model for Receptor-Mediated Toxicity of Diazoxon, the Active Metabolite of Diazinon, in *Daphnia magna*. Environ. Sci. Technol. 2011, 45 (11), 4980–4987. DOI: 10.1021/es1042386.

19. Jager, T.; Albert, C.; Preuss, T. G.; Ashauer, R. General Unified Threshold Model of Survival — a Toxicokinetic-Toxicodynamic Framework for Ecotoxicology. Environ. Sci. Technol. 2011, 45 (7), 2529–2540. DOI: 10.1021/es103092a.

20. Ashauer, R.; Hintermeister, A.; Caravatti, I.; Kretschmann, A.; Escher, B. I. Toxicokinetic and Toxicodynamic Modeling Explains Carry-Over Toxicity from Exposure to Diazinon by Slow Organism Recovery. Environ. Sci. Technol. 2010, 44 (10), 3963–3971. DOI: 10.1021/es903478b.

21. Ashauer, R.; Hintermeister, A.; O’Connor, I.; Elumelu, M.; Hollender, J.; Escher, B. I. Significance of Xenobiotic Metabolism for Bioaccumulation Kinetics of Organic Chemicals in *Gammarus pulex*. Environ. Sci. Technol. 2012, 46 (6), 3498–3508. DOI: 10.1021/es204611h.

22. Nyman, A.-M.; Schirmer, K.; Ashauer, R. Toxicokinetic-Toxicodynamic Modelling of Survival of *Gammarus pulex* in Multiple Pulse Exposures to Propiconazole: Model Assumptions, Calibration Data Requirements and Predictive Power. Ecotoxicology 2012, 21 (7), 1828–1840. DOI: 10.1007/s10646-012-0917-0.

23. Ashauer, R.; Boxall, A. B. A.; Brown, C. D. Modeling Combined Effects of Pulsed Exposure to Carbaryl and Chlorpyrifos on *Gammarus pulex*. Environ. Sci. Technol. 2007, 41 (15), 5535–5541. DOI: 10.1021/es070283w.

24. Jager, T.; Ashauer, R. Modelling Survival under Chemical Stress: A Comprehensive Guide to the GUTS Framework; Leanpub, 2023. https://leanpub.com/guts_book.

25. EFSA PPR Panel (EFSA Panel on Plant Protection Products and their Residues); Ockleford, C.; Adriaanse, P.; Berny, P.; Brock, T.; Duquesne, S.; Grilli, S.; et al. Scientific Opinion on the State of the Art of Toxicokinetic/Toxicodynamic (TKTD) Effect Models for Regulatory Risk Assessment of Pesticides for Aquatic Organisms. EFSA J. 2018, 16 (8), e05377. DOI: 10.2903/j.efsa.2018.5377.

26. EFSA PPR Panel. Guidance on Tiered Risk Assessment for Plant Protection Products for Aquatic Organisms in Edge-of-Field Surface Waters. EFSA J. 2013, 11 (7), 3290. DOI: 10.2903/j.efsa.2013.3290.

27. Kretschmann, A.; Ashauer, R.; Hollender, J.; Escher, B. I. Toxicokinetic and Toxicodynamic Model for Diazinon Toxicity — Mechanistic Explanation of Differences in the Sensitivity of *Daphnia magna* and *Gammarus pulex*. Environ. Toxicol. Chem. 2012, 31 (9), 2014–2022. DOI: 10.1002/etc.1905.

28. Li, H.; Zhang, Q.; Su, H.; You, J.; Wang, W.-X. High Tolerance and Delayed Responses of *Daphnia magna* to Neonicotinoid Insecticide Imidacloprid: Toxicokinetic and Toxicodynamic Modeling. Environ. Sci. Technol. 2021, 55 (1), 458–467. DOI: 10.1021/acs.est.0c05664.

29. Focks, A.; Belgers, D.; Boerwinkel, M.-C.; Buijse, L.; Roessink, I.; Van den Brink, P. J. Calibration and Validation of Toxicokinetic-Toxicodynamic Models for Three Neonicotinoids and Some Aquatic Macroinvertebrates. Ecotoxicology 2018, 27 (7), 992–1007. DOI: 10.1007/s10646-018-1940-6.

30. Raths, J.; Schinz, L.; Mangold-Döring, A.; Hollender, J. Elimination Resistance: Characterizing Multi-compartment Toxicokinetics of the Neonicotinoid Thiacloprid in the Amphipod *Gammarus pulex* Using Bioconcentration and Receptor-Binding Assays. Environ. Sci. Technol. 2023, 57 (24), 8890–8901. DOI: 10.1021/acs.est.3c01891.

31. Castilho, R. F.; Vicente, J. A. F.; Kowaltowski, A. J.; Vercesi, A. E. 4,6-Dinitro-o-cresol Uncouples Oxidative Phosphorylation and Induces Membrane Permeability Transition in Rat Liver Mitochondria. Int. J. Biochem. Cell Biol. 1997, 29 (7), 1005–1011. DOI: 10.1016/S1357-2725(97)00041-1.

32. Nyman, A.-M.; Hintermeister, A.; Schirmer, K.; Ashauer, R. The Insecticide Imidacloprid Causes Mortality of the Freshwater Amphipod *Gammarus pulex* by Interfering with Feeding Behavior. PLoS One 2013, 8 (5), e62472. DOI: 10.1371/journal.pone.0062472.

33. Lee, J.-H.; Joe, C. Y. Code for: Hazard Structure and Repair Kinetics: A 2×2 Classification that Resolves the Stochastic-Death / Individual-Tolerance Dilemma; Zenodo, 2026. DOI: 10.5281/zenodo.23488393.

