## Supporting Information for "Hazard Structure and Repair Kinetics: A 2×2 Classification that Resolves the Stochastic-Death / Individual-Tolerance Dilemma"

#### Contents


| Section | Tables | Figures |
|---|---|---|
| **Part I. Theory** | | |
| **S1** The Four Cells of the 2×2 Classification: What Each Predicts and Why | — | S1 |
| **S2** Existing Models as Limiting Cases | — | — |
| **S3** The Peak-Based Hazard and the Individual-Tolerance Family | S1, S1b | — |
| **S4** Staged Activation and the $t^n$ Rise of Damage | — | — |
| **S5** Saturating (Michaelis–Menten) Repair and Why These Data Cannot Identify It | — | — |
| **Part II. Model Structure and Parameter Estimation** | | |
| **S6** Evidence Independent of Fitting: the Carry-Over Ratio, Its Ceiling and the Lethal Residue | S2 | S2, S3 |
| **S7** Fourteen Compounds: What the Fitted Parameters and the Likelihood Each Show | S3 | S4 |
| **S8** Compartmental Structure and Gap Closure | S4, S5, S6, S7 | S5, S6 |
| **S9** Parameter Identifiability | S8, S8b, S9, S10, S11, S12 | — |
| **S10** Control Simulation and the Sampling Variability of the Margins | S13, S14 | — |
| **S11** Choosing the Calibration Data: Only Combined Calibration Holds Every Treatment | S15, S16 | S7, S8, S9 |
| **Part III. Procedures, Parameter Tables, Design and Deposit** | | |
| **S12** Materials and Methods (Detailed) | M1, M2 | — |
| **S13** Parameter Tables, the Diazinon Results and the Chlorpyrifos–Carbaryl Experiment | S17, S18, S19, S20 | S10 |
| **S14** Designing a Decisive Experiment: Three Conditions and What Each Outcome Settles | S21, S22 | — |
| **S15** Code and Data Availability | — | — |


Tables: 27 · Figures: 10

Part I is analytical and can be read independently of the case studies. Part II holds the structural evidence, the fitted comparisons and the identifiability and separability controls. Part III holds procedures, parameter tables, the prospective design and the deposit.

### Part I. Theory

*How this Part supports the main text.* §S1–S2 derive Table 1 and recover the established models as its limiting cases (§2.2, §2.4). §S3 carries the main text's third result — which structure is the more conservative at matched parameters (§4.2) — and the falsification and observation-window conditions that §2.3 and §S14 rely on. §S4–S5 supply the two mechanisms behind the reduced-framework comparison of §4.4: the $t^n$ rise of damage and a repair capacity the data cannot identify.

#### S1. The Four Cells of the 2×2 Classification: What Each Predicts and Why

Table 1 of the main text (§2.2) crosses two modelling choices — how damage is repaired, and how damage turns into a risk of death — and obtains four different kinds of survival curve. This section shows where each comes from. Each cell is written in two layers: first what it predicts and why, in words; then the short calculation that confirms it, which a reader who accepts the words can skip. All four use the same setting: constant exposure, with the body concentration $C_b$ already at steady state.

**The two choices in words.** *Repair.* Damage $D$ is formed at a rate set by the body concentration and removed by repair. Under proportional repair, removal speeds up as damage grows, like water draining from a tub through an open plughole: the fuller the tub, the faster it drains, so the level settles where inflow and outflow balance. Under constant-rate repair the drain has a fixed capacity; if inflow exceeds it, the level rises without limit. *Hazard.* Here "risk" means the risk accumulated so far — the cumulative hazard $H$ defined below — and the risk per unit time is called the death rate. Under the cumulative structure (Class A), the risk of death builds up for as long as damage stays above a threshold $z$, and every hour spent above it adds to it. Under the peak-based structure (Class B), the risk depends only on the highest damage reached so far, the running maximum $M(t)$ — a high-water mark that never goes down.

**Notation.** $[x]^+$ is $x$ when $x$ is positive and zero otherwise, so $[D-z]^+$ reads "how far damage is above the threshold, or nothing if it is below". $t \gg 1/k_r$ means after several recovery times, when damage has practically stopped changing; it is 95% of the way to steady state at $t = \ln 20/k_r \approx 3/k_r$. Survival is $S = e^{-H}$, where $H$ is the cumulative hazard: the larger $H$, the fewer survive.

All derivations follow a single organism. Under Class B its threshold $\theta$ is its own, and population survival follows by averaging over the tolerance distribution $f(\theta)$; under Class A the threshold $z$ is shared, so the single-organism result is already the population's (a distributed $z$ is GUTS proper, §S6.1.3). Background mortality adds $h_b t$ to every cumulative hazard; it is written out for Class B and omitted for Class A.

Throughout §S1 and §S6 the damage equation is written for the one-compartment case, $dD/dt = k_a C_b - k_r D$, in which the input to damage is the parent body concentration $C_b$. The three-compartment case of the main text replaces that input with the activated metabolite, $dD/dt = k_3 A - k_r D$ (eq. 3), and it is precisely that substitution which produces the curvature discussed in §S4. The damage-formation rate $k_a$ of the reduced model and the $k_3$ of the three-compartment model are kept as separate symbols throughout, so that neither is read as the other. Figure S1 shows the four cells with simulated damage trajectories and survival curves.

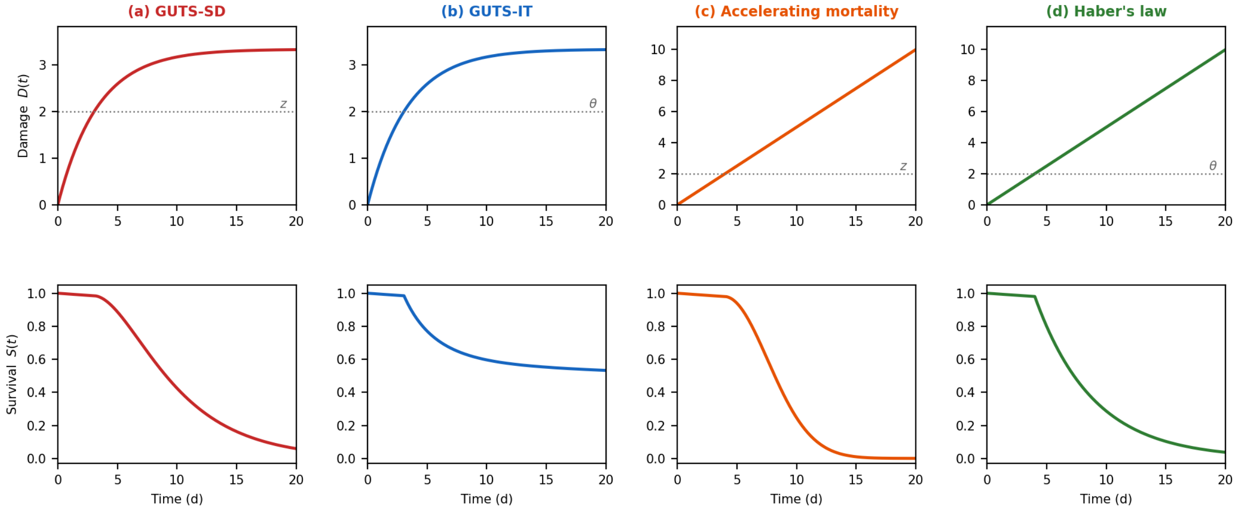


**Figure S1. Detailed behaviour of all four cells in the 2×2 classification.** (a) GUTS-SD, (b) GUTS-IT, (c) accelerating mortality and (d) Haber's law, one column per cell. Upper row: damage trajectories $D(t)$ under constant exposure. Lower row: the corresponding survival curves $S(t)$. One organism, threshold 2.0 ($z$ for the cumulative cells, $\theta$ for the peak-based ones), background hazard 0.005 $d^{-1}$; proportional repair $dD/dt = 1 - 0.3D$, constant-rate repair $dD/dt = 0.5$; $b_w = 0.15$, $k = 0.4$ (`figureS1_four_cells.py`). See §S1.

##### S1.1 Cumulative Hazard, Proportional Repair (GUTS-SD): Death Continues at a Steady Rate

**In words.** Damage rises and levels off at a steady value $D_{ss}$. If that level is above the threshold, every unit of time spent there adds the same amount of risk, so animals keep dying at a constant rate for as long as exposure lasts, and eventually all die.

Under proportional repair, the damage equation $\frac{dD}{dt} = k_a C_b - k_r D$ at constant body concentration $C_b$ yields:

$$D(t) = D_{ss}\left(1 - e^{-k_r t}\right), \quad D_{ss} = \frac{k_a C_b}{k_r}$$

Damage increases toward a finite steady state because repair speeds up proportionally with damage. For $t \gg 1/k_r$, with damage near steady state, the cumulative hazard becomes:

$$H_A(t; z) = \int_0^t b_w\ [D(\tau) - z]^+\ d\tau\ \approx\ b_w\ [D_{ss} - z]^+\ t$$

The survival of an organism with threshold $z$ thus decays exponentially at rate $b_w\,[D_{ss} - z]^+$. Whenever $D_{ss} > z$, every organism eventually dies — the signature of GUTS-SD.

**What to look for.** Once damage has levelled off, survival plotted on a logarithmic axis against time falls along a straight line that does not level off.

##### S1.2 Peak-Based Hazard, Proportional Repair (GUTS-IT): Deaths Stop Once Damage Levels Off

**In words.** The damage is the same as in §S1.1, but the risk is set by the high-water mark. Once damage levels off, the mark stops rising and no new risk is added. Animals whose tolerance lies below $D_{ss}$ are at risk only while damage is still rising — all of them die in the GUTS-IT limit, a fraction of them at a finite killing rate — and the rest survive however long the exposure continues.

Since $D(t)$ increases monotonically toward $D_{ss}$, the running maximum equals the current damage at all times, and the peak damage approaches $D_{ss}$. For $t \gg 1/k_r$ the toxicant part of the hazard therefore reaches a finite ceiling, to which background mortality adds:

$$H_B(t; \theta) = k\,[D_{ss} - \theta]^+ + h_b t$$

Setting aside the background term, an organism with $\theta > D_{ss}$ faces no toxicant hazard at all, while an organism with $\theta < D_{ss}$ faces a finite one, $k\,(D_{ss} - \theta)$, and survives it with probability $\exp\left(-k\,(D_{ss} - \theta)\right)$. As $k$ grows the survival of an exposed organism falls to zero and the cell reduces to GUTS-IT, in which crossing the threshold is itself fatal; at finite $k$ it is the broader structure §S3.1 describes.

**What to look for.** Mortality levels off at the fraction of animals whose tolerance lies below the steady damage level, and the LC50 settles at a fixed value instead of falling with test duration.

##### S1.3 Cumulative Hazard, Constant-Rate Repair: Mortality Accelerates

**In words.** Repair cannot keep up, so damage climbs steadily. Under the cumulative structure the excess above the threshold itself grows, so each day adds more risk than the day before: the death rate rises with time.

When repair runs at a fixed maximum rate regardless of damage level, the damage equation becomes $\frac{dD}{dt} = k_a C_b - k_{r,0}$. If the toxicant causes damage faster than this maximum rate ($k_a C_b > k_{r,0}$), damage grows linearly and does not saturate within the model:

$$D(t) = \left(k_a C_b - k_{r,0}\right) t \equiv v\, t$$

The cumulative hazard then integrates a linearly growing exceedance, which is zero until $t_z = z/v$, the time at which damage first crosses the threshold:

$$H_A(t; z) = \int_0^t b_w \,[v\tau - z]^+ \, d\tau = \frac{1}{2} b_w v \,(t - t_z)^2$$

for $t > t_z$.

The cumulative hazard grows quadratically, so the death rate, $b_w v\,(t - t_z)$, rises linearly with time instead of holding steady as it does under proportional repair, though both Class A cells share the same endpoint: every organism above the threshold eventually dies. We refer to this cell as accelerating mortality.

It is not a named GUTS variant, although GUTS-SD reaches it when repair is negligible ($k_r \to 0$), as GUTS-IT reaches Haber's law (§S1.4).

**What to look for.** The death rate — the fraction of the survivors dying each day — rises as the test goes on, instead of holding steady. It is rarely seen in acute tests: in most of them damage levels off within the test, which this cell does not allow; it requires continuing exposure to a compound the animal neither repairs nor clears. The same accelerating form — a delayed Weibull hazard of shape 2 — is the wear-out pattern of reliability engineering and, with staged damage (§S4), the multistage pattern of carcinogenesis; what these share is the mechanism of §S6.5, not a common toxicokinetic model.

##### S1.4 Peak-Based Hazard, Constant-Rate Repair: Haber's Law

**In words.** Damage climbs steadily, so the high-water mark keeps rising at a steady pace, and once the threshold is passed the death rate is constant. Doubling the concentration doubles the rate of climb, which halves both the time to reach the threshold and the wait after it; concentration times time to death is therefore constant — Haber's law.

With the same linearly growing damage trajectory as §S1.3 but the peak-based hazard, the running maximum equals current damage ($D(t) = vt$ is monotonically increasing), so:

$$H_B(t; \theta) = k[vt - \theta]^+ + h_b t = kv\,(t - t_\theta) + h_b t, \quad t > t_\theta = \theta/v$$

The hazard rate after threshold crossing is constant at $kv + h_b$. Death is exponentially distributed with a delay $t_\theta$. Haber's law, $C_{\text{ext}} \times t_d$ = const for the time to death $t_d$, follows because both parts of $t_d$ scale as $1/C_b$ once the damage rate is well above the fixed repair rate: the delay $t_\theta = \theta/\left(k_a C_b - k_{r,0}\right)$ approaches $\theta/\left(k_a C_b\right)$, and the mean wait after crossing, $1/(kv)$, does the same when background mortality is small; $C_b$ is proportional to $C_{\text{ext}}$ at toxicokinetic steady state.

**A second route, and a difference that matters.** Haber's law also follows from the proportional-repair cell (§S1.2) when repair is negligible ($k_r = 0$) and killing is immediate ($k \to \infty$) under constant exposure at toxicokinetic steady state (uptake $k_u$, elimination $k_e$, so $C_b^{ss} = k_u C_{\text{ext}}/k_e$). Damage then accumulates as $D(t) = k_a k_u C_{\text{ext}} t/k_e$, and death occurs when $D(t_d) = \theta$, giving:

$$t_d = \frac{\theta\, k_e}{k_a\, k_u\, C_{\text{ext}}} \quad \Longrightarrow \quad C_{\text{ext}} \cdot t_d = \frac{\theta\, k_e}{k_a\, k_u} = \text{const}$$

Here the law arises without any integration of hazard: the constant is set by the time for $C_b$ to carry damage from zero to $\theta$. The two routes differ in one respect. With constant-rate repair there is a body residue below which nothing dies: when $k_a C_b \le k_{r,0}$, repair keeps pace with formation, damage never rises, and $H_B = 0$ however long the exposure. That residue, $C_b^* = k_{r,0}/k_a$, is the critical body residue (CBR), consistent with the observation that organisms held below it survive indefinitely under narcotic exposure. Proportional repair gives the same behaviour organism by organism, below $C_b = k_r\theta/k_a$ (§S1.2); with no repair at all there is no such residue, because damage then accumulates without bound.

#### S2. Existing Models as Limiting Cases

Three established TKTD models — GUTS-IT, the Stress Addition Model (SAM) and the damage assessment model DAM — emerge as exact limiting cases of the peak-based survival integral (main text eq. 2, restated in §S2.1). GUTS-SD does not: it occupies the cumulative column (§S1.1).

##### S2.1 GUTS-IT and the Stress Addition Model as the $k \to \infty$ Limit

Survival under the peak-based hazard is an average over the tolerance distribution,

$$S(t) = \int_0^\infty \exp\left(-k\,[M(t) - \theta]^+ - h_b t\right) f(\theta)\, d\theta,$$

with $M(t) = \max_{\tau \le t} D(\tau)$. Fix $t$ and let $k$ grow. For an organism whose tolerance exceeds the peak, $[M(t) - \theta]^+ = 0$ and the integrand stays at $e^{-h_b t}$; for one whose tolerance is exceeded, $[M(t) - \theta]^+ > 0$ and the integrand falls to zero. Any exceedance at all becomes immediately lethal, which is the defining feature of GUTS-IT.

Taking that limit inside the integral needs justification, because a sequence of functions can converge to zero at every point while its integral does not. Here it is legitimate. Because $\exp\left(-k\,[M(t) - \theta]^+\right) \le 1$ for every $k$ and $\theta$, the integrand is at most $e^{-h_b t} f(\theta)$; that bound does not depend on $k$ and integrates to $e^{-h_b t}$, since $f$ is a probability density. The dominated convergence theorem then applies and the limit may be taken pointwise. The surviving fraction is then the mass of the tolerance distribution lying above the peak,

$$S(t) \;\to\; e^{-h_b t}\,[1 - F\,(M(t))],$$

where $F$ is the cumulative distribution function of $f$. This is the standard GUTS-IT survival equation.

The Stress Addition Model follows from the same limit. Combining the GUTS-IT limit ($k \to \infty$) with steady-state conditions ($t \to \infty$), proportional repair (the $K \to \infty$ limit of §2.1), and $J$ additive stressors, the multi-stressor survival becomes, with background mortality set aside,

$$S = 1 - F\left(\sum_{j=1}^{J} \sigma_j\right), \quad \sigma_j = \frac{k_{a,j}\, k_{u,j}}{k_{r,j}\, k_{e,j}} C_{\text{ext},j}$$

where $\sigma_j$ is the stress contributed by stressor $j$. This follows because under proportional repair with $J$ stressors, damage additivity gives $D_{ss} = \sum_j k_{a,j} C_{b,j}^{ss} / k_{r,j}$, and at toxicokinetic steady state $C_{b,j}^{ss} = k_{u,j} C_{\text{ext},j} / k_{e,j}$. Applying the GUTS-IT limit at steady state ($M = D_{ss}$) yields $S = 1 - F\left(\sum_j \sigma_j\right)$ — exactly the Stress Addition Model.[17]

##### S2.2 The Shared-Threshold Limit, DAM as Its Thresholdless Case, and What the Shared Threshold Means for the Lower Tail

When all organisms share the same threshold — written $D_0$ to keep it distinct from the Class A threshold $z$ — the tolerance density collapses to $f(\theta) = \delta(\theta - D_0)$. Substituted into the survival integral of §S2.1 the delta sifts out the single value $\theta = D_0$, leaving

$$S(t) = \exp\left(-k\left[M(t) - D_0\right]^+ - h_b t\right)$$

This is the shared-threshold form of the peak-based hazard. With $D_0 = 0$ and damage that does not decrease, as under constant exposure, the running maximum is the current damage and it reduces to the damage assessment model DAM, $S = \exp(-kD(t) - h_b t)$.[16]

The rest of this subsection asks what $D_0$ is, because the answer bears on the lower tail of the dose–response and therefore on protective endpoints. The running maximum makes identical individuals behave as if their tolerances differed: defining an effective individual tolerance $\theta_i = D_0 + U_i/k$ where $U_i$ follows a standard exponential distribution, individual $i$ survives if and only if $M(t) < \theta_i$, and the population survival equals $\exp\left(-k\left[M(t) - D_0\right]^+\right)$, background mortality set aside. The effective tolerance distribution is therefore a shifted exponential with location $D_0$ and scale $1/k$. $D_0$ can be read in two ways, and it means something different under each. Read as written, $D_0$ is a threshold every organism shares, and death above it is a matter of chance. Read through $\theta_i$, no organism has a tolerance of $D_0$ at all, since $U_i > 0$ almost surely; $D_0$ is instead the infimum of the tolerance distribution, the damage level below which nobody dies. The mean tolerance is then $D_0 + 1/k$ and its standard deviation $1/k$. The same symbol is a physiological constant in one reading and a population lower bound in the other, and the survival data cannot tell them apart. Both readings predict that nothing dies below $D_0$; what differs is whether that floor is a property of every animal or only of the exponential shape the running maximum imposes on the tolerance distribution — and that floor is the quantity a protective endpoint reports.

The shifted-exponential distribution the running maximum generates differs from the log-logistic or log-normal distributions typically fitted in ecotoxicological practice: it places zero probability below $D_0$, so mortality is predicted to be exactly zero for peak damage below $D_0$. Because low-effect endpoints such as $LC_{10}$ — and the $HC_5$ values built from them — are governed by the lower tail, the shared-threshold form of this subsection may be non-conservative there relative to a fitted log-logistic, and an explicit tolerance distribution $f(\theta)$ should be calibrated to data whenever low-effect percentiles drive the assessment.

The same equivalence between chance and between-animal difference holds for any tolerance density, not only a shared threshold (§S3.1).

#### S3. The Peak-Based Hazard and the Individual-Tolerance Family

This section makes four points about the peak-based structure (Class B). §S3.1 shows that it is the same thing as individual tolerance; §S3.2 states what would refute it, and where that has been seen; §S3.3 shows which structure predicts more deaths, and that the answer depends on how long exposure lasts; §S3.4 says when the two structures can be told apart at all. As in §S1, each result is stated first in words; the formal statement and proof that follow are there to be checked and can be skipped.

**Probability notation.** $\Pr(\cdot)$ is the probability of the event in brackets. A tolerance distribution $f(\theta)$ describes how thresholds vary between animals; its cumulative form $F(x)$ is the fraction of animals whose threshold lies below $x$, so $1 - F(x)$ is the fraction still alive once damage has reached $x$. The log-logistic distribution used throughout has a median $\theta$ and a shape $\beta$: the larger $\beta$, the more tightly thresholds cluster around the median, and the fewer animals are very sensitive. $f = \delta$ means that every animal has the same threshold. $E \sim Exp(1)$ is a random quantity whose chance of exceeding any value $x$ is $e^{-x}$. $\mathbb{E}[\cdot]$ is an average over animals. The symbols $[x]^+$ and $S = e^{-H}$ are as in §S1.

##### S3.1 Chance and Tolerance Cannot Be Told Apart Under the Peak-Based Hazard

Practitioners are used to treating stochastic death and individual tolerance as rival accounts of the same data. Under stochastic death the animals are alike and the ones that survive were lucky; under individual tolerance they differ and the ones that survive were tolerant. For the peak-based hazard the two are not rivals at all, and this section shows why.

**In words.** Give each animal a hidden safety margin, drawn at random, on top of its own threshold. Under the peak-based hazard an animal dies once damage has, at its highest, exceeded its threshold by more than that margin. A finite killing rate $k$ is exactly such a margin, of average size $1/k$ — the buffer referred to below: saying that an exceedance $x$ kills with probability $1 - e^{-kx}$ is the same as saying that the animal dies if $x$ is larger than a random margin $E/k$. The survival curve cannot tell an animal that survived because its threshold was high from one that survived because its margin was large; it sees only their sum, $\theta + E/k$. Within the peak-based structure, chance (the stochastic-death reading) and between-animal difference (the individual-tolerance reading) are therefore two descriptions of the same survival curve, mixed in any proportion, and no survival experiment can apportion them; the GUTS-SD model, whose hazard is cumulative (Class A), is a different matter. What the data can identify is the hazard structure — which feature of the damage trajectory the risk depends on — not the mix.

The GUTS-IT limit of §S2.1 ($k \to \infty$, the buffer shrinking to nothing) and the shared-threshold limit of §S2.2 (no spread but the buffer) are not special properties of those cases: both are corners of one identity, which holds for every finite $k$ and every $f$.

**Proposition S1.** *Let $E \sim Exp(1)$ be independent of $\theta \sim f$. For any tolerance density $f$ on $[0, \infty)$ and any $k > 0$,*

$$S(t) = e^{-h_b t} \int_0^\infty \exp\left(-k\,[M(t) - \theta]^+\right) f(\theta)\, d\theta = e^{-h_b t}\, \Pr\left(\theta + \frac{E}{k} > M(t)\right) = e^{-h_b t}\, [1 - F_\oplus(M(t))]$$

*where $\oplus$ denotes the distribution of a sum of independent random variables, so that $F_\oplus$ is the cumulative distribution function of the animal's own threshold plus its exponential buffer.*

*Proof.* Take one animal with tolerance $\theta$ and average over $\theta$ at the end. If $\theta \geq M(t)$ the animal was never pushed past its threshold: $k\,(M(t) - \theta) \leq 0$, so the exponential reserve is certainly the larger of the two and $\Pr(E > k\,(M(t) - \theta)) = 1$, which equals $\exp\left(-k\,[M(t) - \theta]^+\right)$ because the positive part is zero. If $\theta < M(t)$ the exceedance is positive and $\Pr(E > k(M(t) - \theta)) = \exp(-k(M(t) - \theta))$, which is again $\exp\left(-k[M(t) - \theta]^+\right)$. The two cases collapse into one expression precisely because $[\cdot]^+$ clips the negative case to zero. Averaging over $\theta$ gives the identity. ▫

The identity depends only on the hazard being a function of $M(t)$, and not at all on how $M(t)$ is generated, so it holds equally in the reduced and the full framework and under proportional or saturating repair.

**What it means for fitting.** The peak-based hazard belongs to the individual-tolerance family at **every** killing rate, not only in the limit: §S2.1 and §S2.2 are the two ends, $F_{\oplus} \to F$ and $f = \delta$. Two things follow for anyone fitting these models. First, fitting GUTS-PM with a finite killing rate and some tolerance spread returns the same survival curves as fitting GUTS-IT with a wider spread and no killing rate at all, so survival data cannot say how much of the observed variation is between-animal difference and how much is chance. The structural dichotomy of the 2×2 is therefore between Class A and Class B, with Class B and the IT family the same object — which also explains something long unexplained: GUTS-IT fits pulse-recovery data well because it is the peak-based structure, which adds no hazard for a repeated pulse once damage has reset (§S6.2), and that is the pattern those data show. Second, because the two parameterisations place the threshold differently, a value fitted under one cannot be carried into an assessment built on the other (§5.3).

The redundancy behind both statements — how much tolerance spread a finite killing rate is worth, and how far the two parameterisations place the threshold apart — is quantified with the other identifiability results in §S9.4.

##### S3.2 Flat Survival While Damage Falls: the Falsification Condition and Where It Has Been Tested

The same identity fixes what would refute the peak-based structure. Because $S(t)$ depends on the trajectory only through $M(t)$, and $M$ is constant once damage begins to decline, survival must be flat during any interval in which damage is falling.

**In words.** Once nothing more is being fed into the damage state, damage can only stay level or fall, whatever the repair process does. A high-water mark cannot rise while the water is falling, so the peak-based risk of death stops increasing and survival stays flat, apart from background deaths. Any death in that period that background mortality cannot explain refutes the peak-based structure — provided the input really has stopped.

**Proposition S2 (freezing of the peak-based hazard).** *Let damage obey $dD/dt = u(t) - R(D)$, with input $u(t) \geq 0$ and any repair function satisfying $R(D) \geq 0$. If $u(\tau) = 0$ for all $\tau \geq t_0$, then the running maximum $M(t) = \max_{\tau \leq t} D(\tau)$ is constant on $[t_0, \infty)$, and so is the Class B cumulative hazard $H_B = k\,[M(t) - \theta]^+$. Survival is therefore flat over that interval, whatever the form of $R$.*

*Proof.* For $\tau \geq t_0$ the input vanishes, so $dD/d\tau = -R(D(\tau)) \leq 0$ and $D$ is non-increasing on $[t_0, \infty)$. Hence for any $t \geq t_0$,

$$M(t) = \max\left\{M(t_0),\ \max_{t_0 \leq \tau \leq t} D(\tau)\right\} = \max\{M(t_0), D(t_0)\} = M(t_0),$$

the last step because $D(t_0) \leq M(t_0)$ by definition of the running maximum. Since $H_B$ depends on the trajectory only through $M$, it is constant on $[t_0, \infty)$, and $S = e^{-H_B}$ with it. ▫

The proof uses nothing about repair except that it never adds damage, so the result holds for every repair law. That is what makes it a falsification condition rather than the prediction of one model: no choice of repair kinetics, saturating or otherwise, can produce a death while the input is off.

The operative condition is that the input to the damage state has reached zero, and that is not the same as clearance of the parent compound. In a multi-stage trajectory the input to the terminal state is the preceding state, and the compound held at that stage — bound to receptors, say — can stay in the tissue while the rest clears, so that a total-residue measurement does not show whether the input has stopped.

That the operative pool can be persistent and invisible is not hypothetical: for a neonicotinoid in *Gammarus pulex* the receptor-bound fraction does not decline measurably while the free fraction clears, so a total-residue measurement sees the parent compound leaving and cannot see that the input to the damage state has not.[30]

The condition has been tested once on measured damage, and main text §5.3 rests the organophosphate recommendation on that test. Kretschmann *et al.* moved *Daphnia magna* to clean water 21 h into a diazinon exposure and measured acetylcholinesterase and immobilisation together;[18] the series used here is the one tabulated in the Supporting Information of a later paper by the same group.[S1] The enzyme fell for a further ten hours, then recovered, but stayed below the threshold fitted to this series in the later paper for at least the next thirteen hours. With that threshold and killing rate (25.6 pmol $g^{-1}$ and 0.0237 g $pmol^{-1}$ $h^{-1}$ in its Supporting Information; 25.4 and 0.025 in its Table 1), the cumulative hazard accrues $\Delta H = 0.60$–$1.49$ (across the duplicate enzyme measurements) over the 12.9 h from the enzyme minimum to the next sample, so survival should have fallen from 22.5% to 5–12%. Immobilisation was recorded continuously in the same two vials of twenty animals,[18, S1] so nine animals were unaffected at the start of that window, and the probability that none of them was then immobilised is $1 \times 10^{-6}$ to 0.006 across the duplicate measurements and the two parameter sets, below 0.01 in every case. With the values of the original study instead (23.0 pmol $g^{-1}$, 40% of control activity, and 0.0163 g $pmol^{-1}$ $h^{-1}$),[18] the deficit below that threshold is smaller and has closed in one duplicate by the next sample, the cumulative hazard accrues 0.12–0.47 and the probability is 0.014–0.35; the refit of both structures to the whole series below depends on neither set. The peak-based hazard predicts no change at any parameter value, and none was observed (`clean_water_test_diazoxon.py`).

Fitting both hazard structures to the whole series — the measured enzyme trajectory as input, two hazard parameters each — the running-maximum form reproduces the plateau in survival exactly and the cumulative form cannot: it lags the observed drop, predicting 47% survival where 22.5% was observed, and then keeps falling past the observations to 11%, because a positive deficit keeps accruing hazard for as long as it lasts. The likelihood favours the peak-based structure by 4.2 units (`clean_water_test_diazoxon.py`). Kretschmann *et al.* linked the enzyme to immobilisation through a cumulative hazard; refitted with the running maximum, the same data favour the peak-based form.

This is a fitted comparison of the kind §5.3 warns against, and it is informative here only because the damage state was measured rather than modelled and because the design includes the falling phase that §S14 requires; while damage is only rising, the control simulation of §S10 finds the two structures indistinguishable. That observation is one experiment on one compound and species.

The condition has also been applied as a test. The reduced peak-based structure is rejected for imidacloprid and satisfied again once the damage model is enlarged. On diazinon the condition is met in one treatment and not another. The fact that the answer tracks the fitted damage model rather than the hazard structure is the subject of a companion analysis, which reports the comparison in full.[7] What matters here is only that the condition is testable and has been tested.

##### S3.3 Which Structure Is the More Conservative at Matched Parameters

The guidance directs that where both reduced variants pass validation the more conservative be used, and its criteria are computed inside the validation experiment (main text §5.3). The ordering that rule observes inside the validation experiment is not the ordering that holds over the exposures the assessment covers, and that follows from §S1.1 and §S1.2 without further assumptions. That the ordering is not obvious has been said before: it cannot be known in advance which variant will predict the higher mortality under a given exposure pattern.[3] What follows says when it can; "more sensitive" throughout this section means predicting the lower survival, which is the more conservative choice.

Take constant exposure, so that damage rises monotonically to $D_{ss} > z$ and the running maximum equals the current level. Write the peak-based survival as $S_B(t) = 1 - F(D(t))$: for individual tolerance this is the definition, and a finite killing rate changes only which distribution $F$ stands for, not the form (§S3.1). The cumulative structure has $S_A(t) = \exp(-H_A)$ with $H_A$ as in §S1.1. In this subsection $\theta$ denotes the median of that log-logistic, as in Tables S1 and S1b and main text Figure 2, and $v_0 = k_r D_{ss}$ is the initial rate at which damage rises.

**In words.** The two structures kill in different ways, and that fixes which predicts more deaths at which time. The peak-based structure kills animals as damage climbs past their individual thresholds: the most sensitive die first, and once damage has levelled off no one else is reached, so its kill comes early and then stops. The cumulative structure kills only once damage is above its threshold, and then keeps killing for as long as damage stays there, so its kill starts later and never stops. Given enough time — provided damage settles above the cumulative threshold and some animals' tolerance lies above that level — the cumulative structure therefore predicts more deaths (part (i) below). Early on, the answer depends on how many very sensitive animals there are. If some animals have thresholds below the cumulative threshold $z$, the peak-based structure kills them before the cumulative one has started (the first case of part (ii)). With no cumulative threshold at all, it depends on the shape of the tolerance distribution: a broad one ($\beta < 2$) contains many very sensitive animals and the peak-based structure leads, while a narrow one ($\beta > 2$) contains few and the cumulative structure leads (the second case).

**Proposition S3 (the conservatism ordering is fixed at long exposure and not at short).** *Let $D(t) \to D_{ss} > z$ and let the tolerance distribution place some mass above $D_{ss}$, so that $F(D_{ss}) < 1$.*

*(i) There is a time beyond which $S_A(t) < S_B(t)$, at every parameter value: the cumulative structure is the more sensitive at long exposure.*

*(ii) At short exposure neither ordering holds generally. If $z > 0$ and the tolerance distribution places mass below $z$, as the log-logistic of §S3.1 does, the peak-based structure is the more sensitive on $(0, z/v_0)$, where $v_0 = k_r D_{ss}$; if $z = 0$ and the tolerance is log-logistic of shape $\beta$, the cumulative structure is the more sensitive when $\beta > 2$ and the peak-based one when $\beta < 2$.*

*Proof.* For (i), the integrand of $H_A$ tends to $b_w(D_{ss} - z) > 0$, so $H_A \to \infty$ and $S_A \to 0$, while $S_B \to 1 - F(D_{ss}) > 0$. For (ii), write $D \approx v_0 t$ at short times. Then $H_A$ is exactly zero on $(0, z/v_0)$ while $H_B = -\log(1 - F(v_0 t))$ is positive there, which gives the first case. When $z = 0$ the two accumulate as powers of time, $H_A \simeq \frac{1}{2} b_w v_0 t^2$ against $H_B \simeq (v_0 t/\theta)^\beta$, and the larger exponent is the smaller hazard as $t \to 0$. ▫

In the last step, "the larger exponent is the smaller hazard" means that for a short time $t$ a higher power is smaller — $t^3$ is less than $t^2$ when $t < 1$ — so the hazard that grows with the higher power starts more slowly.

Throughout this section the population is uniform apart from the peak-based tolerance: every organism shares the same killing rate and, under the cumulative structure, the same threshold. The restriction matters for part (i) only. If animals differ in killing rate but share a threshold, every one of them still accumulates hazard without limit, so the population dies out and part (i) stands (formally, $\mathbb{E}\left[e^{-b_w I}\right] \to 0$ whenever the accumulated exceedance $I = \int_0^t [D - z]^+ \, d\tau$ grows without bound). If they differ in threshold, those whose threshold lies above the steady damage never accumulate any, so $S_A(\infty) = P\,(z \geq D_{ss}) > 0$ and the long-exposure ordering is no longer guaranteed. Comparing the structures at heterogeneous parameters is a different question from the one here, as is comparing them after each has been fitted separately (below).

Table S1 runs both structures at constant exposure over a spread of parameter values, with the same damage dynamics given to each so that the only difference is how the hazard depends on them.

**Table S1. Which structure is the more sensitive, early and late.** Both structures are given $D(t) = D_{ss}\left(1 - e^{-k_r t}\right)$ and the peak-based one a log-logistic tolerance of shape $\beta$ and median $\theta$. Crossings are counted only where at least one structure has killed a measurable fraction.

| $k_r$ | $D_{ss}$ | $z$ | $b_w$ | $\beta$ | $\theta$ | early | late | crossings |
|---|---|---|---|---|---|---|---|---|
| 0.50 | 3.0 | 1.0 | 0.30 | 4.00 | 2.0 | peak-based | cumulative | 1 |
| 0.10 | 3.0 | 1.0 | 0.30 | 4.00 | 2.0 | peak-based | cumulative | 1 |
| 0.50 | 3.0 | 0.5 | 0.60 | 3.00 | 2.0 | peak-based | cumulative | 1 |
| 2.00 | 5.0 | 1.0 | 0.20 | 6.00 | 3.0 | peak-based | cumulative | 1 |
| 0.50 | 3.0 | 2.5 | 0.30 | 6.00 | 1.0 | peak-based | cumulative | 1 |
| 0.50 | 3.0 | 0.0 | 0.30 | 1.50 | 2.0 | peak-based | cumulative | 1 |
| 0.50 | 3.0 | 0.0 | 0.30 | 4.00 | 2.0 | **cumulative** | cumulative | **0** |
| 0.10 | 3.0 | 1.0 | 0.05 | 2.44 | 2.0 | peak-based | cumulative | 1 |
| 2.00 | 5.0 | 1.5 | 1.00 | 3.91 | 2.5 | peak-based | cumulative | 1 |

The late column of Table S1 is the same in every row, as Proposition S3(i) requires; the early column is not, and what sets it is exactly $F(z)$ — the fraction of the population whose tolerance lies below the cumulative structure's threshold, which over these rows runs from under one per cent to almost all of it. The one row with no crossing is the thresholdless, steep-tolerance case of Proposition S3(ii), where $t^2$ beats $t^\beta$ throughout. $F(z)$ is set by $z/\theta$ together with $\beta$, and neither is read off the data: the threshold is identified only jointly with the damage scale and the killing rate (§S9.1), and the shape is fixed rather than estimated (§S9.3).

Table S1b generalises the nine rows to a Monte Carlo sample of twenty thousand parameter draws, scored only where at least one structure has killed between five and ninety-five per cent so that the comparison is informative.

**Table S1b. The ordering over the parameter space, by exposure duration.** Draws are log-uniform over ranges drawn from the fitted values of §S13 and §S7. The $k_r$ range, 0.01 to 5 $d^{-1}$, covers fourteen of the seventeen recovery rates in Table S3 that are not at a fitting bound (three faster fits, 7.1–22.5 $d^{-1}$, lie above it; the other eleven of the twenty-eight sit at a bound and are not estimates, §S7.1); $b_w$ from 0.05 to 5, $\beta$ from 1.5 to 8, $z/\theta$ from 0 to 1.5, $D_{ss}/\theta$ from 0.5 to 8.

| duration (d) | draws scored | cumulative more sensitive | median survival gap |
|---|---|---|---|
| 1 | 8,858 | 9.7% | 0.354 |
| 2 | 10,154 | 17.0% | 0.304 |
| 4 | 11,128 | 25.4% | 0.267 |
| 7 | 11,536 | 33.0% | 0.243 |
| 14 | 11,497 | 44.4% | 0.234 |
| 28 | 10,989 | 56.0% | 0.228 |
| 96 | 10,201 | 68.8% | 0.259 |

| duration (d) | draws scored | cumulative more sensitive | median survival gap |
|---|---|---|---|
| 365 | 9,991 | 72.8% | 0.277 |

The share of draws in which the cumulative structure is the more conservative rises monotonically with duration and crosses one half between fourteen and twenty-eight days, which is Proposition S3(i) appearing as a frequency rather than as a limit: at any fixed duration some draws have not yet reached the crossing, and the share that have grows toward one with time. At the durations a standard acute test runs the ordering goes the other way in three cases out of four. The gap column of Table S1b is drawn as main text Figure 2b. The U shape of that column is the overtaking that Proposition S3 implies, seen as a curve. At short duration the peak-based structure has already killed the fraction $F(D)$ whose thresholds damage has passed, while the integral hazard has barely accumulated, so the gap is at its largest; as $H_A \propto t$ grows the cumulative kill catches up and the median gap falls; past the overtaking $S_A$ runs to zero while $S_B$ saturates at $1 - F(D_{ss})$, and the gap re-opens toward that asymptotic separation. The dip is not a pile-up of draws near their own sign change: on the same ensemble the share of draws with $|S_A - S_B| < 0.05$ stays between 4% and 6% from one to twenty-eight days and falls to 0.3% by a year (`conservatism_ordering_scan_gapshare.csv`, from the deposited scan) — the whole distribution shifts, few draws sit near zero at any one duration.

Main text Figure 2c resolves the four-day share by the two quantities Proposition S3(ii) names. That fraction runs from about 65% where the threshold sits well below the median tolerance to under 5% where it sits above it, and the tolerance shape matters only at the shallow end. Both quantities are the ones the calibration does not determine (§S9.1, §S9.3).

These frequencies are a property of the model pair, not of any fit: no experiment observes the two structures at the same parameters (§S7.3). Fitted separately to the fourteen compounds of §S7, neither fits better systematically — nine likelihood differences favour individual tolerance and five the cumulative form, a split a sign test does not separate from chance ($p = 0.42$, Table S3) — which is the distinction main text §5.5 draws. Brock *et al.* found the reduction in predicted risk under time-variable profiles larger under individual tolerance than under stochastic death;[S2] that is the opposite of the ordering above at short exposure, and for the same reason: two calibrated models are being compared, not two hazards.

Two things follow for practice. First, a rule that determines conservatism inside a short validation experiment determines it where the answer turns on those two quantities. Second, it applies that determination to exposures where the answer is fixed and, in eight of the nine rows, opposite. The rule is therefore not conservative in the sense it intends: it can select the structure that predicts less mortality over the period the assessment actually covers.

##### S3.4 When the Two Structures Can Be Told Apart: the Observation Window

While damage is still rising the two classes are not distinguishable, because the running maximum coincides with the current value and both hazards are monotone functions of the same trajectory. Given the same threshold, they also start killing at the same instant: the accumulated excess above a threshold cannot become positive before damage has passed that threshold, and the high-water mark passes it at that very moment (formally, $\int_0^t [D(s) - z]^+ \, ds > 0$ exactly when $\max_{s \le t} D(s) > z$), so the two structures place the onset of mortality at the identical event. They diverge only in the **post-equilibrium phase**, after damage has peaked and begun to fall, and that phase is reached only when the observation window exceeds $\max(3/k_e, 3/k_r)$.[29]

Three mechanisms keep standard tests inside the rising phase, where the two structures are equivalent: slow uptake, slow damage recovery, and saturating repair, each of which prolongs the rising phase. How far the recovery rate can differ between compounds is visible in an older experiment on *Chironomus riparius*, where two one-hour pulses separated by six hours in clean water reduced the effect of four carbamates from 58–81% to 10–12%, of parathion from 93% to 48%, and of malathion not at all. Acetylcholinesterase measured in the same animals recovered to control activity between carbaryl pulses.[S3] The ordering is graded rather than a split between chemical classes, and the companion paper compares it with the recovery times fitted here.[7] Six hours suffices for a carbamate, partly for one organophosphate and not at all for the other — the same graded recovery that leaves every organophosphate comparison of Table S2 short of the compound's own recovery time, while two carbamate comparisons clear it.

Designs with a long enough interval do exist for an anticholinesterase in another species: Naddy *et al.* gave *Daphnia magna* two chlorpyrifos pulses three to fourteen days apart.[S4, S5] They report mortality after each pulse: at 0.5 µg/L the first pulse killed 15–20% and the second most of the survivors at every interval, and the authors attribute part of the difference to a change of sensitivity with age between the pulses, which confounds the ratio with development. In the archive, where per-pulse survival is reported for an anticholinesterase, the interval is too short. The diazinon experiments of ref 20 give daily survival for two identical pulses separated by two, seven and fifteen days, and the ratios computed from them over matched spans are 0.57, 0.53 and 0.71 (`carryover_ratio_diazinon2010.py`). Every one is below unity, and every interval is shorter than diazinon's own 95% recovery time (167 days under the individual-tolerance fit of Table S3 and 20 days under the stochastic-death fit; in the three-compartment fits of §4.4, the 300-day bound for the cumulative structure (21 days without it, §S9.2), while the peak-based structure goes to the slow end of the recovery-rate range), so the counting rule of §S6.1.1 marks all three as non-informative before the values are read. Those values are what residual damage produces: a ratio below unity is what both structures give when damage has not reset, which is why only a ratio above unity discriminates (§S6.1.1). The position is general for the anticholinesterases — their recovery times exceed the intervals the published pulse designs leave — so the archive of §S6.1 remains the only source of informative ratios. This is why the classification cannot be settled by re-analysing existing acute data, and why §S10 finds the two structures indistinguishable even when the generating structure and the toxicokinetics are known exactly.

A further limit is that the window has to extend past the exposure. Cold and Forbes held *Gammarus pulex* at 100% survival through esfenvalerate pulses of up to 2 µg/L and then saw mortality rise, in places markedly, after transfer to clean water.[S6] A design that stops counting at the end of exposure cannot see that, and it is the behaviour that separates the two structures most sharply (main text §5.4).

Another limit is in the endpoint itself. The datasets on which delayed mortality has been demonstrated most clearly use immobilisation as a proxy for death.[15] Immobilisation is reversible, whereas the irreversibility on which both hazard structures rest (§2.1) concerns death; a model of either class applied to a reversible endpoint is outside the domain in which its structural argument holds. How far the two diverge has been measured. Andersen *et al.* gave *Daphnia magna* two identical pulses separated by 48 hours in clean media and scored mobile, immobile and dead separately. After the second dimethoate pulse only 5% were dead immediately and 85% two days later. After a single pirimicarb pulse every animal was immobile at the end of exposure and almost all had recovered a day later.[S7] Immobilisation therefore understates mortality for one compound and overstates it for the other, and which way it errs follows the same reactivation that graded the compounds above — pirimicarb is a carbamate and dimethoate an organophosphate. The consequences for repeated-pulse designs are taken up in the companion paper.[7] In the clean-water experiment of §S3.2 the unaffected fraction stayed constant in both duplicate vials from the enzyme minimum to the end of observation, 48 h later, so the endpoint showed no recovery there.

#### S4. Staged Activation and the $t^n$ Rise of Damage

The 1-compartment RED framework compresses the toxicokinetics and the damage dynamics into a single rate constant $k_r$. For compounds with multi-stage activation pathways (e.g., organophosphates: parent → oxon → phosphorylated acetylcholinesterase, AChE-P), this compression loses information about the shape of the damage trajectory, which determines the mortality pattern (main text §2.5). §S4.1 shows why each stage adds a power of time, and when that applies.

##### S4.1 Why Each Stage Adds a Power of Time, and When It Applies

For $n$ sequential first-order stages under constant input, the output of the $n$-th stage scales as $x_n(t) \propto t^n$ for $t \ll \min_j \left(1/k_j\right)$, where $k_j$ are the decay rates of intermediate stages. This is what successive integration gives. Stage 1, fed at a constant rate $u$, rises linearly, $x_1(t) \approx ut$. Stage 2 is fed by stage 1, so, with generic stage rates $k_1, k_2, \ldots$ — not the compartment rates of eq. 3 — $\frac{dx_2}{dt} \approx k_1 x_1 = k_1 ut$, and integrating once gives $x_2 \approx k_1 ut^2/2$. Each further stage integrates the one before it and so adds one power of $t$, which is why the $n$-th behaves as $t^n$.

For the organophosphate three-compartment model (parent → oxon → AChE-P), after the fastest stage equilibrates (parent TK, ~0.2 days), the effective two-stage process produces $D(t) \propto t^2$ over most of the 4-day test, which is why the three-stage pathway counts as $n = 2$. With the two sets of borrowed toxicokinetic constants of the dimethoate case study (§S8.1) the exponent fitted over the 4-day test is 1.78 and 2.43, against 0.98 for the reduced model (Figure S5a; by phase in Table S4).

The scaling requires the activation step to be slow relative to the stages around it, and for this compound class that has been measured rather than assumed. In *Daphnia magna* the oxidative desulfuration of diazinon to diazoxon proceeds at 0.014 $h^{-1}$ (0.34 $d^{-1}$) against a total elimination of the parent of 0.266 $h^{-1}$ (6.4 $d^{-1}$), so activation is rate-limiting by a factor of nineteen and carries only about 5% of the parent's turnover.[S1] Where activation is instead fast the intermediate equilibrates at once, the cascade collapses to a single stage, and damage rises linearly. The argument of §2.5 is therefore made for compounds requiring metabolic activation and not for direct-acting ones.

Not every multi-compartment model is a cascade, and the difference matters for two-pulse designs. A series of first-order stages is a convolution, and the damage input it produces is log-concave — its logarithm curves downward, so the input never decays fast and then lingers. A parallel decomposition into a positive mixture of exponentials, as two-compartment distribution kinetics gives, is not: it decays fast and then slowly. The two are easily confused because a cascade also yields a difference of exponentials, but only the cascade gives the $t^n$ rise above. The companion analysis shows log-concavity of the input to be exactly what keeps peak damage from rising as two pulses are moved further apart, for every non-negative repair law.[7]

#### S5. Saturating (Michaelis–Menten) Repair and Why These Data Cannot Identify It

The 2×2 classification (main text §2.2) makes the repair function an axis in its own right. The Michaelis–Menten form $R(D) = k_r D/(1 + D/K)$ is proportional repair at rate $k_r$ while $D \ll K$ and saturates at $R_{\max} = k_r K$ when $D \gg K$, so it interpolates between the two rows of Table 1: $K \to \infty$ at fixed $k_r$ gives proportional repair, and $K \to 0$ at fixed $R_{\max}$ gives constant-rate repair at $R_{\max}$ — the $k_{r,0}$ of main text §2.1 and §S1.3–S1.4. Every fit in this Supporting Information works in the proportional limit $K \to \infty$. This section states why.

The restriction is not a claim that these compounds lie in the proportional regime. It is a claim that these data cannot distinguish proportional repair from saturating repair. Giving the reduced peak-based model of the dimethoate case study (§4.4) Michaelis–Menten repair and refitting it at each $K$ from 10 to $10^9$ moves the calibration likelihood by at most 1.45 units from the proportional-repair fit, the largest change at the smallest $K$, and by less than 0.05 units above $K = 10^5$ (`michaelis_menten_repair.py`). Repair saturates at the highest test concentration once $K$ falls below about $2 \times 10^5$, where the maximum repair rate $R_{\max} = k_r K$ drops below the damage input, and at every concentration below about $10^3$; the likelihood does not separate either regime from proportional repair. The companion analysis of the clean-water test reaches the same conclusion for a receptor agonist by profiling $K$ directly, where the likelihood is flat over seven orders of magnitude.[7]

The same conclusion follows for the recovery rate itself. The peak-based calibration likelihood is identical to two decimals whether the recovery time is left free or held at 30, 100 or 300 days or longer (Tables S9 and S10), and its fitted $k_r$ goes to the slow end of the permitted range for both compounds. If the data cannot pin down the repair rate, they cannot pin down the shape of the repair function either.

Michaelis–Menten repair would become detectable in a design that met three conditions: a recovery rate intermediate enough for repair to act within the test duration (~0.1–1 $d^{-1}$); concentrations bracketing the point at which repair is overwhelmed, which for the three-compartment model is the external concentration at which the steady-state damage input $k_3 A$ equals $R_{\max}$, $C^*_{\text{ext}} = k_r K\, k_{\text{out}} k_{2,\text{out}} / (k_3 k_2 k_{\text{in}})$; and time-resolved observations spanning the transition. None of the designs available here meets them.

### Part II. Model Structure and Parameter Estimation

*How this Part supports the main text.* §S6 holds the first two results of the main text, the exclusion and the ceiling (§4.1), together with the evidence that constrains without discriminating. §S7–S9 hold the fourth result, the fitted comparisons of §4.4 and what limits them, with the identifiability analysis behind §5.3. §S10 is the premise of §4.3, the control simulation; §S11 asks whether a change of data could substitute for a change of model.

#### S6. Evidence Independent of Fitting: the Carry-Over Ratio, Its Ceiling and the Lethal Residue

This section holds the paper's central result, that measured survival counts exclude the cumulative structure without any model being fitted. As in §S1 and §S3, each result is stated first in words, and the formal statements and proofs can be skipped. The argument runs in four steps.

1. **Measure one number.** Give animals two identical pulses and compare how the survivors of the first fare in the second with how everyone fared in the first. That is the carry-over ratio $\mathcal{R}$ (§S6.1.1).
2. **The cumulative structure allows only one answer.** If damage has fully reset between the pulses, it predicts $\mathcal{R} = 1$ exactly, at every parameter value; if it has not, it predicts less than 1. A measured ratio above 1 is therefore impossible under it (Proposition S4, §S6.1.2).
3. **Close the one way out.** A ratio above 1 could still arise if animals differed in how quickly damage kills them, because the first pulse would then remove the fragile ones. The ordinary concentration–response data limit how large that difference can be, and the limit is too small to produce five of the six measured ratios that can be checked this way (§S6.1.3).

4. **Check the alternative's own prediction.** The peak-based structure predicts not just a ratio above 1 but a specific ceiling, $1/S_1$, approached as recovery becomes complete; the measured ratios approach it (Proposition S5, §S6.2).

Seven measured comparisons from four compounds have 95% confidence intervals lying wholly above 1 (§S6.1.4). §S6.3 adds a constraint from the lethal residue, and §§S6.4–S6.6 three considerations consistent with the peak-based structure that do not discriminate by themselves. The two propositions are stated here, beside their test, rather than in Part I, because both are predictions of one two-pulse design.

##### S6.1 The Carry-Over Ratio Excludes the Cumulative Structure

**S6.1.1 The Ratio and When It Can Be Read** The carry-over ratio quantifies whether surviving a first pulse makes an organism more vulnerable or more resistant. Writing $S_1$ for the survival fraction over the first pulse and $S_{2|1}$ for the conditional survival of its survivors over an identical second pulse, both corrected for control mortality over matching windows,

$$\mathcal{R} = \frac{S_{2|1}}{S_1},$$

so $\mathcal{R} < 1$ means survivors weakened by what they carry over and $\mathcal{R} > 1$ means survivors more resistant than the population they came from.

For example, suppose a first pulse kills half the animals ($S_1 = 0.5$). If half of the survivors then die in an identical second pulse ($S_{2|1} = 0.5$), $\mathcal{R} = 1$: surviving the first pulse said nothing about an animal. If only one in ten of them dies ($S_{2|1} = 0.9$), $\mathcal{R} = 1.8$: the first pulse picked out the tougher animals. The largest possible value is $1/S_1 = 2$, reached when every survivor of the first pulse survives the second.

The ratio is what a two-pulse experiment measures, and the two structures predict different values for it (§S6.1.2, §S6.2.1). That animals given long enough between pulses return close to their original resistance does not have to be assumed: it has been shown separately in an amphipod for a metal and an uncoupler,[S8] and in a midge for four carbamates, with the enzyme activity recovering alongside the whole-animal response.[S3] Whether the measurement can be read at all depends on the interval. Both structures give $\mathcal{R} < 1$ when damage has not reset, because the second pulse then starts from an elevated level, so a ratio below unity is uninformative unless the interval exceeds the compound's own recovery time. Under the individual-tolerance fits that time runs from under a day for the baseline narcotics to 167 days for diazinon and 258 for 2,4,5-trichlorophenol (Table S3), so fifteen days is ample for a narcotic and insufficient for diazinon. A ratio above unity refutes Class A whatever the interval.

**S6.1.2 Memorylessness of the Cumulative Hazard** The first of the two propositions concerns what the integral hazard can represent at all.

**In words.** Under the cumulative structure as GUTS-SD implements it, every animal has the same parameters, so whether it dies in a pulse is a matter of chance alone — like whether a coin comes up heads. Surviving the first pulse therefore says nothing about the animal: it is no tougher than one that never saw the pulse. If damage has fully reset, the second pulse is an exact repeat of the first, the same fraction of the survivors dies in it as of all the animals in the first, and the ratio is exactly 1, whatever the threshold, killing rate or exposure shape.

**Proposition S4 (memorylessness of the cumulative hazard).** *Let $D(t)$ be a damage trajectory driven by two identical exposure pulses separated by an interval long enough that $D$ returns to zero between them, in a population whose members share the same parameters.*

*Then under Class A the carry-over ratio $\mathcal{R} = S_{2|1}/S_1$ equals unity for every value of $b_w$, $z$, $k_a$ and $k_r$.*

*Proof.* Complete recovery makes the second pulse an exact replicate of the first, so each contributes the same increment to the cumulative hazard over its window of length $\Delta t$ (the pulse and its recovery),

$$\Delta H = \int_0^{\Delta t} b_w \, [D(\tau) - z]^+ \, d\tau,$$

independently of what preceded it. Hence $H\,(2\text{ pulses}) = 2\,\Delta H$ and $S\,(2\text{ pulses}) = e^{-2\Delta H} = S\,(1\text{ pulse})^2$, so $\mathcal{R} = 1$. □

**If damage has not reset.** The proposition also fixes what happens when recovery is incomplete. Residual damage makes the second pulse's trajectory everywhere at least as large as the first's, so the cumulative hazard of two pulses is at least twice that of one and $\mathcal{R} \le 1$ under Class A at every parameter value, with equality exactly at complete recovery. A ratio below unity is therefore consistent with the cumulative structure and excludes nothing. That asymmetry is why a ratio at or below unity has to be read together with the recovery interval (§S14), whereas one above unity needs no such check.

**S6.1.3 What the Theorem Does Not Exclude, and How Far the Data Bound It** What makes the result decisive is that it holds for every value of the threshold, the killing rate and the exposure shape, provided the animals share them. Class A is memoryless between pulses whenever recovery is complete, at every point in its parameter space, so a single measurement of $\mathcal{R} > 1$ excludes the cumulative hazard as GUTS-SD implements it, with no fitting involved.

It does not exclude every model with a cumulative hazard. The variation the theorem does not cover is in the killing rate, not in the threshold. Distributing the threshold turns the cumulative structure into GUTS proper, which in the limit becomes individual tolerance — the family Proposition S1 identifies with the peak-based structure — so it does not rescue the cumulative structure; once that route is taken, a killing rate that varies between individuals is what remains. A killing rate that varies between individuals lets the survivors of pulse 1 be enriched in the less susceptible and produces a ratio above unity without any running maximum; how far the data allow that variation is quantified below and in Figure S2. The lethal residue of §S6.3 does not separate a varying killing rate from a running maximum either: the predicted fall of the lethal residue with time (§S6.3) survives that variation, but so does Class A's way of matching the observation, since a threshold placed at the observation flattens the predicted fall whatever the variance.

**In words.** If animals differ in how fast damage kills them — some fragile, some robust — the first pulse removes the fragile ones, and the survivors are tougher, which gives a ratio above 1 with no running maximum at all. But that variation would leave two fingerprints in data already published alongside each pulse experiment. It would make the concentration–response curve shallower, and it would make the death rate under constant exposure slow down over time, as the fragile animals are used up. Both were checked. The curves allow only a small variation, too small to produce the largest measured ratio in any of the three compounds that can be checked this way, and the death rates do not slow down; if anything they speed up.

What separates the two is the concentration–response curve, which that variation flattens as surely as it lifts the ratio, so the LC50 series published alongside each pulse experiment bounds that variation. Fitting a gamma-distributed multiplier on the killing rate (a frailty — a factor, different for each animal, by which its killing rate is multiplied; $\nu$ measures how much the factor varies, and $\nu = 0$ means that all animals are alike) of variance $\nu$ to those series returns $\hat{v} = 0$ for all six compounds that carry one, with 95% profile limits between 0.17 and 1.14 (`frailty_variance_bound.py`). Three of the four compounds behind the seven comparisons whose interval excludes unity have such a series. Holding $v$ at its upper limit and matching the first-pulse survival of the treatment concerned, the largest ratio the population could produce is 2.04, 1.99 and 1.55 for 4,6-dinitro-*o*-cresol treatments C, A and B, against 4.42, 2.18 and 2.87 observed; 1.26 and 1.04 for the two 2,4-dichloroaniline treatments against 2.36 and 1.45; and 1.01 for 2,4-dichlorophenol against 1.21. In five of the six the frailty-limited value lies below the ratio's 95% interval; for 4,6-dinitro-*o*-cresol A it lies inside it (1.08–4.41). Between-individual variation of the size these curves allow therefore accounts for between a twentieth and 84% of each ratio's excess over unity, under a third in five of the six (`frailty_variance_bound.py`; Figure S2). The fourth compound, 1,2,3-trichlorobenzene, carries no concentration series and is not tested this way.

Figure S2 quantifies that comparison. With killing rates varying between individuals, first-pulse survival and the carry-over ratio are set by the same variance, so at the measured first-pulse survival the ratio is a function of that variance alone: panel (a) draws it for four compounds at their largest measured ratio, marks the variance the compound's own concentration series allows, and shows the ratio actually measured. The two never meet — 4,6-dinitro-*o*-cresol would need a variance near 19 where its series allows 0.56, and 2,4-dichloroaniline near 1.4 where its series allows 0.17 (`figureS2_frailty_envelope.py`). Panel (b) repeats the curve for lognormal and inverse-Gaussian frailties, which reach lower ratios than the gamma at the same variance, so the shortfall is not an artefact of that choice.

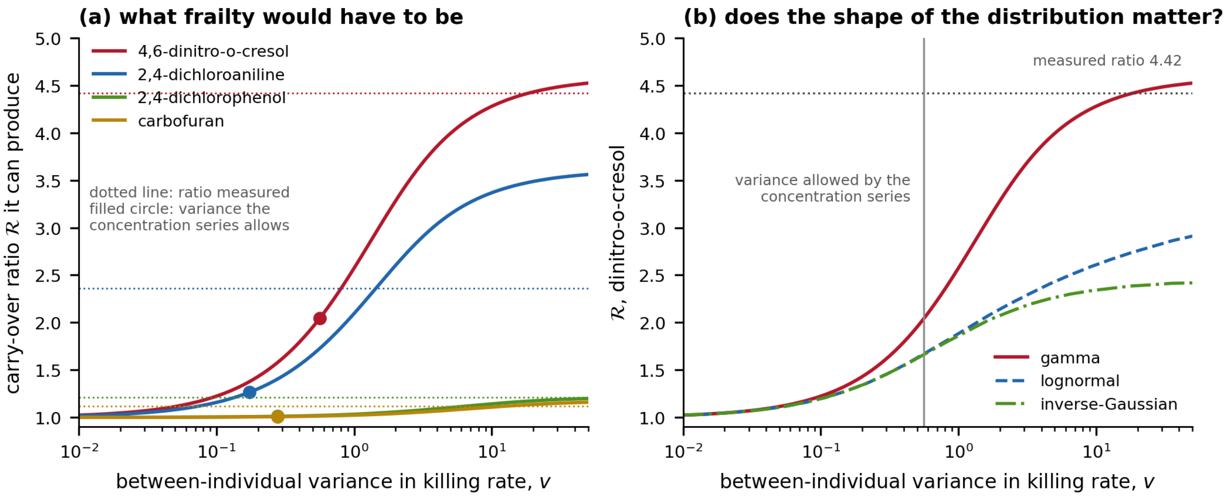


**Figure S2. How much between-individual variation the measured ratios would need.** Under a cumulative hazard whose killing rate varies between individuals, first-pulse survival $S_1$ and the carry-over ratio are both determined by the Laplace transform of the frailty distribution (the average, over animals, of each one's chance of surviving its own share of the hazard), so at the measured $S_1$ the ratio depends only on the variance $v$. **(a)** The ratio 4,6-dinitro-*o*-cresol, 2,4-dichloroaniline, 2,4-dichlorophenol and carbofuran could produce at the first-pulse survival of their largest measured ratio, against $v$ (curves); the variance its own concentration–response series allows (filled circles, §S6.1.3); and the ratio actually measured (dotted lines). **(b)** The same curve for 4,6-dinitro-*o*-cresol under three frailty families, with the allowed variance marked. The gamma curve is closed form; the other two families use $2 \times 10^5$ draws at a fixed seed (§S15).

The same series carry a second reading that assumes no particular frailty distribution. In plain terms: if the animals are alike, the logarithm of survival falls along a straight line once damage has built up — the same share of the survivors dies every day; if they differ, the line bends, because the fragile die first and those left are harder to kill. Formally, write $\Lambda(t) = -\log S(t)$ for the population cumulative hazard. If the animals are alike, $\Lambda$ grows linearly once damage has reached steady state; if the killing rate varies between them, the susceptible die first and $\Lambda$ bends over, because population survival is the Laplace transform of the frailty density,[S9] and the logarithm of a Laplace transform is convex for every density, so $\Lambda$ is concave. The curvature is the variance itself: for a gamma frailty $\Lambda'' = -\nu r^2/(1+\nu r t)^2$, so $\nu = -\Lambda''(0)/\Lambda'(0)^2$. Fitting linear and concave growth to the time course at each concentration, and using only the days after damage has built up, the linear form is preferred for all six compounds — by 6.4 likelihood units for carbofuran and 2.2 for 4,6-dinitro-*o*-cresol (`frailty_curve_checks.py`). At the concentrations that resolve the shape best, $\Lambda$ rises by more each day rather than less. This reading uses no concentration axis, so it is untouched both by measurement error in the exposure and by the threshold that lets a steep curve coexist with a large variance. The same series show no sign of saturation: in both uncouplers four-day survival keeps falling with concentration up to the highest tested, to 5% for 4,6-dinitro-*o*-cresol and 30% for 2,4,5-trichlorophenol, where a saturating receptor pool would hold it level (`frailty_curve_checks.py`).

Three limits attach to the readings above. One of the four compounds whose ratios exclude unity has no concentration series. The bound on $\nu$ is derived for a gamma frailty, though the curvature reading is not; Figure S2b shows that two other families leave the shortfall in place, but a bound that assumes no particular frailty distribution would remove this limit altogether, and deriving one is left to further work. And the LC50 series and the pulse experiment are separate cohorts, so the two are assumed comparable. The design conditions of §S14 avoid all three, since both curves would come from the same animals.

Class B is equally refutable in the same experiment, by a condition of its own: survival must be flat through any interval in which damage is falling, so a death during clean water excludes it (§S3.2). That condition has been tested and is not always met.[15,27,28]

**S6.1.4 What the Published Counts Give** Repeated-pulse survival counts and exposure schedules are available for the fourteen compounds of §S7.[2] Twenty-one two-pulse comparisons, from eleven compounds, have a first pulse that killed between 5 and 95% of the animals; carbofuran A and malathion B and C fail that filter. Fifteen of the twenty-one are informative: a ratio above unity counts whatever the interval, and one at or below unity counts only where the interval exceeded the compound's own damage recovery time, since below that both structures predict $\mathcal{R} < 1$. The six uninformative comparisons are reported and marked.

**Table S2. Carry-over ratio recomputed from the raw survival counts.** Control-corrected, matched windows; where the published survival record of the first window runs past the start of the second pulse, the first window is ended at the second pulse. $t_{95} = \ln 20/k_r$ is the compound's damage recovery time under the individual-tolerance fit (Table S3), the slower of the two fitted recoveries and so the stricter test; † $k_r$ at the lower fitting bound. Bold marks a ratio above unity; a tick marks the comparisons whose 95% confidence interval lies wholly above unity. Carbaryl treatments are labelled as in the published data file. The last three rows are the 2010 diazinon pulse experiment (ref 20; `carryover_ratio_diazinon2010.py`): every interval is shorter than the compound's own $t_{95}$ under either fit (167 d individual-tolerance, 20 d stochastic-death), so none of the three is informative. The interval condition applies only to $\mathcal{R} \leq 1$: a ratio above unity excludes GUTS-SD whatever the interval, since residual damage can only push the ratio down (Proposition S4). Under the stochastic-death recovery times instead (`carryover_ratio.py --sd`), nineteen comparisons from ten compounds are informative, fourteen at or above unity and the same seven excluding it; three more become informative and lie below unity with intervals excluding it, 2,4,5-trichlorophenol A (0.48, 0.31–0.75), malathion A (0.81, 0.66–0.99) and chlorpyrifos III (0.52, 0.37–0.73). Once damage has reset, neither structure predicts a ratio below unity (Propositions S4 and S5), so these three rows indicate that damage had not reset, and that the stochastic-death recovery times are too fast for these three compounds (for 2,4,5-trichlorophenol that fit sits at the upper bound of $k_r$, with $z = 0$).

| Compound | Mode of action | Trt | Interval (d) | $t_{95}$ (d) | $S_1$ | $S_{2\|1}$ | $\mathcal{R}$ | 95% CI | Excludes unity |
|---|---|---|---|---|---|---|---|---|---|
| 1,2,3-Trichlorobenzene | baseline | A | 4 | 0.4 | 0.656 | 0.876 | **1.34** | 1.05–1.70 | ✓ |
| 1,2,3-Trichlorobenzene | baseline | B | 7 | 0.4 | 0.524 | 0.581 | **1.11** | 0.73–1.68 | |
| 1,2,3-Trichlorobenzene | baseline | C | 11 | 0.4 | 0.640 | 0.828 | **1.29** | 0.98–1.71 | |
| 2,4-Dichloroaniline | baseline | A | 4 | 0.003 | 0.415 | 0.622 | **1.50** | 0.98–2.30 | |
| 2,4-Dichloroaniline | baseline | B | 7 | 0.003 | 0.277 | 0.653 | **2.36** | 1.36–4.08 | ✓ |
| 2,4-Dichloroaniline | baseline | C | 11 | 0.003 | 0.600 | 0.871 | **1.45** | 1.13–1.87 | ✓ |
| 2,4-Dichlorophenol | baseline | B | 7 | 0.2 | 0.919 | 0.795 | 0.86 | 0.72–1.04 | |
| 2,4-Dichlorophenol | baseline | C | 11 | 0.2 | 0.823 | 0.995 | **1.21** | 1.04–1.40 | ✓ |
| 2,4,5-Trichlorophenol | uncoupler | B | 5 | 258 | 0.608 | 0.643 | **1.06** | 0.74–1.52 | |
| 4,6-Dinitro-o-cresol | uncoupler | A | 2 | 2.7 | 0.226 | 0.493 | **2.18** | 1.08–4.41 | ✓ |
| 4,6-Dinitro-o-cresol | uncoupler | B | 4 | 2.7 | 0.327 | 0.940 | **2.87** | 1.96–4.21 | ✓ |
| 4,6-Dinitro-o-cresol | uncoupler | C | 8 | 2.7 | 0.218 | 0.965 | **4.42** | 2.71–7.22 | ✓ |
| Carbofuran | carbamate | B | 5 | 0.8 | 0.851 | 0.950 | **1.12** | 0.93–1.34 | |
| Aldicarb | carbamate | B | 5 | 4.1 | 0.851 | 0.882 | **1.04** | 0.85–1.26 | |
| Sea-nine | reactive | A | 6 | ≥ 30,000† | 0.718 | 0.828 | **1.15** | 0.93–1.44 | |
| *$\mathcal{R} \leq 1$ with interval below $t_{95}$ — uninformative* | | | | | | | | | |
| 2,4,5-Trichlorophenol | uncoupler | A | 2 | 258 | 0.741 | 0.357 | 0.48 | 0.31–0.75 | |
| Aldicarb | carbamate | A | 2 | 4.1 | 0.914 | 0.848 | 0.93 | 0.78–1.11 | |
| Malathion | OP | A | 5.2 | 36 | 0.938 | 0.761 | 0.81 | 0.66–0.99 | |
| Carbaryl | carbamate | 7TE-B | 9 | 178 | 0.787 | 0.131 | 0.17 | 0.05–0.62 | |
| Carbaryl | carbamate | 9TE-A | 9 | 178 | 0.772 | 0.703 | 0.91 | 0.68–1.21 | |
| Chlorpyrifos | OP | III | 14 | ≥ 30,000† | 0.939 | 0.487 | 0.52 | 0.37–0.73 | |
| Diazinon | OP | A | 2 | 167 | 0.924 | 0.527 | 0.57 | 0.42–0.77 | |
| Diazinon | OP | B | 7 | 167 | 0.876 | 0.462 | 0.53 | 0.34–0.82 | |
| Diazinon | OP | C | 15 | 167 | 0.891 | 0.632 | 0.71 | 0.52–0.96 | |

Among the fifteen informative comparisons, from eight compounds and four modes of action, the ratio is at or above unity in fourteen, and in seven its confidence interval lies wholly above unity; the median is 1.29 and the range 0.86–4.42. The intervals are delta-method intervals formed on the log scale from the two survival proportions and the numbers alive at the end of each window (§S12.3); the ratio is bounded above by $1/S_1$, so the intervals are asymmetric and are reported as computed rather than symmetrised. Correcting for the fifteen tests by the Benjamini–Hochberg procedure (two-sided) leaves six of the seven below $q = 0.05$; the seventh, 4,6-dinitro-*o*-cresol A, has $q = 0.063$. Treatments within a compound share a cohort, so the compound is the independent unit: taking the largest ratio in each of the eight gives 1.34, 2.36, 1.21, 1.06, 4.42, 1.12, 1.04 and 1.15, at or above unity in all eight and excluding unity in four — three baseline narcotics and 4,6-dinitro-*o*-cresol. Because $\mathcal{R} = 1$ is a theorem of the cumulative structure rather than a fitted outcome, each comparison whose interval lies above unity is on its own sufficient to exclude GUTS-SD for the compound concerned. The anticholinesterases give two informative ratios, carbofuran B and aldicarb B, both above unity with intervals spanning it. The one informative comparison below unity, 2,4-dichlorophenol B, had a first pulse that killed only 8%, leaving almost no survivor selection to detect. The computation is reproduced by `carryover_ratio.py`, which reads the published data file directly.

##### S6.2 The Ratio Against Its Ceiling

**S6.2.1 Attainment of the Ceiling In words.** If damage resets fully, the second pulse pushes damage only as high as the first did. Under the peak-based hazard the risk is set by the high-water mark, which the second pulse does not raise, so every animal that survived the first pulse survives the second, apart from background deaths. The survivors of the first pulse are exactly the animals whose effective threshold — their own threshold plus the random margin of §S3.1 — lay above its peak, and they have nothing more to fear from a repeat. The ratio then reaches its largest possible value, $1/S_1$. For the 4,6-dinitro-*o*-cresol treatment C, $S_1 = 0.218$ gives a ceiling of 4.59, and the measured ratio is 4.42.

**Proposition S5 (attainment of the ceiling).** *Let two identical pulses be separated by an interval long enough that damage returns to its pre-exposure level. Then under Class B the carry-over ratio equals* $1/S_1$ *exactly, for every killing rate and every tolerance density.*

*Proof.* The second pulse reproduces the trajectory of the first, so the running maximum at the end of the second equals its value at the end of the first. The hazard $k\,[M(t) - \theta]^+$ is a function of $M$ alone, so it does not advance during the second pulse, every animal that survived the first survives the second, and $S_{2|1} = 1$. □

The value $1/S_1$ is a ceiling for any model at all, since $S_{2|1} \le 1$; the content of Proposition S5 is that Class B attains it rather than approaching it. Background mortality lowers $S_{2|1}$ by $e^{-h_b \Delta t}$, which the control correction removes, and an interval too short for damage to reset lets the maximum advance, so the ratio falls below the ceiling.

What the peak-based structure predicts depends on the recovery rate, and the sign of $\mathcal{R} - 1$ turns on whether damage has reset between the pulses.

When recovery is incomplete, pulse 2 starts from elevated damage and reaches a higher peak than pulse 1, ratcheting the running maximum upward. Pulse-1 survivors then face increased hazard from pulse 2 — their conditional survival is lower than that of naive organisms, giving $\mathcal{R} < 1$, the sign the cumulative structure would also give.

$\mathcal{R}(k_r)$ therefore runs from below unity when nothing recovers to above it when damage resets. In the simulation of Figure S3b, at fixed exposure, it crosses unity once, at $k_r^* = 0.17\ \mathrm{d}^{-1}$, approaches the ceiling $1/S_1$ of Proposition S5 near $k_r = 1\ \mathrm{d}^{-1}$, and returns to unity at faster recovery, where the first pulse no longer kills anyone (`figureS3_carryover_vs_repair.py`). The crossing is a property of the experiment as much as of the chemistry, so a compound sitting near it returns $\mathcal{R} \approx 1$ under either structure; §S3.4 and §S14 give the design conditions that keep a test away from it.

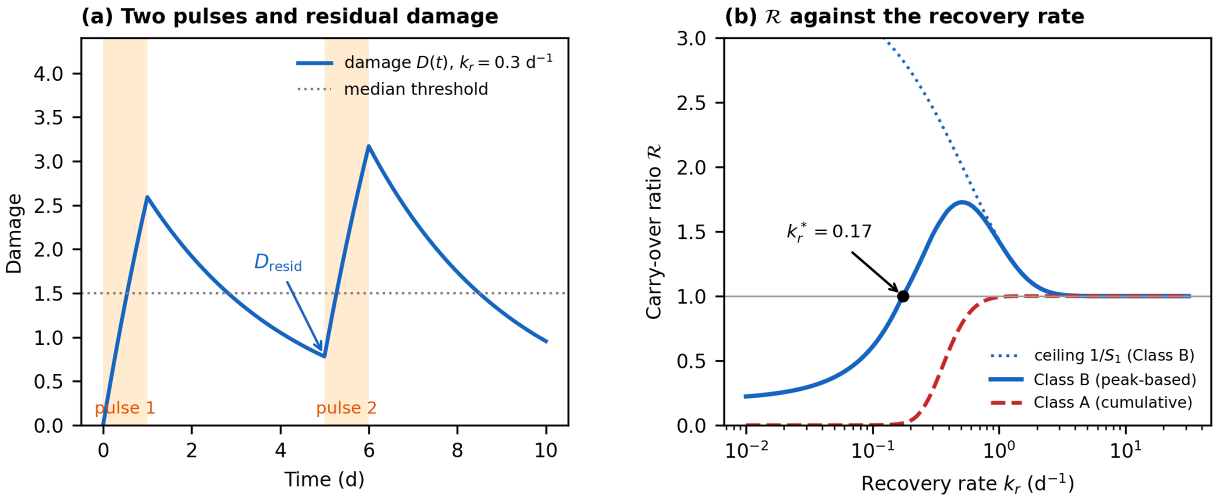


**Figure S3. The carry-over ratio against the damage recovery rate.** Two one-day pulses four days apart; damage $dD/dt = C - k_r D$ with $C = 3$; log-logistic tolerance with median 1.5 and $\beta = 5$; killing rates 1. (a) Damage at $k_r = 0.3$ d$^{-1}$; the residual damage at the second pulse is what lowers $\mathcal{R}$ when recovery is slow. (b) $\mathcal{R}$ against $k_r$ over matched five-day windows: the peak-based structure (blue) crosses unity at $k_r^*$, approaches its ceiling $1/S_1$ (dotted) and returns to unity once the first pulse kills no one; the cumulative structure with a shared threshold (red dashed) stays below unity and reaches it only when damage resets between the pulses, which is Proposition S4 (§S6.1.2).

**S6.2.2 The Ceiling in the Published Counts** Excluding GUTS-SD is not the same as accepting the peak-based structure, and the ratio alone does not accept it: a ratio above unity is what that structure predicts, not what it proves. The peak-based structure does make a further prediction the cumulative structure cannot, stated as Proposition S5 above.

Class A cannot exceed unity at all, while the peak-based structure approaches the ceiling $1/S_1$ as recovery becomes complete. Figure S3b draws that approach at fixed parameters; in the measured comparisons the ceiling is each treatment's own $1/S_1$, and main text Figure 1b plots every ratio as a fraction of it. The measured 4.42 sits at 96% of its ceiling — the second pulse killed 3.5% of the survivors of the first, which is what near-complete recovery looks like — and Table S2 can be read against the ceiling directly, since $\mathcal{R}S_1$ is simply the tabulated $S_{2|1}$.

That prediction can be checked: the fraction of the ceiling reached should rise with the interval measured against the compound's own recovery time. Six compounds were tested at two or more intervals. In five of them — 2,4-dichloroaniline, 2,4-dichlorophenol, 2,4,5-trichlorophenol, 4,6-dinitro-*o*-cresol and aldicarb — the fraction of the ceiling reached rises with the interval throughout; in 1,2,3-trichlorobenzene it does not (0.88, 0.58, 0.83). Ten of twelve treatment pairs within a compound rise with the interval (sign test $p = 0.019$; main text Figure 1b, `carryover_ceiling_approach.py`); the pairs share treatments, so the count by compound, five of six, is the more conservative reading. Every treatment that clears the 5–95% filter enters this count, including those Table S2 marks uninformative, because the prediction concerns the approach to the ceiling rather than the sign of $\mathcal{R} - 1$; compounds tested at a single interval are not tested, and pairs with equal intervals are excluded. The pooled rank correlation across compounds is weaker (Spearman +0.32, $p = 0.17$) because first-pulse survival, which sets the ceiling, varies more between compounds than the interval does. One alternative is consistent with both the ratio above unity and its approach to the ceiling: a defence induced by the first pulse would leave survivors made more tolerant by the first pulse rather than selected for tolerance by it. Induction develops and then wanes, so it would make the ratio peak at some interval rather than climb, which is the pattern in none of the five compounds that rise throughout and in 1,2,3-trichlorobenzene only as a dip, not a peak.

##### S6.3 What the Lethal Residue Constrains

Narcotics show permanent survival below a critical body residue and death above it.[S10] Whether the residue at death stays constant as exposure lengthens differs between studies. For chlorinated benzenes in *Poecilia reticulata* it did (below).[S11] For polycyclic aromatic hydrocarbons in *Hyalella azteca* it fell with exposure time, and the authors of the damage assessment model concluded that the toxicity threshold cannot be defined as a body residue.[16] That fall does not separate the structures either: under Class B the lethal residue falls for as long as damage is still rising at a steady body residue, which slow repair prolongs, and it becomes constant only once damage equilibrates. The argument of this section therefore applies where the lethal residue has been measured to be constant.

This section makes an argument about parameter freedom rather than a refutation. Class A can reproduce the observation, but only from a corner of its parameter space that carries a further prediction of its own; Class B reproduces it from most of its own. The carry-over ratio of §S6.1 is the result that excludes rather than constrains.

**What each structure is free to do.** Both structures can reproduce a single lethal residue at one time by placing their threshold below it. They separate on whether that residue moves. Under Class B the hazard is fixed once damage equilibrates, so the lethal residue is constant thereafter; under Class A it keeps accumulating, so the lethal residue — the body residue at which half the animals die within time $t$ — continues to fall as $z + \ln 2/(b_w t)$, approaching $z$ only as the exposure grows very long. The same calculation delivers Haber's law[14] from Class B through toxicokinetics alone, with no additional assumption (§S1.4).

**The measurement.** Van Hoogen and Opperhuizen exposed *Poecilia reticulata* to a series of chlorinated benzenes until death and found the lethal body burden between 2.0 and 2.5 mmol kg$^{-1}$ and **independent of exposure time and concentration**.[S11] Class B predicts no change over that window; Class A predicts a fall whose size depends on where the threshold sits. Scanning the $(b_w, z)$ plane, the combinations that keep the predicted residue inside 2.0–2.5 mmol kg$^{-1}$ throughout are not empty, but they require $z \geq 1.92$ mmol kg$^{-1}$ and $b_w \geq 0.6$ — a no-effect threshold at 77–96% of the measured lethal residue (`lethal_residue_scan.py`). The claim is therefore two-step: Class A can match the observation only by placing its threshold at the observation, and a threshold there predicts that nothing at all dies below 1.92 mmol kg$^{-1}$ however long the exposure. That claim is stronger than the data support, and the cumulative structure gains nothing by it, since Class B reproduces the same constancy with a threshold anywhere below the residue.

That the lethal residue does not move with exposure time is a property of baseline narcosis, not of every compound called a narcotic. For 2,3,4,5-tetrachloroaniline the lethal body burden falls from 1.8 to 0.7 mmol kg$^{-1}$ between fish dying within two days and later, which the reviewers of that work attribute to a shift away from narcosis toward a specific mode of action.[S12] The argument here is therefore made for baseline narcotics, and the recommendation is restricted accordingly.

**Confirmation from the fitted parameters.** The fits bear this out in a different way. Across the fourteen compounds of §S7, Class A drives the damage recovery rate to its upper fitting bound for six of them, including all three baseline narcotics, and a parameter at a bound is not an estimate (§S7). Raising $k_r$ does not in any case escape the unbounded growth of the cumulative hazard: it lowers $D_{ss} = k_a C_b / k_r$, but $D_{ss}$ is still constant in time under constant exposure, so the integrand stays positive and the integral still grows without limit. That bound is where the optimiser goes when the structure cannot represent the observation, not a property of the compound.

##### S6.4 Acute Death Mechanisms Are Threshold Transitions

The three death mechanisms that dominate acute aquatic toxicity are all threshold transitions rather than accumulating processes. Mitochondrial permeability transition opens only once the pore's conductance threshold is crossed, and a sub-threshold load does not open it partially. What an exposure has to achieve is reaching that threshold rather than staying near it, and openings themselves may be transient or long-lasting.[8] Phosphorylation of acetylcholinesterase proceeds to an aged, irreversibly inhibited adduct once the oxon has bound, a two-step irreversible mechanism demonstrated in vitro for diazoxon,[18] and inhibition is used as an exposure indicator on that basis.[9] Oxidative collapse follows depletion of the reducing pool past the point where it can be regenerated.[10] In each case the organism's fate is set by whether damage *reached* a critical level, not by how long it remained above one. The distinction between a threshold and an accumulating quantity is the one Ashauer and Brown draw when they place the existing models in a common framework.[11] A structural beam fails when the applied load exceeds its capacity, whether the load is applied for a second or an hour; the integral of load over time is not the quantity that breaks it. The peak-based hazard encodes exactly this and the cumulative hazard encodes the opposite.

The three mechanisms above are examples of the class, not its definition. What the peak-based structure requires is only that death be the crossing of a per-individual threshold, and that the crossing be irreversible — which it is, since an organism does not recover from being dead, so the running maximum follows whether or not the lesion itself can be undone. Sub-threshold exposure must then leave no separate memory, and it is that condition, rather than the chemistry, that the cumulative structure denies (§S6.5). A killing rate that varies between individuals satisfies the first two requirements as well, which is why the ratio alone does not separate it from the running maximum (§S6.1). The scope of the recommendation in the main text is narrower than this because it is set by where the carry-over ratio has been measured and where the two further decisions of §5.4 can be settled, not by where the structure is defensible.

##### S6.5 Independent Killing Events Give a Constant Death Rate, and What Gives a Rising One

Set $k_r = 0$, so that damage accumulates without repair and rises linearly under constant exposure. Class B then gives a constant death rate, because the running maximum advances at a constant rate once the threshold is passed; Class A gives a rate increasing linearly in time, because the integrand grows. For a lesion defined by a threshold on a large pool of targets, independent molecular killing events must produce a constant rate: if each of many targets is hit at random and independently, the chance that the fatal hit comes in any given hour is the same every hour (the standard Poisson argument).[4,5] The condition matters: the argument holds for a target pool large enough that depletion is negligible over the exposure, and not for a lesion requiring occupancy of a small discrete set. That exception bears directly on the anticholinesterases, where death follows from inhibiting a fixed enzyme pool past a critical fraction, so the constant-rate result should be read as the behaviour of the hazard away from that fraction rather than at it.

The constant-rate result also bears only on killing driven by the formation of damage. A hazard driven by the damage level gives the rising rate that Class A describes, and it can arise internally or externally. Internally, the organism can become progressively more fragile as damage accumulates. Write the instantaneous rate of fatal events as $h(t) = b_w\,[D(t) - z]^+$: each fatal event still arrives at random, but the rate at which they arrive is set by how much damage the animal is already carrying, so the same increment of stress is more likely to be fatal late in an exposure than early. Consuming a reserve gives the same form, with the remaining reserve playing the role of $-D$. That is the wear-out hazard of §S1.3, and a laboratory test removes neither it nor the reserve case. Externally, an encounter whose outcome the current impairment decides gives the same shape, and an acute test does remove such processes by design: predators, pathogens and competition for food are all absent.

##### S6.6 Heterogeneity Generated by the Running Maximum

Class B generates a tolerance distribution among organisms that begin identical. The running maximum acts as a ratchet: two organisms exposed to the same fluctuating profile, differing only in the random endurance of §S3.1, part ways irreversibly at the first peak that exceeds one's effective threshold and not the other's, and the divergence persists because the maximum never decreases. A spread of effective thresholds is not by itself peculiar to either structure: any hazard model can be written as death when the cumulative hazard first exceeds an independent exponential clock, so every organism carries an effective threshold whether or not one is assumed. What is peculiar to Class B is what that threshold is a threshold on. Under Class B it is a level of peak damage, so the spread has the units of damage, and the ratchet makes the divergence between two individuals irreversible. Under Class A it is the exponential clock on accumulated hazard, and an exponential clock has no memory: an animal whose clock has not yet run out after the first pulse is, statistically, as good as new — which is the coin-toss argument of Proposition S4 seen from the other side. No subgroup is selected. That is why the ratchet, and not any assumed heterogeneity, is what lets the peak-based structure exceed unity.

#### S7. Fourteen Compounds: What the Fitted Parameters and the Likelihood Each Show

Both hazard structures have been fitted to the same fourteen-compound *Gammarus pulex* dataset with toxicokinetics held identical between them.[2] Because the toxicokinetics are shared, the fitted toxicodynamic parameters can be compared directly. Two things can then be asked of them that goodness of fit cannot answer: whether the inferred damage recovery is consistent with the known chemistry of the target adduct, and whether the likelihood prefers one structure in a way that tracks mechanism.

**Table S3. Damage recovery rate under both hazard structures (Ashauer *et al.* 2015, supplementary parameter tables).**

| Compound | MoA | $k_{r,\mathrm{SD}}$ (d$^{-1}$) | $k_{r,\mathrm{IT}}$ (d$^{-1}$) | ratio | $t_{95}$ SD (d) | $t_{95}$ IT (d) | ΔNLL |
|---|---|---|---|---|---|---|---|
| 1,2,3-Trichlorobenzene | baseline | 1000 | 7.11 | 141 | 0.003 | 0.4 | +41 |
| 2,4-Dichloroaniline | baseline | 1000 | 1000 | 1.0 | 0.003 | 0.003 | +51 |
| 2,4-Dichlorophenol | baseline | 1000 | 17.1 | 58 | 0.003 | 0.2 | +91 |
| Pentachlorophenol | uncoupling | 1000 | 0.0677 | 14,771 | 0.003 | 44.2 | −15 |
| 2,4,5-Trichlorophenol | uncoupling | 1000 | 0.0116 | 86,207 | 0.003 | 258 | +52 |
| 4,6-Dinitro-o-cresol | uncoupling | 22.5 | 1.12 | 20 | 0.13 | 2.7 | +24 |
| Aldicarb | carbamate | 3.02 | 0.727 | 4.2 | 1.0 | 4.1 | +32 |
| Carbofuran | carbamate | 1000 | 3.53 | 283 | 0.003 | 0.8 | +11 |
| Carbaryl | carbamate | 0.122 | 0.0168 | 7.3 | 24.6 | 178 | −144 |
| Malathion | OP | 1.08 | 0.084 | 12.9 | 2.8 | 35.7 | +34 |
| Chlorpyrifos | OP | 0.630 | 0.0001 | 6,300 | 4.8 | 29,957 | +12 |
| Diazinon | OP | 0.150 | 0.0179 | 8.4 | 20.0 | 167 | −6 |
| 4-Nitrobenzyl-chloride | reactive | 0.0001 | 0.0001 | 1.0 | 29,957 | 29,957 | −164 |
| Sea-nine | reactive | 0.222 | 0.0001 | 2,220 | 13.5 | 29,957 | −1 |

$t_{95} = \ln 20/k_r$; ratio is $k_{r,\mathrm{SD}}/k_{r,\mathrm{IT}}$. $\Delta$NLL = $\mathrm{NLL_{SD}} - \mathrm{NLL_{IT}}$; positive values favour the individual-tolerance structure, which stands in for the peak-based one in the reduced framework (§S3.1). The fitting interval for $k_r$ was 0.0001–1000 $d^{-1}$, the range the tabulated values span.

**A note on incipient lethal concentrations.** Ashauer *et al.* state that incipient lethal concentrations cannot exist under stochastic death.[2] That holds in the limit $z \to 0$. For $z > 0$ the lethal concentration converges to $z$ itself, which is the sense used in main text §1. Their own fits give $z > 0$ for thirteen of the fourteen compounds; the exception, 2,4,5-trichlorophenol, has $z$ at zero with a killing rate of $1.26 \times 10^7$ and is degenerate.

##### S7.1 Most of the Recovery-Rate Ratios Are Not Estimates

Main text Figure 4 shows why. Nine of the fourteen compounds have at least one fit on a bound, and six of the cumulative-structure fits sit at the upper fitting bound of 1000 $d^{-1}$ — a four-minute recovery time, which is where the optimiser puts a parameter the data do not determine. A summary statistic taken over all fourteen therefore describes the bounds, not the compounds. Restricting to the five for which both fits are interior — 4,6-dinitro-*o*-cresol, aldicarb, carbaryl, malathion and diazinon — gives ratios of 20.1, 4.2, 7.3, 12.9 and 8.4 respectively, a factor of about 4 to 20 with a geometric mean of 9. That range, not a summary over all fourteen, is what main text §5.3 quotes. The direction, however, holds without qualification: the cumulative structure infers the faster recovery in twelve of the fourteen and never the slower, the remaining two having both fits at the same bound. The same asymmetry has been reported independently: of thirty-six datasets fitted under both structures, eighteen gave different dominant rate constants, and in every one of those the individual-tolerance estimate was the lower.[S13] The fitted rates themselves are main text Figure 4; recovery times and fit differences for the same compounds are in Figure S4.

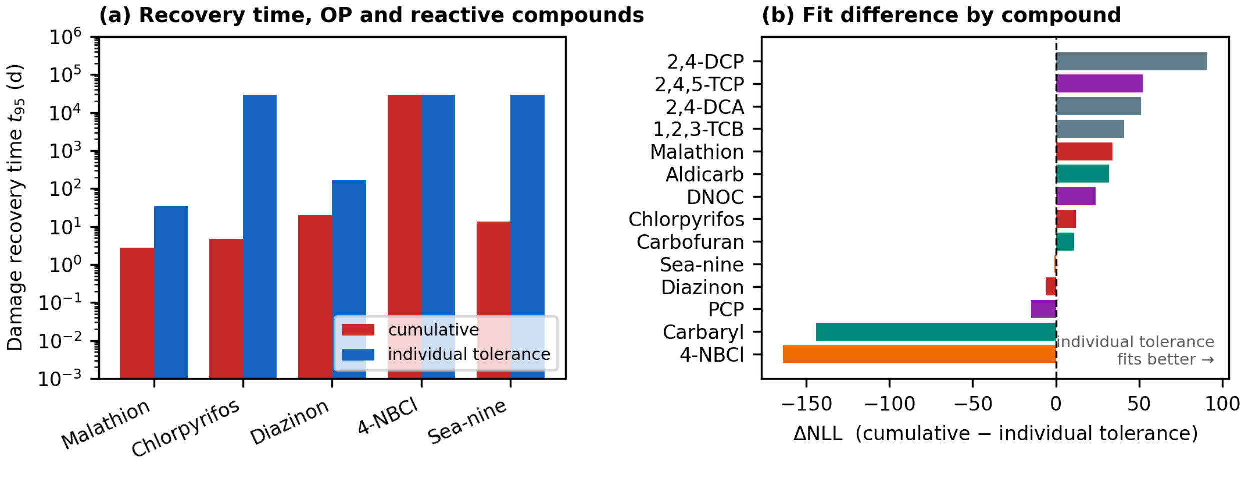


**Figure S4. Recovery times and fit differences across fourteen compounds (Ashauer *et al.* 2015).** The fitted recovery rates themselves are main text Figure 4. (a) Damage recovery time $t_{95}$ for the organophosphates and reactive compounds under each structure. (b) $\Delta$NLL by compound, sorted; the sign does not group by mode of action. Values in Table S3.

##### S7.2 What the Fitted Recovery Rates Measure

The cumulative structure infers damage recovery times of 2.8 d for malathion, 4.8 d for chlorpyrifos, 20 d for diazinon and 13.5 d for the reactive isothiazolone Sea-nine. The peak-based structure places chlorpyrifos and Sea-nine at the lower fitting bound, 0.0001 $d^{-1}$, a recovery time of some 30,000 days, which is a bound rather than an estimate.

The same comparison at the other end of the range is asymmetric in the same way. The cumulative structure drives $k_r$ to its **upper** bound for six of the fourteen compounds, among them all three baseline narcotics and pentachlorophenol; the peak-based structure reaches that bound for one. A bound is not an estimate, so neither number measures a recovery time. What the asymmetry shows is where each structure has to go to accommodate the same data, and the cumulative structure has to leave the identified region six times as often.

**Proposition S6 (a lower bound on the recovery rate under the cumulative hazard).** *In the reduced framework, $\dot{D} = k_r(C - D)$ with $D(0) = 0$, let one rectangular pulse of concentration $C_0$ and length $\tau_p$ raise damage above the threshold, $D\left(\tau_p\right) > z$, with no exposure afterwards. Under Class A, $h = b_w[D - z]^+$, the cumulative hazard after the pulse, relative to that during it, satisfies $\rho > 1/\left(e^{k_r \tau_p} - 1\right)$ for every $b_w$, $z$ and $C_0$, and the bound is approached as $D\left(\tau_p\right)/z \to 1$. Equivalently, $k_r > \ln(1 + 1/\rho)/\tau_p$. Under Class B, $\rho = 0$ at every $k_r$ (Proposition S2).*

*Proof.* Write $x = D\left(\tau_p\right)/z > 1$ and $r = 1/\left(e^{k_r \tau_p} - 1\right)$. After the pulse $D = xz\, e^{-k_r s}$ exceeds $z$ for $s < \ln(x)/k_r$, so $H_{\text{post}} = (b_w z/k_r)\ g(x)$ with $g(x) = x - 1 - \ln x$. During the pulse $D = C_0\left(1 - e^{-k_r t}\right)$ increases, with $C_0 = xz(1 + r)$; substituting $w = D/z$ gives $H_{\text{pulse}} = (b_w z/k_r)\ I(x)$, $I(x) = \int_1^x (w - 1)/(c - w)\, dw > 0$, $c = x(1 + r)$. Hence $\rho = g(x)/I(x)$. Let $F = g - rI$ and $y = 1 - 1/x$; then $F(1) = 0$ and $F'(x) = (1 + r)\ [\, y - r\ln(1 + y/r)\,] > 0$ for $x > 1$, because $\ln(1 + u) < u$, so $F > 0$ and $\rho > r$. As $x \to 1$, l'Hôpital's rule gives $\rho \to r$. Under Class B the hazard is zero once $D$ stops rising (Proposition S2). ▫

Applied to the first pulse of each two-pulse treatment of the archive (control-corrected, with the post-pulse window running to the second pulse; `sd_recovery_bound.py`), the bound is infinite for the five narcotic treatments whose survival stays flat after the pulse (1,2,3-trichlorobenzene A and C, 2,4-dichloroaniline A, 2,4-dichlorophenol A and B), where the published cumulative fits sit at 1000 $d^{-1}$, and 1.9–3.0 $d^{-1}$ for the other four narcotic treatments. Among the organophosphates it binds malathion: 0.63 and 0.67 $d^{-1}$ for treatments A and C, and no finite value for B, against a fitted 1.08 $d^{-1}$, and chlorpyrifos III: 1.9 $d^{-1}$ against a fitted 0.63 $d^{-1}$, which the bound taken at the upper 95% limit of the post-pulse hazard (0.21 $d^{-1}$) does not exclude. Diazinon A–C lose no more animals within the pulse than the controls, so the proposition sets no bound on their fitted rates. Recorded deaths lag the hazard, which biases the observed ratio upward and the bounds downward. The proposition covers one-compartment damage and a single pulse; it does not constrain the three-compartment fits of §S13.

Two further fitting pathologies appear only under the cumulative structure. For 2,4,5-trichlorophenol it returns $z = 0$ — no threshold — with a killing rate of $1.26 \times 10^7$, eight orders of magnitude above every other compound in that source's supplementary tables, whereas the peak-based structure returns ordinary values for the same data. And for chlorpyrifos the original authors record that fitting on the oxon alone drove $k_r$ to 1000 $d^{-1}$, forcing a change of dose metric.

The proposition is stated for a single threshold, the GUTS-RED-SD case; main text §5.3 and §5.5 use it only to say that a GUTS-SD recovery rate is not a recovery time.

##### S7.3 Goodness of Fit Carries No Mechanistic Signal

The likelihood difference between the two structures ranges from 164 units in favour of the cumulative structure to 91 in favour of the peak-based one, with a median of +18 favouring the latter and a mean of +1.3. The sign does not track mode of action: on likelihood the cumulative structure is ahead for one uncoupler, one carbamate, one organophosphate and two reactive compounds, and the peak-based structure for the remaining nine, with no grouping by mechanism in either set. A sign test on the fourteen does not separate that split from chance ($p = 0.42$). The original authors concluded that both models describe the data adequately for most compounds. They also noted the degeneracy main text §5.3 describes, calling the inference circular because the death mechanism cannot be separated from the speed of damage recovery.[2]

This is the point made by the control simulation of §S10 and by the diazinon comparison of main text §4.4, now at the scale of fourteen compounds and with the toxicokinetics shared between the two fits: the likelihood does not identify the hazard structure. The margins are nevertheless an order of magnitude larger here — reaching 164 units against a simulated median of ±1.6 — and the two regimes differ in exactly one respect. The simulation of §S10 held toxicokinetics and damage kinetics at their true values, so both structures read the same damage trajectory and the likelihood could respond only to the hazard. In the fourteen fits each structure fits its own recovery rate, and §S7.1 shows what it pays: the cumulative structure infers the faster recovery in twelve of the fourteen, by factors of 4 to 20 where both fits are interior and to the 1000 $d^{-1}$ upper bound in six cases. Two fits that disagree about the damage kinetics by an order of magnitude are not fitted to the same trajectory, and a likelihood difference between them measures how efficiently each structure absorbs misfit into the kinetics, not which hazard is true. The simulation frees only the hazard; the fourteen fits free hazard and kinetics together; and both say the same thing about calibration as a discriminator. What does discriminate is a quantity one structure cannot produce, the carry-over ratio of §S6.1; whether the fitted recovery rates can be read as recovery times (§S7.2, Proposition S6) constrains without discriminating, and on diazinon prediction of untested exposure patterns does not separate the structures either (§4.4).

The extrapolation that the choice of hazard structure governs spans a long time. Fish survival under regulatory exposure profiles has been predicted over 485 days from four-day acute tests, and the authors note that the extrapolation is inevitable but rarely stated, so its assumptions are not scrutinised.[S14] Where the structures are indistinguishable on the calibration data and diverge over that interval, the assumption being carried is exactly the one §S3.3 shows the calibration cannot settle.

#### S8. Compartmental Structure and Gap Closure

This section asks what the one-compartment (reduced) framework contributes to the likelihood difference on dimethoate: what the missing compartments do to damage curvature, and how much of the cumulative structure's reduced-framework advantage remains once they are restored. Because no toxicokinetic constant of dimethoate has been measured in this species, every three-compartment result here is reported at both ends of a borrowed range.

##### S8.1 Borrowed Toxicokinetics and an Assumed Metabolite Clearance Rate

No measurement of dimethoate or omethoate toxicokinetics exists for *G. pulex* or any other freshwater invertebrate. The parent compartment is therefore run with the measured *G. pulex* constants of the two compounds that bracket dimethoate (log $K_{OW}$ 0.70–0.78) in hydrophobicity in the same laboratory's set: imidacloprid (log $K_{OW}$ 0.33; $k_{in} = 1.96$ L kg$^{-1}$ d$^{-1}$, $k_{out} = 0.267$ d$^{-1}$) and aldicarb (1.13; 10.80 and 6.62), the MCMC estimates of that study's Table 1.[S15] The two sets are the ends of a range, not estimates for dimethoate, and they differ most in elimination: a 95% elimination time of 11 days for imidacloprid against half a day for aldicarb. The reduced framework needs none of these constants, which is why main text §4.4 leads with it.

The metabolite step is assumed as well. Dimethoate is a phosphorothioate and is not itself an inhibitor: oxidative desulfuration yields omethoate, which phosphorylates acetylcholinesterase, so the middle compartment of eq. 3 is omethoate and $k_{2,\text{out}}$ its elimination rate. No experiment separating omethoate from dimethoate in the tissue of this species has been reported, and $k_{2,\text{out}}$ is held at a nominal 3.0 $d^{-1}$. Two lines of evidence bound what it can plausibly be. First, the same laboratory measured internal concentrations of activated metabolites in this species for three other anticholinesterases.[21] Refitting the metabolite compartment to those series returns elimination rate constants of 1.2 $d^{-1}$ for chlorpyrifos-oxon, 6.8 $d^{-1}$ for aldicarb sulfone and 8.2 $d^{-1}$ for the principal malathion metabolite (`metabolite_clearance_fit.py`). Second, the only aquatic invertebrate study to resolve a full organophosphate–oxon toxicokinetic scheme reports, for diazinon in *Daphnia magna*, oxidative desulfuration at 0.48 $d^{-1}$ (0.34 $d^{-1}$ in the same group's later toxicokinetic–toxicodynamic model, §S4.1[S1]), oxidative dearylation to the detoxification product at 3.8 $d^{-1}$, and elimination of that product at 18.7 $d^{-1}$.[S16] The assumed 3.0 $d^{-1}$ lies inside the first range and between the desulfuration and dearylation rates of the second.

That study also carries a caveat that applies here. Internal diazoxon was near the limit of detection throughout, and its removal could not be resolved as a first-order clearance: the authors model it as proceeding solely by irreversible reaction with acetylcholinesterase.[S16] If omethoate behaves the same way, what the present model calls metabolite clearance is the damage-forming step itself, and $k_{2,\text{out}}$ and $k_3$ are not separately identified. §S8.3 therefore profiles $k_{2,\text{out}}$ instead of relying on the nominal value.

##### S8.2 What the Restored Compartments Do: Curvature and Delayed Damage

The damage trajectory under constant exposure, at the peak-based calibration fits, was fitted to a power law $D = a\,t^{n_{\text{eff}}}$ by log-log regression. Over the 4-day test the exponent is 0.98 for the reduced model, 1.78 for the three-compartment model with the aldicarb constants and 2.43 with the imidacloprid constants (Figure S5a, regression from 0.05 d). Table S4 resolves the aldicarb case by phase.

**Table S4. Effective power-law exponent $n_{\text{eff}}$ of three-compartment damage, by phase (aldicarb constants).** Peak-based calibration fits at $k_{2,\text{out}}$ = 3.0 $d^{-1}$ and with $k_{2,\text{out}}$ estimated, which goes to its lower bound, 0.05 $d^{-1}$. Regression from the first grid point on a 0.005-d grid.

| Time window | $k_{2,\text{out}}$ = 3.0 (fixed) | $k_{2,\text{out}}$ = 0.05 (estimated) |
|---|---|---|
| Phase 1: 0–0.3 d | 2.81 | 2.87 |
| Phase 2: 0.3–1.5 d | 1.87 | 2.36 |
| Phase 3: 1.5–4.0 d | 1.24 | 2.08 |
| **Full test: 0–4 d** | **1.92** | **2.40** |
| RED model (reference) | 1.00* | — |

The log-log regression weights every grid point equally, so the exponents depend a little on the grid and on where the regression starts: over grids of 0.05 to 0.005 d the whole-test exponent runs from 1.83 to 1.92 at the fixed clearance rate and from 2.35 to 2.40 at the estimated one, and the phase values move by at most 0.08 (`damage_curvature_and_step_size.py`); starting at 0.05 d gives the 1.78 of Figure S5a. *The reduced-model value is the early-time limit ($D \propto t$ for $k_r t \ll 1$); the reduced model cannot exceed an exponent of 1 at any time.

The trajectory passes through a near-$t^3$ phase while all three compartments are still filling and relaxes toward $t^1$ as the metabolite compartment approaches equilibrium; with the slower metabolite clearance it stays near $t^2$ through the test. The imidacloprid constants, whose slow parent elimination adds a further delay, give the steepest rise.

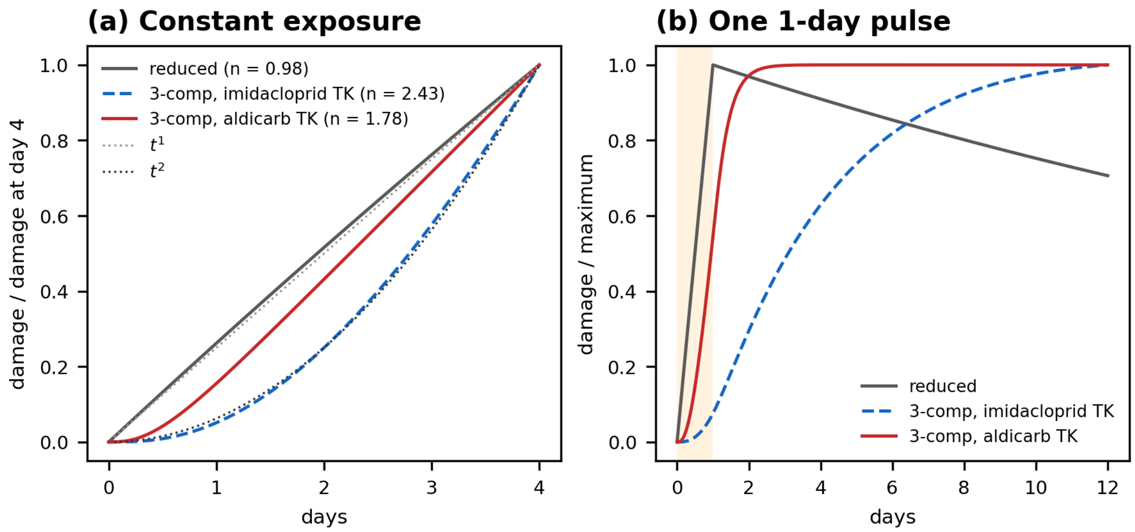


**Figure S5. Damage under the reduced and three-compartment models (dimethoate).** (a) Damage under constant exposure, normalised to its value at day 4, for the reduced model and the three-compartment model with the imidacloprid and the aldicarb constants ($k_{2,\text{out}} = 3.0$ d$^{-1}$; recovery rates of the peak-based calibration fits); the effective exponent over 0.05–4 d is in the legend; dotted lines $t^1$ and $t^2$. (b) Damage after a single 1-day pulse (shaded), normalised to its maximum. `figureS5_damage_curvature.py`.

**Damage after a pulse.** After a 1-day pulse the reduced model's damage peaks when the pulse ends. With three compartments the metabolite left in the middle compartment goes on feeding the damage state after the external exposure has stopped: damage peaks on day 5 with the aldicarb constants and is still rising on day 12 with the imidacloprid constants (Figure S5b). At the recovery rates of the peak-based fits, which sit at the slow end of their range, damage hardly declines after its peak, so within the 10-day pulse treatments the three-compartment fits never enter the phase in which the two hazard structures diverge (§S3.4).

##### S8.3 Gap Closure and the $k_{2,\text{out}}$ Profile

Gap closure is the fraction of the peak-based structure's reduced-framework calibration deficit that restoring the compartments recovers, $\left(\text{NLL}_B^{\text{RED}} - \text{NLL}_B^{\text{FULL}}\right) / \left(\text{NLL}_B^{\text{RED}} - \text{NLL}_{\text{SD}}^{\text{RED}}\right)$ (§S12.3). A value above 100% means that the three-compartment peak-based fit is better than the reduced cumulative one.

**Table S5. Gap closure, dimethoate 4-day LC50 calibration.** Best of 30 starts; exact toxicokinetic integration; recovery rate bounded at $t_{95} \geq 300$ d. In the estimated rows each structure estimates its own $k_{2,\text{out}}$.

| Configuration | Toxicokinetic constants | GUTS-PM NLL | GUTS-SD NLL | ΔNLL | Gap closure |
|---|---|---|---|---|---|
| RED (1-compartment) | none needed | 164.18 | 140.92 | −23.3 | — |
| 3-compartment, $k_{2,\text{out}} = 3.0$ | imidacloprid | 136.78 | 145.60 | +8.8 | 118% |
| 3-compartment, $k_{2,\text{out}}$ estimated | imidacloprid | 136.72 | 141.38 | +4.7 | 118% |
| 3-compartment, $k_{2,\text{out}} = 3.0$ | aldicarb | 148.40 | 137.84 | −10.6 | 68% |
| 3-compartment, $k_{2,\text{out}}$ estimated | aldicarb | 137.16 | 137.76 | +0.6 | 116% |

The peak-based fits change by at most 0.15 units with the recovery bound, since they go to the slow end of the recovery rate in almost every case; the cumulative fits change by up to 8 units, and every cumulative three-compartment fit sits at the bound (Table S17). With the aldicarb constants, freeing $k_{2,\mathrm{out}}$ sends the peak-based estimate to the lower bound, 0.05 $\mathrm{d}^{-1}$, and the cumulative structure's to 4.0 $\mathrm{d}^{-1}$.

**Table S6. Frozen-parameter prediction to the 10-day pulse treatments.** Calibrated parameters of Table S5, held fixed. The last column repeats the difference without the recovery bound (§S9.2).

| Configuration | Toxicokinetic constants | GUTS-PM NLL | GUTS-SD NLL | ΔNLL | ΔNLL, no bound |
|---|---|---|---|---|---|
| RED | none needed | 701.0 | 575.9 | −125.1 | −125.1 |
| 3-compartment, $k_{2,\mathrm{out}} = 3.0$ | imidacloprid | 570.0 | 1250.6 | +680.6 | +21.2 |
| 3-compartment, $k_{2,\mathrm{out}}$ estimated | imidacloprid | 576.4 | 1070.1 | +493.8 | +3.0 |
| 3-compartment, $k_{2,\mathrm{out}} = 3.0$ | aldicarb | 578.8 | 708.4 | +129.6 | +16.8 |
| 3-compartment, $k_{2,\mathrm{out}}$ estimated | aldicarb | 613.7 | 685.8 | +72.2 | −29.8 |

In the reduced framework the cumulative structure predicts the pulse treatments better by 125 units (Table S6). With three compartments and the recovery bound the peak-based structure predicts better in every configuration, by 72 to 681 units; the size is set by the borrowed constants and by the bound, and without the bound the advantage shrinks to 3–21 units or, with the aldicarb constants and $k_{2,\mathrm{out}}$ estimated, turns to the cumulative structure by 30 units.

**Numerical note.** The toxicokinetic chain is linear with piecewise-constant forcing, so it is propagated exactly by matrix exponential; only the Class A hazard integral requires quadrature. The choice of integrator matters. Under a forward-Euler scheme the calibration fits at $k_{2,\mathrm{out}} = 3.0$ (aldicarb constants; $\tau_{\mathrm{out}} = 1/k_{\mathrm{out}} = 0.15$ d, the parent's elimination time constant) depend on the step size (`damage_curvature_and_step_size.py`):

| $\Delta t$ (d) | $\Delta t/\tau_{\mathrm{out}}$ | GUTS-PM | GUTS-SD | ΔNLL |
|---|---|---|---|---|
| 0.1 | 0.66 | 154.3 | 138.3 | −16.1 |
| 0.05 | 0.33 | 151.4 | 137.9 | −13.5 |
| 0.025 | 0.17 | 149.9 | 137.8 | −12.1 |
| 0.0125 | 0.08 | 149.1 | 137.8 | −11.3 |
| exact | 0 | 148.4 | 137.8 | −10.6 |

Coarse steps worsen both fits, the peak-based one more (by 5.9 units at 0.1 d against 0.4), and so widen the cumulative structure's advantage from 10.6 to 16.1 units. Exact or high-order schemes are therefore needed whenever hazard structures are compared.

Because $k_{2,\mathrm{out}}$ was not measured, both structures were refitted at 15 values spanning 0.05–30 $\mathrm{d}^{-1}$ with all other parameters free, with each set of borrowed constants (Figure S6; `figureS6_metabolite_clearance_profile.py`).

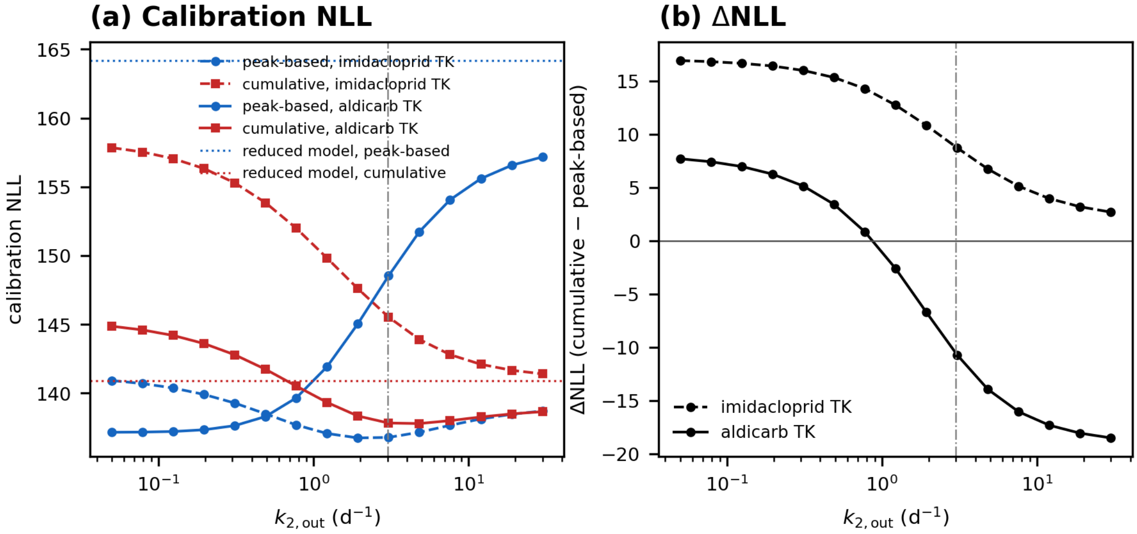


**Figure S6. Profile over the metabolite clearance rate $k_{2,\mathrm{out}}$ (dimethoate).** Recovery rate bounded at $t_{95} \geq 300$ d. (a) Calibration negative log-likelihood of the peak-based (blue) and cumulative (red) structures refitted at 15 fixed values of $k_{2,\mathrm{out}}$, with the imidacloprid (dashed) and aldicarb (solid) constants; dotted lines, the reduced-framework fits. (b) ΔNLL = $\mathrm{NLL}_{\mathrm{SD}} - \mathrm{NLL}_{\mathrm{PM}}$ across the same range. The vertical line marks the nominal 3.0 $d^{-1}$.

With the recovery bound in place and the imidacloprid constants, ΔNLL falls from +16.9 at 0.05 $d^{-1}$ to +8.7 at 3.0 and +2.7 at 30 without changing sign; gap closure stays between 100% and 118%. With the aldicarb constants it changes sign between 0.8 and 1.2 $d^{-1}$ and falls to −10.7 at 3.0 and −18.5 at 30; gap closure exceeds 50% up to 4.8 $d^{-1}$ and falls to 30%. Over the range of the three measured metabolite elimination rates, 1.2–8.2 $d^{-1}$ (§S8.1), ΔNLL runs from +12.8 to +5.1 with the imidacloprid constants and from −2.6 to −16.1 with the aldicarb constants. The sign of the three-compartment calibration comparison therefore depends on constants that have not been measured for dimethoate, and restoring the compartments closes 43–118% of the reduced-framework gap of 23.3 units within that range (30–118% over the whole profile).

Table S7 collects the calibration and out-of-sample margins of both case studies.

**Table S7. Calibration and out-of-sample ΔNLL, GUTS-SD minus the peak-based structure.** Positive favours the peak-based one; out-of-sample values freeze the calibrated parameters (main text §4.4). Recovery rate bounded at $t_{95} \geq 300$ d. Dimethoate three-compartment rows at $k_{2,\mathrm{out}} = 3.0$ $d^{-1}$; diazinon with the toxicokinetics measured in the same study,[20] calibrated on treatment A.

| Compound | Damage model | Calibration | Out of sample |
|---|---|---|---|
| Dimethoate | 1-compartment (RED) | −23.3 | −125.1 |
| Dimethoate | 3-compartment, imidacloprid constants | +8.8 | +680.6 |
| Dimethoate | 3-compartment, aldicarb constants | −10.6 | +129.6 |
| Diazinon | 1-compartment (RED) | +3.3 | −40.2 |
| Diazinon | 3-compartment | +4.8 | +4.6 |

#### S9. Parameter Identifiability

Four identifiability results bear on the case studies. The damage recovery rate and the damage scale are not identified, though the predictions are (§S9.1); the cumulative structure depends on any bound placed on the recovery rate, and so does the sign of the diazinon comparison (§S9.2); the tolerance shape is fixed rather than estimated, and §S9.3 reports what that changes; and the killing rate and tolerance shape are identified only jointly (§S9.4).

##### S9.1 Neither $k_r$ nor the Damage Scale Is Identified; the Predictions Are

The recovery rate of the damage state, $k_r$, is not identified by these data, and the two structures go to opposite ends of its range. Within the bound of the main analysis, $10^{-5}$–$10^{-2}$ d$^{-1}$ ($t_{95} \geq 300$ d; Table M1), it goes to the lower end for the peak-based structure on both compounds ($t_{95} \approx 3 \times 10^5$ d) and to the upper end, the bound itself, for the cumulative structure (Table S17). Without the bound the cumulative structure returns a $t_{95}$ of 15 d for dimethoate (aldicarb constants) and 21 d for diazinon (§S9.2).

**The peak-based likelihood is insensitive to $k_r$.** Its calibration likelihood is the same to two decimals, 148.40 for dimethoate and 195.84 for diazinon, whether $k_r$ is left free or held at $t_{95} \geq 30$, 100 or 300 d (Tables S9 and S10). The data are consistent with recovery as slow as the range allows and do not separate any value of $t_{95}$ above 30 d from another.

**The parameters are not identified; the predictions are.** Two peak-based fits to dimethoate with the same calibration and prediction likelihoods (148.40 and 578.8) show this. One ties the recovery rate to the damage-formation rate, $k_r = 0.01\,k_3$, which holds $k_3$ at its lower bound of 0.01 d$^{-1}$; the other estimates $k_r$ on its own. Between them the median threshold moves by a factor of 13, from 497 to 38. Damage scales with the product $k_2 k_3$, and that product, the threshold and the reciprocal of the killing rate all move by the same factor (Table S8).

**Table S8. The three quantities that move together between two equally good fits (dimethoate, peak-based structure).** Tied imposes $k_r = 0.01\,k_3$; free estimates $k_r$ within $10^{-5}$–$10^{-2}$ d$^{-1}$. Three compartments, aldicarb constants, metabolite clearance 3.0 d$^{-1}$ in both (`recovery_rate_free_refit.py`).

| Quantity | tied | free | free/tied |
|---|---|---|---|
| $k_2 k_3$ (d$^{-2}$) | 0.0803 | 0.00617 | 0.077 |
| $\theta_{\text{med}}$ | 497 | 38.2 | 0.077 |
| $k^{-1}$ (damage units · d) | 3479 | 267 | 0.077 |

The agreement is exact rather than approximate. Because $H_B = k\,[M(t) - \theta]^+$ and $M \propto k_2 k_3$, the hazard is invariant under the rescaling $\left(k_2 k_3,\ \theta,\ k^{-1}\right) \to \lambda\,\left(k_2 k_3,\ \theta,\ k^{-1}\right)$ for any $\lambda > 0$. Damage is a latent variable with no observable scale, so this rescaling is a symmetry of the likelihood and no quantity of survival data can break it: only the dimensionless ratio $M/\theta$ is estimable. The same relation holds for the cumulative structure, where the product, the threshold and the reciprocal of the killing rate move by a factor of about 9 between the same two fits. The size of the factor carries no information: it is set by where the optimiser stops on the ridge, and only the equality of the three ratios is a property of the model. Reported parameter values are therefore one point on a curve of equally good fits rather than estimates.

The six dimethoate fits show what that costs. Across two hazard structures and three levels of compartmental detail the fitted threshold spans 275-fold, from 5.3 to 1,459. Divided by the damage the calibration exposure actually produced, the same six values span 4.1-fold and place the onset of killing at 0.6–2.4% of peak damage (Table S8b).

**Table S8b. The threshold alone and against the damage it is compared with (dimethoate).** Fitted by maximum likelihood on the LC50 series; three-compartment rows with the aldicarb constants. Each threshold is one point on its likelihood ridge and can differ from the refit of Table S17; the last column is the identified quantity (`threshold_scale_check.py`).

| Configuration | Structure | $\theta$ | max $D$ | $\theta$ / max $D$ |
|---|---|---|---|---|
| RED (1-compartment) | SD | 170 | 25,965 | 0.0065 |
| RED (1-compartment) | PM | 633 | 25,965 | 0.0244 |
| 3-compartment, $k_{2,\text{out}}$ fixed | SD | 5.3 | 880 | 0.0060 |
| 3-compartment, $k_{2,\text{out}}$ fixed | PM | 1459 | 98,526 | 0.0148 |
| 3-compartment, $k_{2,\text{out}}$ estimated | SD | 252 | 40,595 | 0.0062 |
| 3-compartment, $k_{2,\text{out}}$ estimated | PM | 192 | 31,990 | 0.0060 |

That a threshold reported without its damage scale is not interpretable is the practical content of the symmetry, and it is why main text §5.3 compares thresholds only after dividing each by the damage its own fit produces. Tying $k_r$ to $k_3$ does not identify anything; it fixes the scale by fixing $k_3$, and does so silently. The control simulation of §S10 makes a related point: two structures can fit the same data equally well even when the damage trajectory is known exactly.

The difficulty is not peculiar to this dataset. The companion analysis finds the repair capacity $K$ flat over seven orders of magnitude for a receptor agonist. Raths *et al.*, fitting a receptor-binding model to thiacloprid in this species, report that the association rate could not be identified at all. The optimiser reached the upper bound imposed on it, and the authors conclude that binding may as well be treated as instantaneous.[30] Three rate constants are therefore not recoverable from survival or residue data: $k_r$ here, the repair capacity of the companion analysis, and the association rate of Raths *et al.*, in three frameworks fitted by three groups. This is the reason the classification here is argued on structural rather than parametric grounds.

##### S9.2 What a Bound on the Recovery Rate Does to the Cumulative Structure

The main analysis bounds the damage recovery rate at $t_{95} \geq 300$ d, on the ground that irreversible phosphorylation of acetylcholinesterase implies a recovery time of the order of the enzyme's resynthesis. The value 300 d is a convention, not a measurement, and the recovery inferred from the lesion itself is shorter: rates fitted to clean-water acetylcholinesterase data in diazinon-exposed *Daphnia magna* place 95% recovery from 60% inhibition at about 17 d[18] and 99% recovery at about sixteen days,[S1] both extrapolated from about 80 h of data (§S14.3). Tables S9 and S10 therefore report the main results with the bound relaxed to 100 and 30 d and removed. Where the peak-based likelihood is indifferent to the recovery rate, the cumulative one depends on it (Table S9 for dimethoate, Table S10 for diazinon).

**Table S9. Sensitivity to a bound on the recovery rate (dimethoate, three compartments, aldicarb constants, $k_{2,\text{out}} = 3.0$).** `recovery_rate_bound_sensitivity.py`.

| Bound on $k_r$ | GUTS-PM NLL | GUTS-SD NLL | ΔNLL | SD $\hat{t}_{95}$ (d) | prediction ΔNLL |
|---|---|---|---|---|---|
| $t_{95} \geq 300$ **d (main analysis)** | **148.40** | **137.84** | −10.6 | **300** | +129.6 |
| $t_{95} \geq 100$ d | 148.40 | 137.79 | −10.6 | 100 | +111.6 |
| $t_{95} \geq 30$ d | 148.40 | 137.66 | −10.7 | 30 | +61.7 |
| none | 148.40 | 137.60 | −10.8 | 15 | +16.8 |

The same sensitivity for diazinon is in Table S10.

**Table S10. The same for diazinon (three compartments, measured toxicokinetics; calibration on treatment A, prediction of B and C).**

| Bound on $k_r$ | GUTS-PM NLL | GUTS-SD NLL | calibration ΔNLL | prediction ΔNLL | SD $\hat{t}_{95}$ (d) |
|---|---|---|---|---|---|
| $t_{95} \geq 300$ d **(main analysis)** | **195.84** | **200.62** | +4.8 | +4.6 | **300** |
| $t_{95} \geq 100$ d | 195.84 | 199.02 | +3.2 | +3.0 | 100 |
| $t_{95} \geq 30$ d | 195.84 | 195.96 | +0.1 | −1.5 | 30 |
| none | 195.84 | 195.56 | −0.3 | −3.6 | 21 |

Three things follow. First, the peak-based likelihood is the same in every row. Second, without a bound the cumulative structure returns recovery times of 15 d for dimethoate and 21 d for diazinon, of the same order as the lesion-based estimates above and as the 20 d of its fit to diazinon in the fourteen-compound set (Table S3). Third, the bound moves the comparisons by different amounts. On dimethoate the calibration difference hardly moves (by 0.2 units), while the prediction difference falls from +129.6 with the bound to +16.8 without it. On diazinon both differences change sign between the 300-d bound (+4.8, +4.6) and no bound (−0.3, −3.6), so the sign of the diazinon comparison is set by an assumed bound that the lesion-based estimates do not support, and stays within 5 units either way.

The recovery rate also decides whether the post-equilibrium phase of §S3.4 is reached. With the peak-based fits at the lower bound of $k_r$, their three-compartment damage does not decline within any treatment of either experiment (Figure S5b for dimethoate).

##### S9.3 The Fixed Tolerance Shape: What Changes When It Is Freed

The case-study fits hold the tolerance shape $\beta$ at 5.0 for both structures. The value is a fixed convention, not an estimate: it is the midpoint of the 2–8 interval over which the ordering scan of §S3.3 is run, and holding it equal is what isolates the hazard-structure comparison (main text §3). It is above what the case-study data return when the shape is estimated: 1.1–3.0 for dimethoate where the estimate is not at a bound, and 2.2–2.4 for diazinon (Tables S11 and S12); the published shapes for the fourteen compounds span 0.63–8.23 (ref 2, supplementary Table S5). All fits use the recovery bound of the main analysis and are by `tolerance_shape_sensitivity.py`.

**Table S11. Dimethoate calibration and prediction with the tolerance shape fixed and estimated.** Three-compartment rows with the aldicarb constants. The fixed rows repeat Tables S5 and S6; in the estimated rows each structure estimates its own $\beta$.

| Framework | $\beta$, PM / SD | Class B NLL | Class A NLL | $\Delta NLL_{cal}$ | gap closure | $\Delta NLL_{pred}$ |
|---|---|---|---|---|---|---|
| RED | 5.0 fixed | 164.18 | 140.92 | −23.3 | — | −125.1 |
| 3-comp, $k_{2,out} = 3.0$ | 5.0 fixed | 148.40 | 137.84 | −10.6 | 68% | +129.6 |
| 3-comp, $k_{2,out}$ estimated | 5.0 fixed | 137.16 | 137.76 | +0.6 | 116% | +72.2 |
| RED | 1.46 / 1.00† | 162.42 | 140.52 | −21.9 | — | −130.3 |
| 3-comp, $k_{2,out} = 3.0$ | 2.96 / 1.24 | 147.84 | 137.31 | −10.5 | 67% | +93.7 |
| 3-comp, $k_{2,out}$ estimated | 15.85† / 1.14 | 136.90 | 135.76 | −1.1 | 117% | −17.7 |

† at a bound of $\beta$.

On dimethoate (Table S11), freeing the shape changes the calibration differences by at most 1.7 units and the gap closure by about one percentage point. The reduced-framework comparison keeps its size in prediction as well (−125.1 and −130.3). The three-compartment prediction at the nominal clearance rate stays in favour of the peak-based structure (+129.6 and +93.7); with the clearance rate estimated as well it turns to the cumulative structure (−17.7), with the peak-based shape at its upper bound. The cumulative structure gains little from compartmental structure with the shape free (140.5 to 135.8–137.3); the peak-based structure gains 14.6–25.5 units.

**Table S12. Diazinon calibration (treatment A) and prediction (B and C) against the tolerance shape.** Three compartments, measured toxicokinetics, recovery bounded at $t_{95} \geq 300$ d; the first four rows hold $\beta$ equal for both structures, the last estimates it for each. Best of three multi-start runs of 30 starts each.

| $\beta$ (PM / SD) | $\Delta$NLL$_{\text{cal}}$ | Class B, prediction | Class A, prediction | $\Delta$NLL$_{\text{pred}}$ |
|---|---|---|---|---|
| 2.0 | +2.9 | 430.4 | 417.9 | −12.5 |
| 3.0 | −2.0 | 424.7 | 415.0 | −9.7 |
| 5.0 | +4.8 | 403.3 | 407.9 | +4.6 |
| 8.0 | +4.9 | 405.9 | 407.9 | +2.0 |
| 2.23 / 2.44 (estimated) | +1.6 | 446.4 | 409.7 | −36.7 |

On diazinon (Table S12) the calibration differences stay within 5 units of zero at every shape. The prediction difference changes sign with the shape: it favours the peak-based structure at 5 and 8 (+4.6, +2.0) and the cumulative structure at 2 and 3 (−12.5, −9.7) and at the shapes the data prefer when each structure estimates its own (−36.7), where both structures fit the calibration treatment better than at 5.0 (by 7.0 and 10.2 units). Only the last value lies outside the sampling interval of Table S14 (−16.0 to +25.5).

**The two comparisons answer different questions.** Holding the shape equal changes one thing — the form of the hazard — and isolates the hazard-structure choice. Allowing it to differ changes two, and the outcome then reflects the flexibility of the tolerance distribution as much as the form of the hazard. All comparisons in §4 hold the shape equal at 5.0 for that reason.

##### S9.4 Killing Rate and Tolerance Shape Are Not Separately Identifiable

The killing rate $k$ and the shape parameter of $f$ enter $F_{\oplus}$ jointly (§S3.1). They are separately identifiable only when the damage trajectory is externally constrained, either mechanistically (a measured or mechanistic damage model) or by giving every animal the same threshold. In the latter case the two-parameter form $(D_0, k)$ of §S2.2 carries the same parameter count as GUTS-RED-IT but derives its tolerance shape from the killing rate instead of assuming a log-logistic. A three-parameter reduced peak-based model $(\theta_{\text{med}}, \beta, k)$ is correspondingly over-specified, and the redundancy is quantitative. Across $k \in [0.05, 5]$ d$^{-1}$ and $\beta \in \{2, 5\}$ (scaled damage with $k_d = 1$ d$^{-1}$, seven concentrations in a $\sqrt{2}$ series with the top one killing about 90% in four days), a two-parameter log-logistic GUTS-RED-IT fitted to the peak-based survival reproduces it to within $\max|\Delta S| = 0.035$ within the 4-day constant-exposure design and 0.027 under frozen-parameter extrapolation to a 14-day scenario of two one-day pulses a week apart (`killing_rate_shape_redundancy.py`). That is well inside the binomial standard error of $\approx 0.11$ at $n = 20$ per concentration. Fitted median tolerances differ substantially between the two parameterisations ($m_w/\theta_{\text{med}} = 1.2$–15 over the same range, the largest at the slowest killing rate), so $\theta_{\text{med}}$ and the GUTS-RED-IT $m_w$ are not interchangeable as regulatory threshold estimates.

#### S10. Control Simulation and the Sampling Variability of the Margins

##### S10.1 With Everything but the Hazard Correct, the Likelihood Has Low Power, Not a Preference

The case studies of main text §4.4 compare the two structures under damage models of differing completeness, so a likelihood difference between them cannot be assigned to the hazard structure rather than to the adequacy of the damage model. This section removes the second factor by giving both models the same, correctly specified damage trajectory.

Two sets of $N = 300$ synthetic 96-hour datasets were generated, one from each structure, on the same damage trajectory: a two-stage chain in which the internal concentration equilibrates at $k_e = 0.8$ d$^{-1}$ and damage is formed at $k_a = 1.0$ and repaired at $k_r = 0.5$ d$^{-1}$, so that damage rises as $t^2$ early (§S4). The peak-based set has $k = 0.45$ and median tolerance $\theta_{\text{med}} = 3.0$; the cumulative set has $b_w = 0.75$ and median threshold $z_{\text{med}} = 3.0$, chosen so that survival at the top concentration matches the peak-based set to within a few per cent. Both use the log-logistic shape $\beta = 5.0$ of the case studies (main text §3.2) and no background mortality. Each dataset was fitted by both structures with the damage parameters at their true values, so that only the two hazard parameters were free. The design follows OECD TG 203: seven concentrations in a $\sqrt{2}$ series spanning an eightfold range, twenty animals each, observations at 24, 48, 72 and 96 h; each fit took the best of twenty Nelder–Mead starts. The tolerance integral uses the unit-sum quadrature of §S12.1, and that choice matters: weights that sum to less than one (0.88 in an earlier implementation) multiply every survival probability by their sum, and that error alone was enough to make the two structures look separable.

Table S13 summarises both directions.

**Table S13. Control simulation: what the calibration likelihood recovers when the damage trajectory is correct.** $N = 300$ per direction. The sign convention of main text §3.3 puts a positive ΔNLL in favour of the peak-based structure.

| Generating structure | Picks the generating structure | Margin under two units | Median ΔNLL |
|---|---|---|---|
| Class B (peak-based) | 92% | 55% | +1.6 |
| Class A (cumulative) | 84% | 60% | −1.6 |

Main text Figure 3 shows the two distributions the table summarises. They are mirrored about zero, which is why each structure is picked most of the time when it generated the data (first column); they overlap across most of their range, which is why the margin is usually under two units (second column). The likelihood therefore points the right way more often than not, but rarely by enough to decide: it has low power, not a preference. Against these distributions the calibration margins of main text §4.4 fall in two groups. The diazinon margin with measured toxicokinetics, +4.8, lies inside both: at the 87th percentile of the peak-based set and the 99.7th of the cumulative-generated one. The dimethoate margins in the reduced framework, −23.3, and with three compartments and the aldicarb constants, −10.6, lie beyond the most negative of the 600 replicates (−8.6; `control_simulation_both_directions.csv`). With the damage trajectory right, the hazard alone does not produce margins of that size, so they are not evidence about the hazard; for dimethoate they are consistent with the missing activation stage of main text §2.5 and with toxicokinetic constants that have not been measured (§S8). When each structure is also free to choose its own damage recovery rate, as in the fourteen-compound fits, the margins are about ten times larger than in the simulation (median +18; §S7.3).

##### S10.2 How Much the Reported Margins Vary with the Survivor Counts

The margins of main text §4.4 are point values, and their variability was obtained in two ways (Table S14). At the reported fit, the survivor counts of each treatment were redrawn from their observed conditional survival probabilities and the margin recomputed from the calibrated survival curves, which are held fixed (5,000 replicates, `margin_intervals.py`). For the dimethoate reduced-framework calibration, the margin the main text leads with, the animals were resampled and both structures refitted on each replicate (120 replicates, `margin_bootstrap_refit.py`), which includes the uncertainty of the fit itself.

**Table S14. Sampling variability of the margins of main text §4.4.** Percentile intervals; three-compartment rows with the recovery bound of the main analysis, dimethoate with the aldicarb constants at $k_{2,\mathrm{out}} = 3.0$.

| Margin | Method | Point | 95% interval |
|---|---|---|---|
| Dimethoate, reduced framework, calibration | refit, 120 replicates (median −23.3) | −23.3 | −33.7 to −14.4 |
| Dimethoate, three-compartment calibration | counts redrawn, 5,000 | −10.6 | −20.5 to −0.3 |
| Diazinon, three-compartment calibration | counts redrawn, 5,000 | +4.8 | −15.6 to +24.7 |
| Diazinon, three-compartment prediction | counts redrawn, 5,000 | +4.6 | −16.0 to +25.5 |

Both dimethoate intervals lie wholly below zero; their size is the subject of §S10.1 and §S8. Both diazinon intervals include zero and extend about 16–25 units either side of it: with measured toxicokinetics, the recovery bound and the common shape $\beta = 5$ the counts do not separate the structures in calibration or in prediction; with the shape estimated for each structure the prediction margin lies outside its interval (§S9.3). The resampled intervals give the variability of each comparison at fixed parameters; §S9.3 reports separately how far the margins move when the tolerance shape is freed.

#### S11. Choosing the Calibration Data: Only Combined Calibration Holds Every Treatment

Restoring compartments changes the model; this section holds the model fixed — the peak-based structure with three compartments — and changes the data it is calibrated on. Five calibration strategies, differing in concentration range and in the number of exposure patterns, are compared on how well each predicts the three dimethoate treatments.

##### S11.1 The Five Strategies

S1 uses the LC50 test alone: a standard four-day dose–response, which supplies concentration–response information and no temporal dynamics. S2 uses treatment C, a sustained exposure at about 9,000 nmol $L^{-1}$. S3 uses treatment A, two one-day pulses at about 44,000 nmol $L^{-1}$ one day apart, which supplies damage and recovery at a single concentration. Treatment B has the same pulses five days apart. S4 combines A, B and C, so both concentrations and all three exposure patterns are represented. S5 adds the LC50 series to S4.

**Table S15. Calibration strategies and their information content.** Every strategy fits the same five parameters of the peak-based three-compartment model (aldicarb constants, $k_{2,\mathrm{out}} = 3.0\ \mathrm{d}^{-1}$; bounds in Table M1).

| Strategy | Calibration data | Conc. range (nmol $L^{-1}$) | Duration | Exposure patterns | Free parameters |
|---|---|---|---|---|---|
| S1: LC50 | 96h LC50 (7 conc.) | 895–218,529 | 4 d | Constant, 7 conc. | 5 |
| S2: Trt C | Treatment C only | ~9,000 | 10 d | Sustained | 5 |
| S3: Trt A | Treatment A only | ~44,000 | 10 d | Pulses 1 d apart | 5 |
| S4: A+B+C | Treatments A+B+C | ~9,000 + ~44,000 | 10 d | Pulses 1 d and 5 d apart; sustained | 5 |
| S5: LC50+ABC | LC50 + A+B+C | Full range | 4 + 10 d | All of S1 and S4 | 5 |

Predictions under each strategy are shown in Figure S7. The individual parameter estimates move along the likelihood ridge of §S9.1 from one strategy to the next — the peak-based threshold from 7 (S1) to 2,652 (S4) — while the predictions in Table S16 move far less.

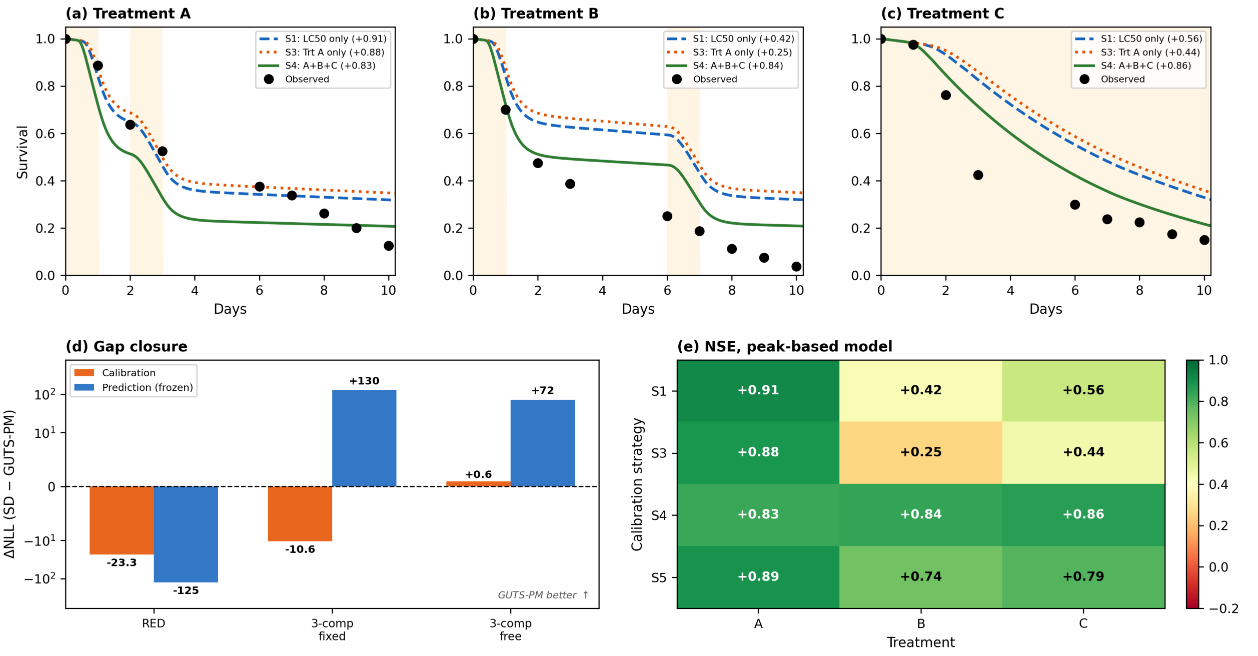


**Figure S7. Dimethoate: calibration strategy comparison and gap closure.** Three compartments, aldicarb constants. (a–c) Survival predictions for Treatments A, B and C under three calibration strategies: S1 (LC50 only, blue dashed), S3 (Treatment A only, orange dotted), S4 (A+B+C combined, green solid), with each strategy's Nash–Sutcliffe efficiency in the legend; observed data as filled circles, exposure windows shaded. (d) ΔNLL (SD − GUTS-PM) for calibration and for frozen-parameter prediction, across RED, 3-compartment with $k_{2,\mathrm{out}}$ fixed, and 3-compartment with $k_{2,\mathrm{out}}$ estimated; symmetric-log scale. (e) Nash–Sutcliffe efficiency for the peak-based model, strategies S1 and S3–S5 of Table S16 × three treatments. Panel (d) is discussed in §S8.3 (`case_dimethoate.py`).

##### S11.2 Single-Source Calibration Fails on at Least One Treatment; Combined Calibration Is the Most Uniform

**Table S16. Nash–Sutcliffe efficiency of each strategy's predictions, by treatment.** Peak-based structure, three-compartment model (aldicarb constants, $k_{2,\mathrm{out}}$ fixed), exact toxicokinetic integration (§S12.1). A treatment used for calibration is predicted in sample.

| Strategy | Trt A NSE | Trt B NSE | Trt C NSE | Calibration NLL |
|---|---|---|---|---|
| S1: LC50 only | +0.91 | +0.42 | +0.56 | 148.4 |
| S2: Trt C only | +0.39 | +0.95 | +0.96 | 157.5 |
| S3: Trt A only | +0.88 | +0.25 | +0.44 | 203.9 |
| **S4: A+B+C** | **+0.83** | **+0.84** | **+0.86** | **561.1** |
| S5: LC50+A+B+C | +0.89 | +0.74 | +0.79 | 714.0 |

Calibration NLL values are not comparable across strategies because each is computed on a different dataset; they are reported only to document the fit achieved. The comparison of interest is how each strategy's efficiencies vary across the three treatments — each row of Table S16 read across. S1 and S3 predict the closely spaced pulses (A) well, +0.91 and +0.88, but not the widely spaced pulses (B) or the sustained exposure (C), down to +0.42 and +0.25; S2, calibrated on C, predicts B and C but not A (+0.39). S4 is the only strategy with no NSE below +0.83, which is the sense in which combined calibration is the most uniform rather than the best on any single treatment.

In Treatments B and C both S1 and S3 predict too little mortality (Figure S7b, c), the non-conservative direction in a regulatory context, and S3, calibrated on the closely spaced pulses alone, by the most. Combined calibration S4 avoids this without additional model complexity. S5, which adds the LC50 series to S4, is less uniform than S4, with its lowest efficiency +0.74 on Treatment B.

##### S11.3 Two Comparisons Outside the Strategies

**GUTS-RED-IT alongside the two structures.** Every other comparison in this SI reports the difference between the cumulative and peak-based structures; Figure S8 instead shows each of three reduced structures on its own — peak-based, cumulative and GUTS-RED-IT — because GUTS-RED-IT is the model a Tier-2C dossier is most likely to contain alongside GUTS-RED-SD. The individual-tolerance fit is the $k \to \infty$ corner of the peak-based family (§S2.1), so the peak-based fit cannot be worse, and on dimethoate it is better by 87 to 98 units at every level of compartmental detail (aldicarb constants for the three-compartment fits; `figureS8_reduced_structures.py`). That gap measures what a finite killing rate is worth on these data — the delay it allows between crossing the threshold and dying — and the same delay is why the IT fit places its threshold at 21–28% of the peak damage the exposure produces, against 0.6–2.4% for the peak-based fits (Figure S8c).

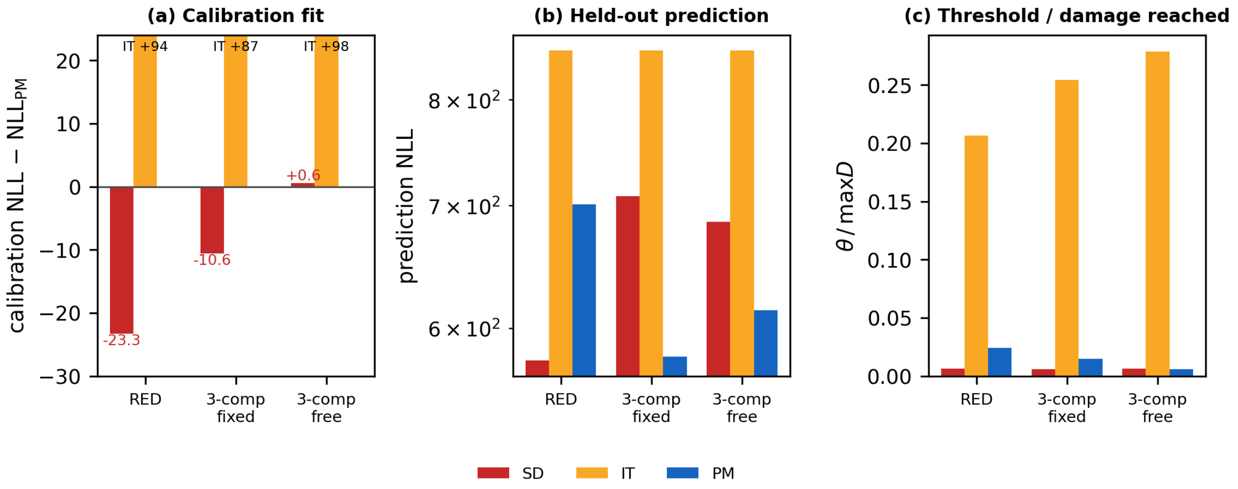


**Figure S8. The three reduced structures side by side, at three levels of compartmental detail (dimethoate).** (a) Calibration NLL relative to the peak-based fit within each configuration; the IT bars run off the axis and carry their values. (b) Held-out prediction with the calibration parameters frozen. (c) The fitted threshold divided by the damage the calibration exposure produced, because the raw value has no observable scale (§S9.1, Table S8b). The IT bars are GUTS-RED-IT as normally implemented, with a log-logistic tolerance and no killing rate — the $k \to \infty$ corner of the peak-based family, not the convolved form that §S3.1 shows to be observationally equivalent to it.

**The two case studies side by side.** Figure S9 sets the calibration and out-of-sample outcomes for dimethoate and diazinon side by side, the two compounds on which both damage models were fitted.

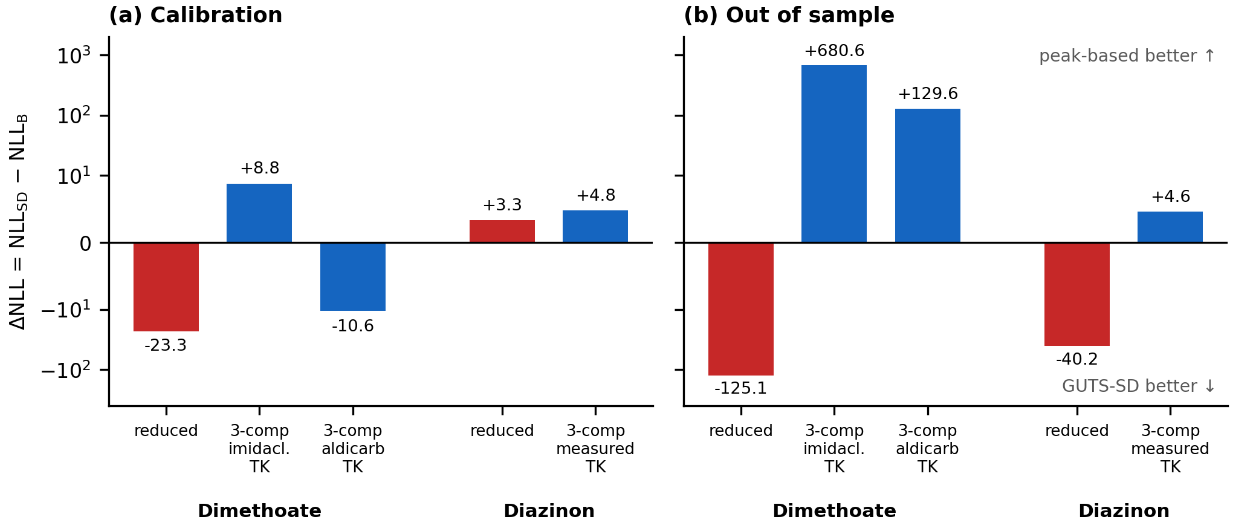


**Figure S9. The two case-study compounds side by side.** $\Delta\text{NLL} = \text{NLL}_{\text{SD}} - \text{NLL}_{\text{B}}$ on a symmetric-log axis; positive values favour the peak-based structure. Red, the reduced framework; blue, three compartments — for dimethoate with the borrowed imidacloprid and aldicarb constants, for diazinon with the toxicokinetics measured in ref 20. (a) Calibration: in the reduced framework the cumulative structure fits dimethoate better by 23.3 units, and the three-compartment fits close 68–118% of that gap; for diazinon both values lie within 4.8 units of zero. (b) Out of sample, with the calibrated parameters frozen: the reduced framework favours the cumulative structure for both compounds; with three compartments the dimethoate predictions favour the peak-based structure at the nominal metabolite clearance rate (Table S6 gives the estimated-clearance rows), and the diazinon prediction differs by 4.6 units in favour of the peak-based structure. Recovery rate bounded at $t_{95} \geq 300$ d. Values are read from Table S7 (`figureS9_case_studies.py`); the damage trajectories behind the dimethoate comparison are Figure S5.

### Part III. Procedures, Parameter Tables, Design and Deposit

*How this Part supports the main text.* §S12 gives the methods of §3 in full, §S13 every fitted and fixed parameter with the chlorpyrifos–carbaryl sequential experiment of §4.4, §S14 the design conditions the recommendation of §5.5 turns on, and §S15 the deposit that regenerates the tables and the figures listed there.

#### S12. Materials and Methods (Detailed)

This section records, at the level a re-implementation needs, the choices the main text's §3 states in outline: numerical integration, software and notation (§S12.1), estimation (§S12.2), the comparison metrics and interval rule (§S12.3) and the data (§S12.4).

##### S12.1 Numerical Integration, Software and Notation

The toxicokinetic chain of the three-compartment model (main text eq. 3, §3.1) is linear with piecewise-constant forcing and is propagated exactly by matrix exponential. In time, only the cumulative (Class A) hazard integral needs a numerical rule: the trapezoidal rule on a grid of $\Delta t = 0.01$–$0.02$ d (§S8.3). The running maximum was tracked incrementally: $M(t_i) = \max(M(t_{i-1}), D(t_i))$. Population survival was computed by integrating over the log-logistic tolerance distribution with a midpoint rule on the probability scale — eight nodes at $p_i = \left(i - \frac{1}{2}\right)/8$, uniform weights summing to unity — which avoids the endpoints, where the log-logistic quantile function diverges. Raising the number of nodes to 16, 32 or 64 changes every case-study calibration likelihood by at most 0.27 NLL units at the fitted parameters, and the calibration differences between the structures by at most 0.27 (`quadrature_nodes_check.py`); the control simulation of §S10, which frees only two hazard parameters, uses 32.

All simulations were implemented in Python; the deposited outputs were produced on Ubuntu 24.04 (Windows Subsystem for Linux 2) with Python 3.13.11, NumPy 2.4.0, SciPy 1.17.0, pandas 2.3.3 and Matplotlib 3.10.8.[S17] Parameters on a likelihood ridge (§S9.1; Tables S8, S8b and S17) can stop at a different point of the ridge under other library versions; the likelihoods and predictions do not change. Complete analysis scripts for all three case studies and the control simulation of §S10 are in the deposit of §S15. Random number generation used fixed seeds for reproducibility.

###### Notation in the Reduced and Three-Compartment Models

| Quantity | Reduced framework | Full cascade (eq. 3) |
|---|---|---|
| external concentration | $C_{\text{ext}}$, constant or time-varying | $C_{\text{ext}}(t)$ |
| internal parent compound | $C_b$ | $C_i$ ($\equiv C_b$) |
| toxicokinetic rates | $k_u$ (uptake), $k_e$ (elimination) | $k_{\text{in}}$, $k_{\text{out}}$ |
| damage input | $k_a C_b$ (lumped) | $k_3 A$, reached through $k_2$, $k_{2,\text{out}}$ |
| damage repair | $k_r$ — shared | $k_r$ — shared |
| hazard and tolerance | $b_w, z$ (Class A) · $k, \theta$ (Class B) · $\beta$, $h_b$ — shared across frameworks | as above |

The abstract stage rates $k_1, k_2, \ldots$ of §S4.1 belong to neither column; they index the generic activation chain of the $t^n$ derivation.

##### S12.2 Parameter Estimation

Parameters were estimated by maximum likelihood. For a cohort of $n_0$ organisms observed at times $t_0 < t_1 < \cdots < t_T$ with $a_j$ survivors at $t_j$, the conditional binomial log-likelihood is:[19]

$$\ell = \sum_{j=0}^{T-1} \left[ a_{j+1} \ln q_j + \left(a_j - a_{j+1}\right) \ln\left(1 - q_j\right) \right]$$

where $q_j = S\left(t_{j+1}\right)/S\left(t_j\right)$ is the conditional survival probability from $t_j$ to $t_{j+1}$. Background mortality was fixed at the control mortality of each experiment (Table M2) rather than estimated jointly with the toxicity parameters; the consequences of that choice for parameter identifiability are discussed by Plantade *et al.*[S13]

The NLL was minimised using the Nelder–Mead simplex algorithm[19] with 6–30 random starting points drawn log-uniformly across the feasible parameter range (Table M1).

**Table M1. Parameter bounds for multi-start optimisation.** Bounds are enforced by the optimiser on the $\log_{10}$ scale; rates in $d^{-1}$.

| Parameter | Symbol | Range (log-uniform) | Shared by |
|---|---|---|---|
| Damage rate, RED (scaled damage: formation and repair share one rate) | $k_d$ | dimethoate $10^{-1.5}$–$10^{0.5}$; diazinon $10^{-4}$–$10^{0}$ | RED models |
| Metabolic activation rate | $k_2$ | dimethoate $10^{-1}$–$10^{1.5}$; diazinon fixed at 0.896 (Table M2) | 3-comp models |
| AChE inhibition rate | $k_3$ | $10^{-2}$–$10^{0}$ | 3-comp models |
| Damage (AChE-P) recovery rate | $k_r$ | $10^{-5}$–$10^{-2}$ ($t_{95} \geq 300$ d); $t_{95} \geq 100$ d, $\geq 30$ d or up to $10^{1.5}$ in the sensitivity of §S9.2 | 3-comp models |
| Metabolite clearance rate | $k_{2,\text{out}}$ | dimethoate 3.0 (fixed) or $10^{-1.3}$–$10^{1.5}$ (estimated); diazinon fixed at 3.278 (Table M2) | 3-comp models |
| Killing rate (Class B) | $k$ | $10^{-8}$–$10^{-1}$ | Class B models |
| Killing rate (Class A) | $b_w$ | $10^{-8}$–$10^{-1}$ | Class A models |
| Median threshold | $\theta_{\text{med}}$ or $z_{\text{med}}$ | $10^{-2}$–$10^{4}$ | All models |
| Shape parameter | $\beta$ | 5.0 (fixed) | All models |

The dimethoate reduced-model fits of both structures end at the lower bound of $k_d$, $10^{-1.5}$ $d^{-1}$ (Table S17). Widening that bound to $10^{-4}$ returns them to the new lower bound, along a ridge with the threshold, and changes the calibration and prediction differences from −23.3 and −125.1 to −22.0 and −100.0, the same sign and a similar size.

**Table M2. Fixed toxicokinetic parameters and background mortality.** Dimethoate has no measured toxicokinetics; its three-compartment model is run with the measured *G. pulex* constants of imidacloprid and of aldicarb as the two ends of a range (§S8.1). Diazinon uses the constants measured in the study that produced the survival data, with parent loss the sum of elimination (8.464 $d^{-1}$) and activation ($k_2$).

| Parameter | Dimethoate (imidacloprid / aldicarb constants) | Diazinon | CPF | Carbaryl | Source |
|---|---|---|---|---|---|
| $k_{\text{in}}$ (L $kg^{-1}$ $d^{-1}$) | 1.96 / 10.80 | 118.9 | **747** | **23.4** | dimethoate ref S15 (Table 1); diazinon ref 20 (Table 1); CPF and carbaryl ref 23 |
| $k_{\text{out}}$ ($d^{-1}$) | 0.267 / 6.62 | 9.36 | **0.45** | 0.27 | as above |
| $k_2$ ($d^{-1}$) | estimated | 0.896 | — | — | diazinon ref 20 |
| $k_{2,\text{out}}$ ($d^{-1}$) | 3.0 (or estimated) | 3.278 | — | — | dimethoate assumed, §S8.1; diazinon ref 20 |
| $h_b$ ($d^{-1}$) | 0.0175 | 0.0245 | 0.0071 | 0.0071 | dimethoate and diazinon: constant-hazard fit to the controls of refs 3 and 20; CPF and carbaryl from ref 23 |

The chlorpyrifos and carbaryl columns are the values used by the Threshold Damage Model of ref 23 (its Table 1) and are the ones behind Figure S10; main text §4.4 uses the two-compartment description of that model rather than the three-compartment one, so no metabolite clearance constant enters.

##### S12.3 Comparison Metrics, the Carry-Over Interval and Table Conventions

$\Delta$NLL = $\text{NLL}_{\text{SD}} - \text{NLL}_B$, so a positive value favours the peak-based structure and a negative one the cumulative structure. The fourteen-compound table of §S7 uses the same convention, with the reduced-framework GUTS-IT fits standing in for the peak-based structure (§S3.1). The gap closure (GCR) is the fraction of the peak-based structure's reduced-framework calibration deficit that restoring the compartments recovers:

$$\text{GCR} = \frac{\text{NLL}_B^{\text{RED}} - \text{NLL}_B^{\text{FULL}}}{\text{NLL}_B^{\text{RED}} - \text{NLL}_{\text{SD}}^{\text{RED}}}$$

For example, dimethoate with the aldicarb constants: GCR = (164.18 − 148.40)/(164.18 − 140.92) = 15.78/23.26 = 68%; with the imidacloprid constants (164.18 − 136.78)/23.26 = 118% (Table S5). A value above 100% means the three-compartment peak-based fit is better than the reduced cumulative one. Nash–Sutcliffe efficiency (NSE) measures how closely predicted survival follows the observed fractions, on calibration and validation data alike:

$$\text{NSE} = 1 - \frac{\sum_j \left(S_{\text{obs},j}/S_{\text{obs},0} - S_{\text{pred},j}\right)^2}{\sum_j \left(S_{\text{obs},j}/S_{\text{obs},0} - \bar{S}_{\text{obs}}\right)^2}$$

where $\bar{S}_{\text{obs}}$ is the mean of the observed fractions $S_{\text{obs},j}/S_{\text{obs},0}$. A negative efficiency is worse than predicting the observed mean itself, which is the reading used in the main text.

**Interval and multiplicity rule for the carry-over ratios.** Both survival fractions entering $\mathcal{R}$ are control-corrected over matched windows. The interval is formed on the log scale by the delta method: $var \log \mathcal{R}$ is estimated as $(1 - p_1)/n_1 + (1 - p_2)/n_2$, where $p_1$ and $p_2$ are the uncorrected within-window survival fractions of the two pulses and $n_1$, $n_2$ the numbers alive at the end of each window, and the reported bounds are $\mathcal{R} \exp\left(\pm 1.96\sqrt{var \log \mathcal{R}}\right)$. This keeps the interval strictly positive and matches the multiplicative form of the ratio; uncertainty in the control correction is not propagated: control survival over the windows is 0.82–0.98, and propagating it lowers each lower bound by at most 0.03 and leaves all seven exclusions in place. The rule is implemented in `carryover_ratio.py`. Across the fifteen informative comparisons the exclusion calls are Benjamini–Hochberg corrected (two-sided, 5%). The diazinon rows of Table S2 (`carryover_ratio_diazinon2010.py`) use the same matched, control-corrected windows but combine exact (Clopper–Pearson) binomial intervals on the log scale; all three are non-informative under the counting rule (§S6.1.1), so the difference touches no exclusion call.

**Notation in the tables.** Bold marks the values quoted in the main text or in the section that introduces the table. An em dash (—) means not applicable: a quantity the configuration does not define, or one that was not estimated. Where a table reports $\Delta$NLL, the sign convention is that of main text §3.3 unless the caption says otherwise.

##### S12.4 Data Sources, Designs and Sample Sizes

All survival data are for *Gammarus pulex*. The dimethoate set is a 4-day LC50 test over seven concentrations ($n$ = 20; ref 3, SI Table S4) and three 10-day treatments ($n$ = 80 each; ref 3, SI Tables S11 and S12): A and B, two one-day pulses one and five days apart, and C, a sustained exposure (§S11.1).[3] The three share a time-weighted average exposure of about 9,000 nmol $L^{-1}$, so exposure pattern can be compared at matched total exposure. The diazinon set is the

2010 pulse experiment of ref 20: three treatments of two one-day pulses at about 100 nmol $L^{-1}$, separated by 2, 7 and 15 days ($n$ = 70 each), with controls, and with the toxicokinetics measured in the same study. Its carry-over ratios are computed in §S3.4 and Table S2. The chlorpyrifos–carbaryl set exposes animals to compound 1 on days 0–1 and compound 2 on days 15–16 in both orders; the parameters are the published ones, so the prediction of main text §4.4 is made a priori and nothing is fitted. The fourteen-compound series supplies both the carry-over ratios of §S6.1 and the parameter comparison of §S7.

#### S13. Parameter Tables, the Diazinon Results and the Chlorpyrifos–Carbaryl Experiment

The parameters behind every case study are collected here: the fitted values for dimethoate and diazinon in Table S17 (the fixed toxicokinetic values are in Table M2), the diazinon calibration–prediction results in Tables S18 and S19 (dimethoate's are Tables S5–S7 of §S8.3), and the published chlorpyrifos–carbaryl parameters in Table S20. §S13.1 gives the fitted parameters, §S13.2 the diazinon calibration–prediction results for each choice of calibration treatment, and §S13.3 the chlorpyrifos–carbaryl experiment.

##### S13.1 Fitted Parameters of the Two Case Studies

**Table S17. Fitted parameters of the two case studies (calibration fits).** One parameter set per damage model (RED, one compartment; 3-comp, three compartments) and hazard structure (PM, peak-based; SD, cumulative), from the multi-start fits of §S12.2 within the bounds of Table M1; † marks a value at a bound or within 0.5% of it; rates in $d^{-1}$, killing rates per damage unit per day, thresholds in damage units. For RED the $k_r$ column holds the single rate $k_d$ of the scaled-damage model. Dimethoate is calibrated on the 4-day LC50 series, with the aldicarb constants for the three-compartment rows; diazinon on treatment A (2-day interval), with $k_2$ and $k_{2,\mathrm{out}}$ fixed at the measured values (Table M2). Every three-compartment fit has $k_r$ at a bound, the peak-based ones at the slow end and the cumulative ones at the 300-d recovery bound; the diazinon cumulative fit reaches a bound in three of its four free parameters. NLL is the calibration negative log-likelihood and agrees with Tables S5 and S18 to within multi-start variation (≤ 0.1). Each set is one point on the likelihood ridge of §S9.1: the combination of $k_2 k_3$, the threshold and the killing rate is identified, the individual values are not, so a refit can return a different point with the same likelihood (Tables S8, S8b). `case_fitted_parameters.py`.

| Compound, damage model | Structure | $k_2$ | $k_3$ | $k_{2,\mathrm{out}}$ | $k_r$ | $k$ or $b_w$ | $\theta_{\mathrm{med}}$ or $z_{\mathrm{med}}$ | NLL |
|---|---|---|---|---|---|---|---|---|
| Dimethoate, RED | PM | — | — | — | 0.0316† | $2.9\times10^{-4}$ | 633 | 164.2 |
| Dimethoate, RED | SD | — | — | — | 0.0316† | $2.8\times10^{-4}$ | 170 | 140.9 |
| Dimethoate, 3-comp, $k_{2,\mathrm{out}}$ fixed | PM | 1.08 | 0.999† | 3 | $1\times10^{-5}$† | $2.1\times10^{-5}$ | 6677 | 148.4 |
| Dimethoate, 3-comp, $k_{2,\mathrm{out}}$ fixed | SD | 0.368 | 0.0523 | 3 | $1\times10^{-2}$† | $1.4\times10^{-3}$ | 47.7 | 137.8 |
| Dimethoate, 3-comp, $k_{2,\mathrm{out}}$ estimated | PM | 0.298 | 0.0433 | 0.0501† | $1\times10^{-5}$† | $4.8\times10^{-4}$ | 192 | 137.2 |
| Dimethoate, 3-comp, $k_{2,\mathrm{out}}$ estimated | SD | 4.28 | 0.0968 | 3.96 | $1\times10^{-2}$† | $7.7\times10^{-5}$ | 817 | 137.8 |
| Diazinon, 3-comp | PM | 0.896 | 0.129 | 3.278 | $1\times10^{-5}$† | 0.0127 | 33.4 | 195.8 |
| Diazinon, 3-comp | SD | 0.896 | 1† | 3.278 | $1\times10^{-2}$† | $1.8\times10^{-4}$ | 0.01† | 200.6 |

##### S13.2 Diazinon: Calibration, Prediction and the Choice of Calibration Treatment

The diazinon comparison of main text §4.4 is set out in Table S18, for the reduced framework and for three compartments with the toxicokinetics measured in the same study.[20]

**Table S18. Diazinon: calibration (treatment A) and prediction (treatments B and C).** Entries are NLL; positive ΔNLL favours the peak-based structure; prediction NLL is summed over treatments B and C. The three-compartment row uses the toxicokinetics measured in the same study (Table M2; `case_diazinon.py`); the reduced-framework row is fitted by `diazinon_reduced_framework.py`.

| Model | Calibration PM | Calibration SD | $\Delta NLL_{cal}$ | Prediction PM | Prediction SD | $\Delta NLL_{pred}$ |
|---|---|---|---|---|---|---|
| RED | 189.2 | 192.5 | +3.3 | 436.6 | 396.4 | −40.2 |
| **3-comp** | **195.8** | **200.6** | +4.8 | **403.3** | **407.9** | +4.6 |

Nash–Sutcliffe efficiency for the three-compartment fits: peak-based +0.78 (A), +0.91 (B), +0.88 (C); cumulative +0.89, +0.96, +0.19. All values from the exact matrix-exponential integrator (§S12.1), with the recovery rate bounded at $t_{95} \geq 300$ d; without the bound the three-compartment differences are −0.3 and −3.6 (§S9.2).

Calibrating on one treatment and predicting the other two can be done three ways, and Table S19 reports all three.

**Table S19. Diazinon prediction for each choice of calibration treatment.** Calibration on one treatment, prediction of the other two. Positive values favour the peak-based structure. Background mortality is estimated from this experiment's own controls ($h_b = 0.0245$ d$^{-1}$); toxicokinetics and bounds are those of Table S18. The worst-case column is the lowest Nash–Sutcliffe efficiency over the three treatments (`diazinon_reduced_framework.py`).

| Calibration | RED, prediction ΔNLL | RED, worst NSE PM / SD | 3-comp, prediction ΔNLL | 3-comp, worst NSE PM / SD |
|---|---|---|---|---|
| A (2-day interval) | −40.2 | +0.34 / +0.15 | +4.6 | +0.88 / +0.19 |
| B (7-day interval) | −7.5 | +0.11 / +0.91 | +22.0 | +0.56 / +0.36 |
| C (15-day interval) | −50.5 | −0.84 / +0.93 | +7.9 | +0.10 / +0.81 |

In the reduced framework the cumulative structure predicts better for every calibration choice, by 7.5 to 50.5 units. With the measured toxicokinetics, three compartments and the recovery bound the prediction differences are +4.6, +22.0 and +7.9, all towards the peak-based structure; without the bound they are −3.6, +9.5 and −2.1 and change sign with the calibration treatment. The worst-case efficiencies favour the peak-based structure when calibrated on A or B and the cumulative one when calibrated on C. Restoring the compartments therefore removes the reduced-framework advantage of the cumulative structure, but which structure then leads depends on the assumed recovery bound (§S9.2) and the tolerance shape (§S9.3), so neither advantage is established (main text §5.3). All three intervals are shorter than diazinon's recovery time (§S3.4), so the experiment does not reach the phase in which the structures diverge.

The 2010 experiment is also examined in the companion analysis,[7] for a different quantity: whether survival is flat across the window in which the running maximum is frozen. That is a property of the damage model rather than of the hazard structure.

##### S13.3 Chlorpyrifos–Carbaryl: Recovery Fixes the Direction of the Asymmetry, Not Its Size

Chlorpyrifos inhibits acetylcholinesterase irreversibly after oxidative activation; carbaryl carbamylates it reversibly. Ashauer *et al.* gave *Gammarus pulex* a one-day pulse of one compound, fourteen days of clean water, then a one-day pulse of the other, in both orders.[23] Carbaryl killed 31% of unexposed animals and 55% of animals given chlorpyrifos a fortnight earlier ($p = 0.001$), whereas chlorpyrifos killed 12% and 21% in the mirrored comparison ($p = 0.13$). The same compound at the same dose killed far more animals when it followed chlorpyrifos. The interaction itself is not significant (log odds ratio difference +0.33, 95% CI −0.72 to +1.38, from the counts 31/100 and 46/83 for carbaryl and 12/100 and 13/61 for chlorpyrifos; `cpf_carbaryl_per_compound.py`), so the asymmetry is not established by the counts alone; what the case shows is that its direction follows from the damage trajectory.

The published recovery rates are the whole mechanism the model needs: 0.97 $d^{-1}$ for carbaryl against 0.169 for chlorpyrifos (Table S20), a contrast that follows from the chemistry of adduct reversal (§S7.2). A fortnight nearly clears a carbamate lesion and not a phosphorylation one, so only the chlorpyrifos-first order arrives at the second pulse with much of its damage left.

Using the published parameters without fitting (Table S20), both structures reproduce the direction of the asymmetry (Figure S10): chlorpyrifos leaves 29% of its peak damage at day 14 against 6% for carbaryl, and both predict the absence of that carry-over in the reverse order. Neither reproduces the size of the asymmetry. The quantity the original study reports and predicts is the mortality within each pulse window among the animals alive at its start (ref 23, p. 5539). With each killing rate set to match the unexposed carbaryl pulse — the only quantity taken from these counts — carbaryl mortality rises from 35.0% given first to 38.6% given second under the cumulative structure (+3.6 percentage points) and to 36.4% under the peak-based structure (+1.4), against 31% to 55% observed (+24) and 33% to 47% predicted by the Threshold Damage Model published with the experiment (+14). For chlorpyrifos both structures predict a fall when it follows carbaryl (−4.6 and −3.9 points), against a rise of 9 observed and 1 predicted there (`cpf_carbaryl_per_compound.py`). The re-implementation used here, with one pooled damage state and constant pulse concentrations, therefore does not reproduce the published model's prediction, and the comparison below is between the two hazard structures under that simplified model, not with ref 23. The shortfall exceeds the gap between the two structures, so the experiment does not choose.

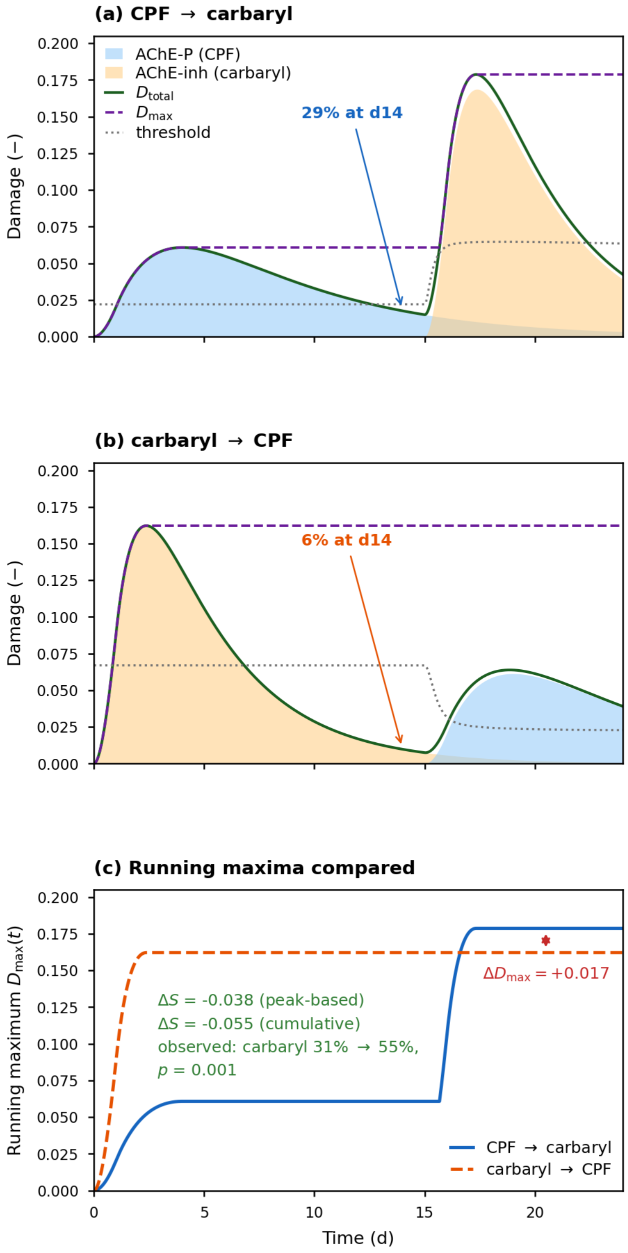


**Figure S10. Chlorpyrifos–carbaryl directional asymmetry.** Parameters and exposures are those published with the experiment;[23] only the killing rate of each structure is set, to match mortality in the carbaryl pulse without prior exposure. (a) Chlorpyrifos followed fourteen days later by carbaryl and (b) the reverse order: shading separates each compound's contribution, the solid line is their sum, the dashed line its running maximum, the dotted line the damage-weighted mixture threshold. (c) The running maxima compared, with day-24 survival differences for both structures; the peak-based killing rate is set so that they agree on the carbaryl pulse without prior exposure. The day-24 differences are differences in total survival, not the per-window mortality compared with ref 23 in the text.

The direction, unlike the size, is fixed by the damage trajectory alone. Under the peak-based structure survival depends on the trajectory only through $D_{\max}$, which is larger in the chlorpyrifos-first order — 0.179 against 0.162 — so, for a threshold that does not depend on the order of the compounds, no choice of threshold, killing rate or tolerance spread can reverse its prediction; the cumulative structure, whose damage integral is also larger in that order, predicts the same direction. The Threshold Damage Model, which first predicted the asymmetry, is cumulative.[23] Under the simplified model used here the cumulative structure comes marginally closer to the observed carbaryl increase than the peak-based one (3.6 against 1.4 percentage points, against 24 observed). Discrimination therefore rests on the carry-over ratio (main text §4.1).

**Table S20. Chlorpyrifos–carbaryl: published toxicokinetic and toxicodynamic parameters used for the a priori predictions.**

| Parameter | Chlorpyrifos | Carbaryl | Source |
|---|---|---|---|
| $k_{\text{in}}$ (L kg$^{-1}$ d$^{-1}$) | **747** | **23.4** | Ashauer *et al.* (2007)[23] Table 1 |
| $k_{\text{out}}$ (d$^{-1}$) | **0.45** | 0.27 | as above (chlorpyrifos from ref S18) |
| BCF = $k_{\text{in}}/k_{\text{out}}$ (L kg$^{-1}$) | 1660 | 87 | Calculated |
| Killing rate $k_k$ (g µg$^{-1}$ d$^{-1}$) | 0.134 | 0.42 | Ashauer *et al.* (2007)[23] Table 1 |
| Damage threshold (–) | 0.022 | 0.067 | as above |
| $k_r^{\text{TD}}$ (d$^{-1}$), TDM | **0.169** | 0.97 | as above |
| Background hazard $h_b$ (d$^{-1}$) | 0.0071 | 0.0071 | as above |
| $k_{r,\text{SD}}$ (d$^{-1}$), GUTS | 0.630 | 0.122 | Ashauer *et al.* (2015)[2], stochastic-death fit |
| $k_{r,\text{IT}}$ (d$^{-1}$), GUTS | 0.0001 | 0.0168 | Ashauer *et al.* (2015)[2], individual-tolerance fit |
| AChE mechanism | Irreversible (phosphorylation) | Reversible (carbamylation) | — |
| Time for damage to fall below 5% of peak, toxicokinetic loss included (d) | 25 | **15** | Ashauer *et al.* (2007)[23] |

Every row above is the value that the Threshold Damage Model of Ashauer *et al.* (2007)[23] actually used, taken from Table 1 of that paper; the chlorpyrifos uptake and elimination constants are attributed there to the direct measurements of Ashauer *et al.* (2006)[S18]. The tabulated values reproduce the internal concentrations of that paper's Figure 1b — 543 and 299 ng g$^{-1}$ computed against 580 and 290 plotted — and its peak mixture damage (0.162 and 0.179 against 0.17 and 0.19). A separate biotransformation experiment[21] measured parent-specific chlorpyrifos elimination and returns a faster constant, near 2 d$^{-1}$; that is a different quantity, since parent-specific loss includes biotransformation, and it is not what the 2007 model used. The recovery time in the last row is the interval the same paper reports for damage to fall below 5% of its maximum.

Each recovery rate belongs to the model it was fitted under, and values from different models cannot be substituted for one another. The 2007 Threshold Damage Model recovery rates ($k_r^{\text{TD}}$ in Table S20) are those used for the a priori prediction of main text §4.4; the TDM places the hazard rate proportional to damage above a threshold and is therefore a cumulative-hazard (Class A) formulation. The GUTS re-analysis of the same two compounds in 2015[2] returns different values under each hazard structure, and the ordering between the compounds is not preserved: under GUTS-SD carbaryl recovers more slowly than chlorpyrifos, under GUTS-IT both rates are below 0.02 d$^{-1}$ (0.0001 and 0.0168).

**Independent refit of the recovery rates.** The Threshold Damage Model was refitted to the original laboratory data for both compounds, with toxicokinetics fixed and background hazard identified from the controls. Chlorpyrifos returns $k_r$ = 0.039 d$^{-1}$ (profile-likelihood interval 0.02–0.12), placing the published 0.169 outside it; carbaryl returns $k_r$ = 4.8 d$^{-1}$ (interval 2.5–8.0), which excludes the published 0.97 by ΔNLL = 7.9. Both departures widen the recovery contrast between the two compounds, from about six-fold (0.97 against 0.169) to over a hundred-fold (4.8 against 0.039). The chlorpyrifos rate is not identifiable at all in the single-compound chlorpyrifos experiment, one of the two laboratory datasets refitted, whose profile is flat across two orders of magnitude — an instance of the observation-window condition of §S3.4, since its five-day recovery intervals are short relative to the 95% recovery time $\ln 20/k_r$.

#### S14. Designing a Decisive Experiment: Three Conditions and What Each Outcome Settles

The two structural tests — the carry-over ratio (§S6.1) and death during a clean-water phase (§S3.2) — make the same demands of an experiment, and the demands are easiest to state as three conditions. The first is a precondition for any comparison at all; the second belongs to the ratio test and the third to the clean-water test, and Table S21 sets out which outcomes then decide something and which decide nothing. Each follows from a result above; the first two scale with the compound's damage recovery time and the third with how its lesion is formed.

##### S14.1 Three Conditions on the Design

**Condition 1 — the observation window must reach the post-equilibrium phase.** While damage is rising the running maximum equals the current value and the two hazards are the same function of the same trajectory (§S3.4), so the observation after each pulse must outlast both the elimination and the damage recovery, $\max(\ln 20/k_e,\ \ln 20/k_r)$, with $k_e$ the elimination rate and $k_r$ the damage recovery rate. Without it no observation of either kind can discriminate, whatever else the design does. The condition is about detectability rather than about the size of any difference. In a shorter window the two structures do not differ in what they predict, so no amount of replication recovers the distinction; above it they diverge, and the design question becomes how large the divergence is at the exposure being run. The companion paper uses the same distinction for the clean-water window.[7]

**Condition 2 — the interval between two identical pulses must exceed the compound's own recovery time.** The theorem of §S6.1 assumes damage resets between pulses. Below that interval the peak-based structure's ratio also falls below unity, so it becomes indistinguishable from the cumulative structure, which is at or below unity in any case (§S6.1). The 2010 diazinon pulse experiments (§S3.4, Table S2) fail this condition at all three of their intervals, and their ratios duly fall below unity. That the interval has to be compound-specific is not particular to the archive used here. Zhao and Newman ran two identical 12-hour pulses on *Hyalella azteca* at four recovery intervals and estimated the interval needed for full recovery at 83 hours for dissolved copper against 18 for pentachlorophenol, a fivefold difference between two compounds in one species.[S8] The condition is necessary but not sufficient. Naddy *et al.* gave *Daphnia magna* two 12-hour chlorpyrifos pulses at intervals of 3, 7 and 14 days and found survival at or below 15% at every interval, while the same total exposure delivered as two 6-hour pulses left every animal alive at 3 and 14 days.[S4, S5] Where the first pulse alone drives damage past the lethal level, no interval helps.

**Condition 3 — survival must be observed through a phase in which the input driving damage has itself reached zero.** This is the condition under which the falsification test of §S3.2 applies, and it is not the same as clearance of the parent compound. For a compound that binds irreversibly, the input to the damage state does not reach zero even when the tissue burden has fallen.

##### S14.2 What Each Outcome Would Settle

**Table S21. What each outcome would settle.**

| Observation | What it settles | Ground |
|---|---|---|
| $\mathcal{R} > 1$ | excludes **GUTS-SD**, for that compound, at every parameter value; no interval condition is needed | Proposition S4, §S6.1 |
| Death during a clean-water phase meeting Condition 3 | excludes **Class B**, for that compound | Proposition S2, §S3.2 |
| $\mathcal{R}$ at the ceiling $1/S_1$, with damage reset | consistent with Class B, which attains it exactly | Proposition S5, §S6.2 |

| Observation | What it settles | Ground |
|---|---|---|
| $\mathcal{R} \le 1$ | nothing: before damage resets both structures predict it, and after reset the cumulative structure does (§S6.1.2); Condition 2 decides only whether the comparison enters the count of §S6.1.4 | §S6.1 |
| A difference in calibration goodness of fit | nothing on its own; the likelihood points the right way more often than not, and rarely by enough to decide | §S10, main text §5.1 |
| A sign reversal under frozen-parameter extrapolation | excludes neither structure; it is evidence about the damage model both share, of a different kind from a calibration likelihood | main text §4.4 |

Only the first two rows exclude a structure, and neither is a likelihood comparison; that is why this paper argues the classification on structural rather than parametric grounds.

##### S14.3 Minimum Designs by Mode of Action

What the three conditions require of a real test depends on the compound, because two of them scale with its recovery time (Table S22).

**Table S22. Minimum two-pulse design by mode of action.** Recovery times are those the counting rule of §S6.1.1 uses: $t_{95} = \ln 20/k_r$ under the individual-tolerance fits of Table S3. Both minima are one $t_{95}$: Condition 1 sets the window that must follow each pulse, Condition 2 the interval between them, so a design runs for about two $t_{95}$ (`design_feasibility.py` lists each compound). A window of three $t_{95}$ leaves room for the divergence to be measured but is not required for detectability. † $k_r$ at the lower fitting bound, so $t_{95}$ is a lower limit and no finite design is implied.

| Mode of action | Compounds | $t_{95}$ (d) | Minimum design (d) | Reachable from survival alone |
|---|---|---|---|---|
| Baseline narcotic | 3 | 0.003–0.4 | 1.0–1.4 | yes |
| Uncoupler | 3 | 2.7–258 | 6 (4,6-dinitro-*o*-cresol) to 517 (2,4,5-trichlorophenol) | for the fastest only |
| Carbamate | 3 | 0.8–178 | 2 (carbofuran) to 357 (carbaryl) | for two of three |
| Organophosphate | 3 | 36 to ≥ 30,000† | 72 (malathion), 335 (diazinon), none (chlorpyrifos†) | no, in practice |
| Reactive | 2 | ≥ 30,000† | none | no |
| Neonicotinoid | — | — | — | Condition 3 unreachable |

These recovery times are fitted to survival, and for one class they can be set against estimates from the lesion itself. In *Daphnia magna* the model fitted to the activity of acetylcholinesterase inhibited by diazoxon places 99% recovery at about sixteen days, whereas the enzyme carbamylated by carbaryl recovers 95% in nine hours.[S1, S19] The mechanism is resolved in vitro: carbamylated enzyme decarbamylates at 1.9 $h^{-1}$, while the phosphorylated enzyme reactivates at a rate indistinguishable from zero.[S19] The measured lesions separate the two classes by more than an order of magnitude; the survival fits of Table S3 do not, for carbaryl and diazinon have similar fitted recovery times under each structure (178 and 167 d individual-tolerance, 25 and 20 d stochastic-death). Survival-inferred recovery times are therefore the weak link of every design in Table S22, and for the anticholinesterases a measured enzyme trajectory is the better input (§S14.4). The carbaryl figure is fitted to enzyme activity alone and the in vitro reactivation rate to neither organism nor model; the diazinon figure is a model estimate from a simultaneous fit that included survival under a cumulative hazard, so it is the less independent of the two. Both source papers adopt the cumulative hazard for the organism-level step, and nothing in this subsection depends on that choice.

The last row of Table S22 is not a long duration but an impossibility of a different kind. Receptor-bound thiacloprid, a neonicotinoid, does not detectably leave the tissue of *G. pulex* on the experimental time scale so far examined — eight days in the only study to resolve the bound fraction chemically[30] — so the input to the damage state does not reach zero within any test of ordinary length. Condition 3 therefore cannot be satisfied by lengthening the experiment. Whether it can be satisfied at all for this class is an open question.

##### S14.4 Where the Design Is Impractical: Long Recovery and Irreversible Binding

Table S22 shows two kinds of obstacle, and the ways around them differ accordingly.

Where the survival-inferred recovery time is long, the design is long, and its cost is set by background mortality. For malathion, the fastest-recovering organophosphate in the archive, the minimum design runs about 72 days; at the background mortality rates of Table M2 the fraction of control animals still alive at the end is 17% at diazinon's rate, 60% at chlorpyrifos' and 28% at dimethoate's. For diazinon it runs about 335 days and leaves 0.03%, 9% and 0.3%; for chlorpyrifos and the reactive compounds the fitted recovery is at its bound and no finite design follows. Pentachlorophenol (about 89 days) lies between, and 2,4,5-trichlorophenol and carbaryl (over 350 days) with diazinon. Power follows the interval rule of §S12.3: with a first pulse leaving three in five alive, the smallest ratio whose interval clears unity is 1.43 at forty animals per pulse, 1.34 at sixty and 1.25 at a hundred (`design_feasibility.py`). Sixty animals per pulse therefore resolve ratios a little above the median of the informative ratios in the archive (1.29, §S6.1.4), but only where enough controls survive for the correction to mean something; for most organophosphates a two-pulse test designed from survival-inferred recovery is not practical. How rarely published designs meet such conditions is counted in the companion paper, which finds only a small minority of pulsed treatments informative for the clean-water observation it tests.[7]

Two routes shorten the experiment instead of running its full length. The recovery time is set by the chemistry of the lesion — resynthesis of the enzyme that the phosphorylated one has displaced — so raising temperature or moving to a species with faster enzyme turnover shortens $t_{95}$, at the price of a verdict that holds for those conditions. The better route is to stop inferring the damage state and measure it: the anticholinesterases are the one class whose lesion is directly assayable, and the enzyme recovery times quoted above are derived from such measurements. Taken at face value in another species, the diazoxon estimate implies $t_{95} \approx 10$ days and a design of about three weeks, against about 335 days from the survival fit. With the damage trajectory observed rather than modelled, the falsification test of §S3.2 applies directly and no two-pulse ratio is needed.

For the neonicotinoids the obstacle is not cost. Condition 3 asks for a phase in which the input driving damage has reached zero, and a compound that binds its receptor irreversibly never provides one; saturation of the receptor pool then removes the concentration axis as well (main text §5.4). One route survives: a ratio above unity needs no interval condition (Table S21), so a two-pulse experiment can still exclude the cumulative structure if it returns one — though irreversible binding makes that outcome unlikely rather than impossible. Otherwise the choices are to work below the saturating range and accumulate damage through duration instead of concentration, accepting the loss of power; to measure receptor occupancy as the anticholinesterase work measures enzyme activity; or to leave the class to the clean-water analysis of the companion paper,[7] which is why this paper draws no conclusion for it.

#### S15. Code and Data Availability

***What the deposit holds.*** The code that reproduces the numbers, tables and figures of this paper is deposited at Zenodo (https://doi.org/10.5281/zenodo.23488393) under an MIT licence for the code and CC BY 4.0 for the figures. Version 0.9, deposited at submission, holds the Python scripts of the table below and the derived tables they write as CSV; version 1.0, on acceptance, adds the 14 figures as published. The three published data files the analysis reads, spreadsheets from the supplements of refs 2, 20 and 21, are not redistributed; the deposit README names each file and its source. The code requires only numpy, scipy, pandas, matplotlib and an Excel reader; most scripts finish in minutes, and the longest, `tolerance_shape_sensitivity.py`, takes about an hour on one core and can be run one tolerance shape at a time (`--dz`, `--dm`). Scripts locate the data and figure directories relative to their own file, so no paths need editing. Each script is named for what it computes, and the first line of its docstring states the table, figure or section it produces.

***Which figures regenerate.*** Every figure regenerates directly from the deposited code: main-text Figures 1–4 (`figure1_carryover_ratio.py`, `figure2_conservatism_ordering.py`, `figure3_control_simulation.py` and `figure4_recovery_rates.py`) and Figures S1–S10 of this document (`figureS1_four_cells.py`, `figureS2_frailty_envelope.py`, `figureS3_carryover_vs_repair.py`, `figureS4_recovery_times.py`, `figureS5_damage_curvature.py`, `figureS6_metabolite_clearance_profile.py`, `case_dimethoate.py`, `figureS8_reduced_structures.py`, `figureS9_case_studies.py`, `case_chlorpyrifos_carbaryl.` `run_all_figures.py` prints this inventory and checks it against the files present. Every table regenerates from the deposited code; Tables S3, S20 and M2 quote published values. The recovery-rate refit of §S13.3 needs two 2007 laboratory data files that are not redistributed here.

***Which script produces what.*** The scripts that produce the quantitative claims of this Supporting Information are listed below in the order of the sections they serve.

| SI section | Script | Produces |
|---|---|---|
| — | `tktd_core.py` | Shared numerics: damage propagation, survival under both hazard structures, likelihood, multi-start fitting |
| §S3.2 | `clean_water_test_diazoxon.py` | The clean-water diazinon comparison |
| §S3.3 | `conservatism_ordering_table.py` | Which structure is the more sensitive, early and late (Table S1) |
| §S3.3 | `conservatism_ordering_scan.py`, `figure2_conservatism_ordering.py` | The same comparison over the parameter space (Table S1b, main text Figure 2) |
| §S3.4, §S6.1.4 | `carryover_ratio_diazinon2010.py` | The three 2010 diazinon pulse ratios (Table S2) |
| §S1 | `figureS1_four_cells.py` | Damage and survival in the four cells (Figure S1) |
| §S5 | `michaelis_menten_repair.py` | Saturating repair against the dimethoate calibration |
| §S6.1.4 | `carryover_ratio.py`, `figure1_carryover_ratio.py` | The carry-over ratio from raw survival counts (Table S2, main text Figure 1); `--sd` gives the stochastic-death variant |
| §S6.1.2 | `figureS3_carryover_vs_repair.py` | The carry-over ratio against the recovery rate (Figure S3) |
| §S6.1.3 | `frailty_variance_bound.py` | Profile limits on the frailty variance and the frailty-limited ratio per treatment |
| §S6.1.3 | `frailty_curve_checks.py` | Overdispersion, time-curvature and saturation checks |
| §S6.1.3 | `figureS2_frailty_envelope.py` | Figure S2 |
| §S7, §S7.1 | `figure4_recovery_rates.py`, `figureS4_recovery_times.py` | Fitted recovery rates and recovery times of the fourteen compounds (main text Figure 4, Figure S4) |
| §S7.2 | `sd_recovery_bound.py` | Proposition S6: numerical check of the bound and the per-treatment bounds on the cumulative-structure recovery rate |
| §S6.2.2 | `carryover_ceiling_approach.py` | Approach to the arithmetic ceiling |
| §S6.3 | `lethal_residue_scan.py` | The lethal-residue scan over $(b_w, z)$ |
| §S8.1 | `metabolite_clearance_fit.py` | Elimination rates of activated metabolites in the same species |
| §S8.2 | `figureS5_damage_curvature.py` | Damage under constant exposure and after one pulse (Figure S5) |
| §S8.2, §S8.3 | `damage_curvature_and_step_size.py` | Power-law exponents of the damage trajectory (Table S4) and the integration step-size check |
| §S8.3 | `figureS6_metabolite_clearance_profile.py` | Profile over the metabolite clearance rate (Figure S6) |
| §S8.3, §S11 | `case_dimethoate.py` | Dimethoate fits and gap closure (main text §4.4, Tables S5–S7), calibration strategies (Table S16, Figure S7) |
| §S9.1 | `recovery_rate_free_refit.py` | The recovery rate freed from the tie (Table S8) |
| §S9.1 | `threshold_scale_check.py` | What a fitted threshold determines, in damage and exposure units (Table S8b) |

| SI section | Script | Produces |
|---|---|---|
| §S9.2 | `recovery_rate_bound_sensitivity.py` | The case studies under four bounds on the recovery rate (Tables S9, S10) |
| §S8.3, §S9.2, §S13.2, Table M1 | `no_bound_sensitivity.py` | The fits with the recovery bound removed (Table S6 last column, Table S19 text) and with the $k_d$ bound widened |
| §S9.3 | `tolerance_shape_sensitivity.py` | The case studies with the tolerance shape fixed, varied and estimated (Tables S11, S12) |
| §S9.4 | `killing_rate_shape_redundancy.py` | The reduced peak-based model against two-parameter GUTS-RED-IT |
| §S10.1 | `control_simulation.py`, `control_simulation_both_directions.py` | Control simulation, both directions (Table S13) |
| §S10.1 | `figure3_control_simulation.py` | Main text Figure 3 from the bidirectional control simulation |
| §S10.2 | `margin_intervals.py`, `margin_bootstrap_refit.py` | Percentile intervals for the margins (Table S14), counts redrawn and refitted |
| §S11.3 | `figureS8_reduced_structures.py` | The three reduced structures side by side (Figure S8) |
| §S11.3 | `figureS9_case_studies.py` | The two case-study compounds side by side, read from Tables S7 and S18 (Figure S9) |
| §S12.1 | `quadrature_nodes_check.py` | Tolerance quadrature, 8 nodes against 16, 32 and 64 |
| §S13.1 | `case_fitted_parameters.py` | The fitted parameters of the two case studies (Table S17) |
| §S13.2 | `case_diazinon.py` | Diazinon three-compartment fits (Table S18) |
| §S13.2 | `diazinon_reduced_framework.py` | Diazinon in the reduced framework (Table S18) and each choice of calibration treatment in both frameworks (Table S19) |
| §S13.3 | `case_chlorpyrifos_carbaryl.py` | The chlorpyrifos–carbaryl prediction (Figure S10) |
| §S13.3 | `cpf_carbaryl_per_compound.py` | Per-window mortality of each compound in both orders, against ref 23 (main text §4.4) |
| §S13.3 | `tdm_recovery_refit.py` | Threshold Damage Model refit of the recovery rates |
| §S14.3, §S14.4 | `design_feasibility.py` | Minimum designs (Table S22), control survival and resolvable ratio |
| §S15 | `run_all_figures.py` | The figure inventory above, checked against the files present |

The shared module `tktd_core.py` integrates the tolerance distribution with a midpoint rule whose weights sum to unity (§S12.1).

#### SI References

Superscript numbers without the prefix S refer to the reference list of the main article; those with the prefix S are listed below.

(S1) Kretschmann, A.; Ashauer, R.; Hollender, J.; Escher, B. I. Toxicokinetic and Toxicodynamic Model for Diazinon Toxicity — Mechanistic Explanation of Differences in the Sensitivity of *Daphnia magna* and *Gammarus pulex*. *Environ. Toxicol. Chem.* **2012**, *31* (9), 2014–2022. DOI: 10.1002/etc.1905.

(S2) Brock, T. C. M.; Arena, M.; Cedergreen, N.; Charles, S.; Duquesne, S.; Ippolito, A.; Klein, M.; Reed, M.; Teodorovic, I.; van den Brink, P. J.; Focks, A. Application of General Unified Threshold Models of Survival Models for Regulatory Aquatic Pesticide Risk Assessment Illustrated with an Example for the Insecticide Chlorpyrifos. *Integr. Environ. Assess. Manag.* **2021**, *17*, 243–258. DOI: 10.1002/ieam.4327.

(S3) Kallander, D. B.; Fisher, S. W.; Lydy, M. J. Recovery Following Pulsed Exposure to Organophosphorus and Carbamate Insecticides in the Midge, *Chironomus riparius*. *Arch. Environ. Contam. Toxicol.* **1997**, *33*, 29–33. DOI: 10.1007/s002449900219.

(S4) Naddy, R. B.; Johnson, K. A.; Klaine, S. J. Response of *Daphnia magna* to Pulsed Exposures of Chlorpyrifos. *Environ. Toxicol. Chem.* **2000**, *19*, 423–431. DOI: 10.1002/etc.5620190223.

(S5) Naddy, R. B.; Klaine, S. J. Effect of Pulse Frequency and Interval on the Toxicity of

Chlorpyrifos to *Daphnia magna*. *Chemosphere* **2001**, *45*, 497–506. DOI: 10.1016/S0045-6535(01)00019-4.

(S6) Cold, A.; Forbes, V. E. Consequences of a Short Pulse of Pesticide Exposure for Survival and Reproduction of *Gammarus pulex*. *Aquat. Toxicol.* **2004**, *67*, 287–299. DOI: 10.1016/j.aquatox.2004.01.015.

(S7) Andersen, T. H.; Tjørnhøj, R.; Wollenberger, L.; Slothuus, T.; Baun, A. Acute and Chronic Effects of Pulse Exposure of *Daphnia magna* to Dimethoate and Pirimicarb. *Environ. Toxicol. Chem.* **2006**, *25*, 1187–1195. DOI: 10.1897/05-465R1.1.

(S8) Zhao, Y.; Newman, M. C. Effects of Exposure Duration and Recovery Time during Pulsed Exposures. *Environ. Toxicol. Chem.* **2006**, *25*, 1298–1304. DOI: 10.1897/05-341R.1.

(S9) Aalen, O. O.; Borgan, Ø.; Gjessing, H. K. *Survival and Event History Analysis: A Process Point of View*; Springer: New York, **2008**; Chapter 6, Unobserved heterogeneity: the odd effects of frailty. DOI: 10.1007/978-0-387-68560-1.

(S10) McCarty, L. S.; Mackay, D. Enhancing Ecotoxicological Modeling and Assessment: Body Residues and Modes of Toxic Action. *Environ. Sci. Technol.* **1993**, *27* (9), 1719–1728. DOI: 10.1021/es00015a002.

(S11) van Hoogen, G.; Opperhuizen, A. Toxicokinetics of Chlorobenzenes in Fish. *Environ. Toxicol. Chem.* **1988**, *7* (3), 213–216. DOI: 10.1002/etc.5620070305.

(S12) van Wezel, A. P.; Opperhuizen, A. Narcosis Due to Environmental Pollutants in Aquatic Organisms: Residue-Based Toxicity, Mechanisms, and Membrane Burdens. *Crit. Rev. Toxicol.* **1995**, *25* (3), 255–279. DOI: 10.3109/10408449509089890.

(S13) Plantade, J.; Baudrot, V.; Charles, S. hb or Not hb — When and Why Accounting for Background Mortality in Toxicological Survival Models Matters. *MethodsX* **2023**, *10*, 102114. DOI: 10.1016/j.mex.2023.102114.

(S14) Ashauer, R.; Thorbek, P.; Warinton, J. S.; Wheeler, J. R.; Maund, S. A Method to Predict and Understand Fish Survival under Dynamic Chemical Stress Using Standard Ecotoxicity Data. *Environ. Toxicol. Chem.* **2013**, *32*, 954–965. DOI: 10.1002/etc.2144.

(S15) Ashauer, R.; Caravatti, I.; Hintermeister, A.; Escher, B. I. Bioaccumulation Kinetics of Organic Xenobiotic Pollutants in the Freshwater Invertebrate *Gammarus pulex* Modeled with Prediction Intervals. *Environ. Toxicol. Chem.* **2010**, *29* (7), 1625–1636. DOI: 10.1002/etc.175.

(S16) Kretschmann, A.; Ashauer, R.; Preuss, T. G.; Spaak, P.; Escher, B. I.; Hollender, J. Toxicokinetic Model Describing Bioconcentration and Biotransformation of Diazinon in *Daphnia magna*. *Environ. Sci. Technol.* **2011**, *45* (12), 4995–5002. DOI: 10.1021/es104324v.

(S17) Virtanen, P.; Gommers, R.; Oliphant, T. E.; Haberland, M.; Reddy, T.; Cournapeau, D.; Burovski, E.; Peterson, P.; Weckesser, W.; Bright, J.; *et al.* SciPy 1.0: Fundamental Algorithms for Scientific Computing in Python. *Nat. Methods* **2020**, *17* (3), 261–272. DOI: 10.1038/s41592-019-0686-2.

(S18) Ashauer, R.; Boxall, A.; Brown, C. Uptake and Elimination of Chlorpyrifos and Pentachlorophenol into the Freshwater Amphipod *Gammarus pulex*. *Arch. Environ. Contam. Toxicol.* **2006**, *51* (4), 542–548. DOI: 10.1007/s00244-005-0317-z.

(S19) Jeon, J.; Kretschmann, A.; Escher, B. I.; Hollender, J. Characterization of Acetylcholinesterase Inhibition and Energy Allocation in *Daphnia magna* Exposed to Carbaryl. *Ecotoxicol. Environ. Saf.* **2013**, *98*, 28–35. DOI: 10.1016/j.ecoenv.2013.09.033.
